# Oxidative Peptide Backbone Cleavage by a HEXXH Enzyme During RiPP Biosynthesis

**DOI:** 10.1101/2025.10.30.685615

**Authors:** Yao Ouyang, Yue Yu, Lingyang Zhu, Dinh T. Nguyen, Wilfred A. van der Donk

## Abstract

Ribosomally synthesized and post-translationally modified peptides (RiPPs) rely on a diverse array of enzymes to tailor peptide backbones and side chains. In this study, we characterized enzymes from two different biosynthetic gene clusters (BGCs) from Pseudomonas strains (*pfl* and *pos*) that catalyze new transformations in RiPP biosynthesis. Two α-ketoglutarate-dependent HEXXH enzymes, PflC and PosC, perform hydroxylation of multiple consecutive glutamine residues and selectively recognize a C-terminal ARMD tetrapeptide to trigger oxidative backbone cleavage that generates an amide terminus. Mutational analysis pinpoints the first position of this motif as a critical determinant. Notably, PflC displays proteolytic activity in the absence of the leader peptide, indicating that leader peptide–enzyme interactions modulate the observed reaction selectivity. The biosynthetic gene clusters also encode a unique MNIO-nitroreductase fusion enzyme that installs a rare *Z-*dehydrophenylalanine and hydroxylates an Asp residue. Collectively, this work expands both the catalytic repertoire and structural diversity accessible through bacterial RiPP biosynthesis.

## Introduction

Ribosomally synthesized and post-translationally modified peptides (RiPPs) represent one of the most structurally and functionally diverse families of natural products.^1^ Their biosynthesis relies on genetically encoded precursor peptides that undergo extensive enzymatic tailoring, which introduces a broad spectrum of chemical modifications to generate highly diversified scaffolds that display a range of bioactivities.^2–4^ Recent years have witnessed the discovery of numerous emerging RiPP-associated enzymes that mediate unprecedented chemical transformations.^5,6^ For example, multinuclear non-heme iron oxidative enzymes (MNIOs) catalyze a wide variety of different transformations.^7–9^ These findings not only highlight the remarkable diversity of chemical logic underlying RiPP maturation, but also expand the known enzymatic repertoire by revealing new strategies for scaffold modification. Therefore, continuous mining and characterization of novel RiPP enzymes is valuable, both for understanding the biosynthetic pathways and for uncovering new paradigms in enzymatic chemistry with potential applications in biocatalysis and synthetic biology.^6^

Proteins harboring the conserved HEXXH motif constitute a large class of metalloproteases traditionally associated with peptide bond hydrolysis.^10,11^ Recent studies have revealed that not all members conform to this canonical role. Morinaka, Zhang and Nicolet and co-workers characterized a HEXXH-motif protein fused to a radical SAM domain, which contrary to its prior annotation as a metallopeptidase, functions as an α-ketoglutarate (αKG)-dependent mononuclear non-heme iron enzyme that installs a β-hydroxyl group onto aspartate and histidine residues of a RiPP precursor peptide.^12,13^ This activity depends on a preinstalled cyclophane scaffold (Figure 1A). Structural analysis revealed a fold reminiscent of the zincin protease family yet distinct from any previously characterized αKG-dependent enzymes, underscoring an unusual evolutionary trajectory. While αKG-dependent enzymes are widespread in biology,^14,15^ within RiPP biosynthesis their characterized activities remain rare and are largely confined to hydroxylation.^16–21^ Recently, the first αKG-dependent-enzyme-catalyzed crosslink formation between a Cys and Tyr was reported, expanding the repertoire of these enzymes.^22^ These findings highlight the untapped chemistries embedded in RiPP biosynthetic gene clusters.

**Figure 1.**
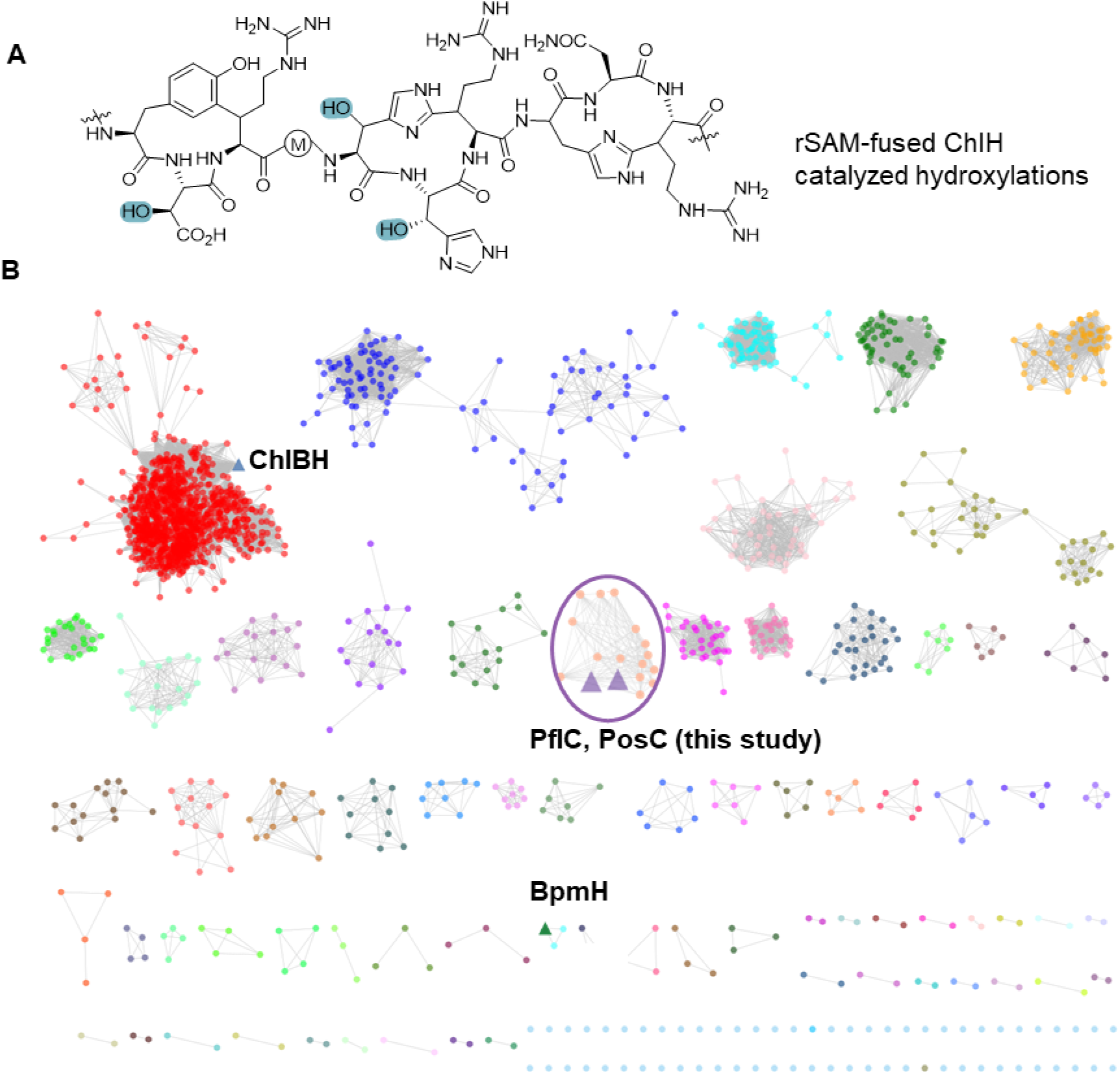
Genome mining of emerging αKG-dependent HEXXH motif enzymes. (A) ChlH catalyzes hydroxylation of His and Asp residues (highlighted in blue) in a peptide with a preinstalled cyclophane scaffold. (B) An SSN of the αKG HEXXH motif enzyme family was generated with the EFI tools using the UniProt database. RepNode 40 is shown as visualized in Cytoscape v3.10 (see Supporting Information). Reported αKG-dependent HEXXH enzymes are highlighted in blue and green triangles. PflC and PosC sequences (Table S2) were added manually because the originally reported Uniprot IDs were withdrawn during recent UniProt consolidation.

Our laboratory recently reported that the αKG-HEXXH protein BpmH is involved in the biosynthesis of biphenomycin-like macrocyclic peptides.^8^ In this study, we characterize two other novel examples of HEXXH proteins as well as an MNIO that is fused to a nitroreductase. The HEXXH enzymes unexpectedly catalyze two different transformations. They hydroxylate a string of consecutive glutamine residues in their peptide substrate and effect oxidative cleavage of the backbone after the last glutamine at a conserved ARMD motif. The MNIO-nitroreductase fusion enzyme hydroxylates an Asp residue and dehydrogenates a Phe residue to form a *Z-*dehydrophenylalanine, a transformation not seen previously in RiPPs. These new activities may provide starting points for development of useful biocatalysts.

## Results

### Genome Mining for Biosynthetic Gene Clusters Encoding HEXXH-motif Enzymes

We first generated a sequence similarity network (SSN) using the Enzyme Function Initiative Enzyme Similarity Tool (EFI-EST)^23,24^ using the αKG-dependent non-heme iron HEXXH-type β-hydroxylase domain family (IPR026337) as input with an alignment score of 30. Network analysis revealed that the largest cluster in the network belongs to rSAM-fused HEXXH enzymes, which contains the previously characterized ChlBH (Figure 1B, highlighted in a blue triangle).^12^ The hydroxylase BpmH from the biphenomycin pathway is located in a different cluster (Figure 1B, highlighted in a green triangle). A smaller cluster features enzymes encoded in BGCs that contain genes encoding a MNIO-fused nitroreductase in addition to the predicted αKG-HEXXH enzyme (Figure 1B, highlighted in a purple circle). Flavin-dependent nitroreductases have been widely found in RiPP BGCs and usually co-localize with YcaO proteins that together typically install azoles.^25^ Nitroreductases that are not associated with YcaO proteins or that are fused to MNIOs are rare and functionally uncharacterized suggesting the possibility of new chemistry. We chose a BGC from the genome of *Pseudomonas fluorescens* CH267 (*pfl,* Figure 2A). It encodes a RiPP precursor peptide (PflA), a member of the GCN-5 related acetyltransferase (GNAT) family (PflB), the HEXXH-motif protein (PflC), the MNIO-fused nitroreductase (PflD), and two transporters (PflE and PflF). A multisequence alignment of PflA with precursor peptides encoded in related BGCs shows a highly conserved C-terminal consecutive Gln motif followed by an ARMD sequence (Figures 2B and S2). No genes encoding proteases or protease domains were found nearby. Assuming that this system, like all previously characterized RiPP biosynthetic pathways,^1^ contains a sequence in the precursor peptide that is used for recognition by the biosynthetic enzymes (typically called a leader or a follower peptide), the proteolytic machinery required to remove this sequence is likely encoded elsewhere in the genome, as observed for a subset of RiPPs.^26^ Bioinformatic analysis suggests that the N-terminal segment of PflA is a member of the nitrile hydratase-derived leader peptides (NHLPs),^27^ but the logo obtained from the multisequence alignment does not reveal a typical double-Gly-type cleavage site for a peptidase domain-containing ABC transporter (Figure 2B and S2).^28,29^ Indeed, a study focused on a similar BGC in *Pseudomonas* sp. Os17, a biocontrol strain, suggested that the full length precursor peptide was present in whole cell lysates based on Western blot analysis.^30^ Posttranslational modifications on this precursor peptide were not characterized in this prior work. ClusterBlast analysis^31^ showed that this type of BGC is prevalent among Pseudomonas species (Figure S1).

**Figure 2.**
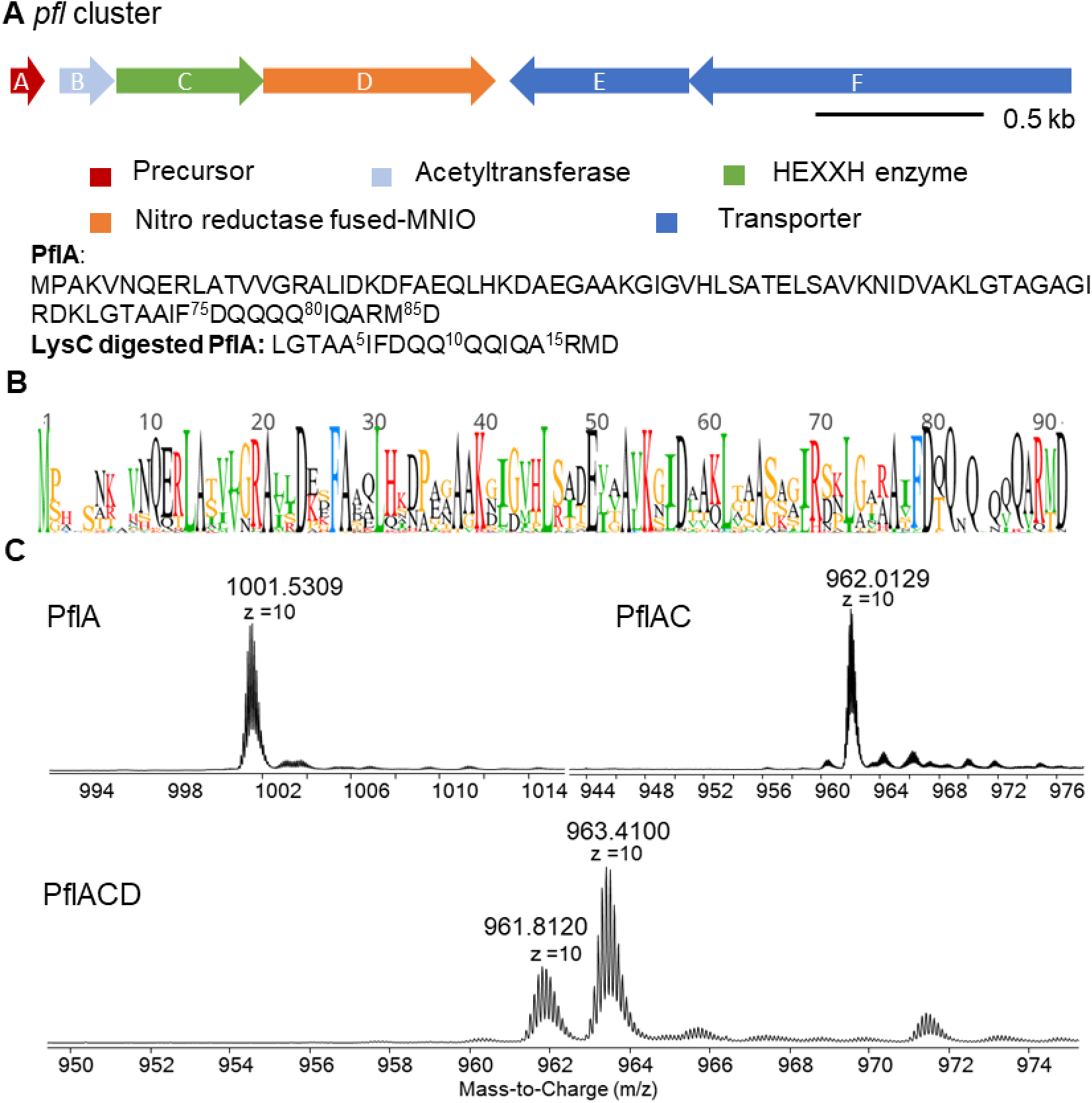
(A) Pfl BGC with the precursor peptide sequence. (B) Sequence logo of precursor peptides. (C) HR-LC-MS spectra of unmodified PflA (calculated m/z 1001.5178, z = 10), co-expressed PflAC (calculated average m/z 962.0599, z = 10), and co-expressed PflACD (calculated average m/z 963.4579, 961.8584, z = 10). Observed average masses are shown. See also Figure S5 for the determination of mass changes.

### Identification of the Posttranslational Modifications of PflA Peptides

We heterologously expressed the N-terminally His_6_-tagged precursor peptide (PflA) with individual enzymes of the *pfl* BGC in *Escherichia coli* using codon optimized genes (Table S1). After Ni^2+^ affinity purification, analysis by high-resolution mass spectrometry (HRMS) demonstrated that the peptide co-expressed with PflC (denoted PflAC) underwent a mass loss of −394 Da (Figure 2C and Figure S5). When PflA was co-expressed with PflD, two products were observed that were changed by +14 Da and −2 Da (Figure S3). Considering that PflD is a fusion of two enzymes, the observed mass changes are likely a combination of −2 Da and +16 Da modifications. To confirm the individual roles of the MNIO and nitroreductase domains, we coexpressed PflA with PflD-H73A in which His73 was mutated to Ala. His73 was expected to coordinate to Fe1 based on structures of other MNIOs^32–34^ as well as an AlphaFold3 model discussed later. The desaturated PflAD (−2 Da) was formed exclusively by this mutant enzyme (Figure S4), strongly supporting that the MNIO is responsible for the +16 Da modification. When PflA was co-expressed with both PflC and PflD, mass changes of −380 Da and −396 Da were observed, consistent with the individual mass shift assignments of PflC and PflD (Figure 2C, S5 and S6).

As noted above, the BGC does not contain a gene encoding a protease. Therefore, to help localize the modifications, we treated the PflCD-modified PflA (denoted as PflACD) with the commercial endoproteinase LysC. Analysis of the proteolysis products by MS showed that the modifications were all located in the C-terminal fragment (Figure S5). Co-expressions of PflA with PflBCD did not lead to an additional change in mass. In the Discussion section we provide possible explanations for the lack of PflB activity.

### Isolation and NMR characterization of Purified Peptides PosAC and Dehydro-PflAD

To assign molecular structures to the observed mass changes introduced by PflC and PflD, we first sought to purify the proteolytic fragments. The LysC-digested PflACD fragment was poorly soluble in most common buffers and organic solvents with or without acidification. Although this insolubility precluded isolation of PflACD for NMR analysis, the modified PflA peptide could still be characterized by MS/MS (see below). We then turned to analyze the post-translational modifications (PTMs) in the homologous BGC from *Pseudomonas sp*. Os17 mentioned in the introduction (*pos* BGC, Figure S7, Table S2). A −410 Da mass change was observed in PosA after co-expression with PosC in *E. coli* (Figure S8). LysC-digestion of the PosAC product resulted in precipitation of the C-terminal fragment with all other fragments remaining in solution, which allowed the isolation of pure PosAC peptide (see Supporting Information for details). Unfortunately, the purification method for PosAC did not allow analogous isolation of LysC-digested PflACD or PflAC.

For the discussion of our NMR and MS data of the LysC-digested peptide, we designate the first Ile residue of the C-terminal proteolytic peptide as residue 1 (Figure 3A, 3B). The purified LysC-digested PosAC peptide was dissolved in d_6_-DMSO. Twelve backbone amide protons were observed in the ^1^H and 2D ^1^H-^1^H TOCSY spectra (Figures S9, S10). Together with 2D ^1^H-^1^H NOESY, ^1^H-^13^C HSQC, and ^1^H-^15^N-HSQC spectra (Figures S9, S10), a complete assignment of the peptide was achieved (Table S3), indicating a 13 amino acids sequence of IGVAAIFDTQQQQ. These NMR data indicated that the last four residues ARMD at the C-terminus of PosA were removed, explaining why the C-terminal LysC fragment that was expected to consist of 17 residues, only showed twelve backbone amide signals. In the ^1^H-^15^N HSQC spectrum, in addition to the four pairs of NH_2_ side chain signals from the four Gln residues, an extra pair of amido NH_2_ signals were detected (Figure S10A). These latter signals showed NOESY cross peaks with the side chain amide and α/β protons of the last Gln residue (Figure 3A), suggesting C-terminal amidation. The ^1^H-^13^C HSQC spectrum displayed multiple downfield CH (methine) cross peaks around 69 ppm that were not anticipated based on the sequence of the peptide, suggesting a nearby electronegative atom (Figure S9C). The TOCSY and ^1^H-^13^C HMBC data showed that each of the four methine signals were part of Gln spin systems in which one of the expected CH_2_ groups was replaced by the methine signals. This conclusion was confirmed by the ^1^H-^13^C HSQC spectrum, and is consistent with hydroxylations of four Gln residues. Detailed analysis of the TOCSY and NOESY data (Figure S10B and S11) confirmed that the OH groups were installed at the β-position of the four Gln side chains. Collectively, these data demonstrate that PosC catalyzed β-hydroxylation of four Gln residues and removal of the ARMD-motif to generate a terminal amide. The net result of these modifications explain the observed mass change of −410 Da (4 x 16 = +64 Da; loss of ARMD = −490 Da; C-terminal amidation +16). The conclusions from the NMR data were corroborated by tandem mass spectrometry. HR-MS/MS analysis (Figure 3B) clearly demonstrated a 15.99 Da mass gain of the four Gln residues and confirmed an amidated C-terminus. These data suggest that the observed −394 Da change observed for PflAC is the result of oxidative removal of the ARMD motif in this peptide resulting in an amide, and hydroxylation of the five Gln residues in the sequence preceding the ARMD motif.

**Figure 3.**
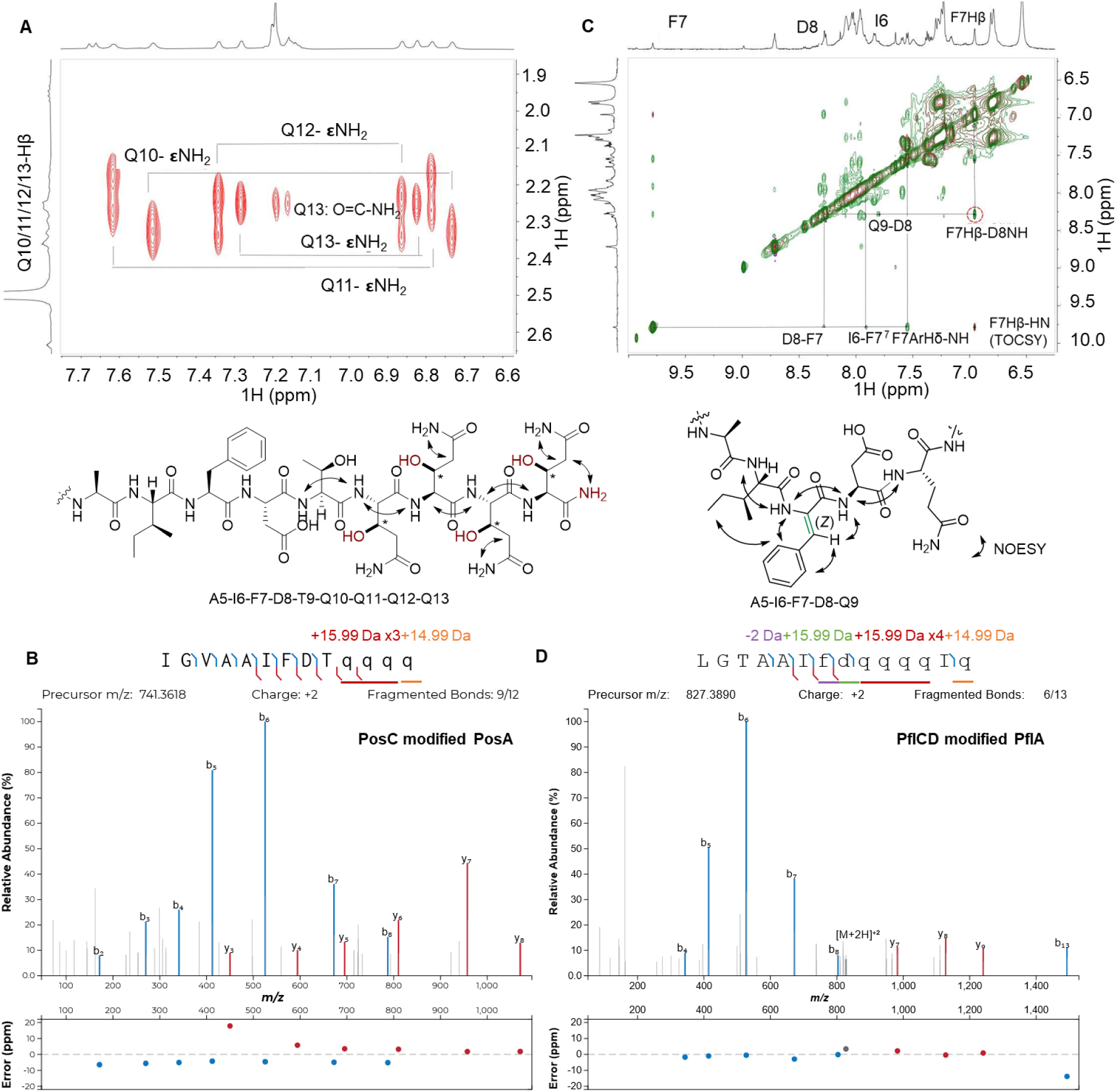
Structure determination of PosAC and dehydro-PflAD after digestion with LysC. (A) NOESY correlations of the C-terminal PosAC LysC-fragment. For the TOCSY spectrum, see Figure S10B. (B) HR-MS/MS spectrum of the C-terminal PosAC LysC-fragment. (C) Overlay of the aromatic region of the TOCSY-NOESY spectra of the dehydro-Phe peptide (dehydro-PflAD). (D) HR-MS/MS spectrum of the C-terminal PflACD LysC-fragment. Fragment ion annotation and ppm errors were generated using the interactive peptide spectral annotator^35^ with residues indicated in lower case q and d entered as hydroxylated residues (M+15.99), and in lower case f as dehydroPhe (–2 Da). The C-terminal Gln was entered as an α-carboxamide.

Because PosD was found to be inactive in *E. coli*, we returned to the *pfl* BGC to characterize the modifications that PflD introduced into PflA (denoted as PflAD) by co-expression of PflA and PflD in *E. coli*. After immobilized metal affinity chromatography (IMAC) purification, LysC digestion, and RP-HPLC purification, 2 mg of a product fragment was isolated from 2 L culture as a mixture of peptide that had undergone a −2 Da change (denoted as dehydro-PflAD) and unmodified PflA. We failed to isolate a pure minor product in PflAD with a mass gain of +14 Da (hydroxy-PflAD) due to its low concentration. We then characterized dehydro-PflAD by a series of one-dimensional (1D) and two-dimensional (2D) NMR experiments.

LysC digestion of dehydro-PflAD generated an 18-mer peptide with the sequence of LGTAAI<u>FD</u>Q-QQQIQARMD. For our discussion of the NMR and MS data of this peptide, we designate its N-terminal Leu as residue 1 (Figure 3C and D). The modifications in the underlined Phe7-Asp8 segment within the sequence were unambiguously assigned based on TOCSY, NOESY and ^1^H-^13^C HSQC spectra (Table S4). The modified part of the peptide was identified via sequential NOESY cross peaks of amide protons at 8.06 ppm (Ala5), 7.91 ppm (Ile6), 9.79 ppm (Phe7), 8.28 ppm (Asp8), and 7.80 ppm (Gln9). The considerable downfield shift of the amide proton of Phe7 is consistent with a loss of 2 Da from this residue to form a dehydro-Phe. The absence of an α-proton at residue 7 and the observation of a singlet at 6.96 ppm consistent with a vinyl proton confirms generation of dehydroPhe. The configuration of the dehydro-Phe residue was established as *Z* by a strong NOE observed between the singlet vinyl proton of former Phe7 at 6.96 ppm and the amide proton at 8.28 ppm of Asp8 (Figure 3C and S12). NOE correlations were also detected between the aromatic side chain (7.55 ppm) of former Phe7 and the side chain protons of Ile6, further supporting a *Z* configuration.

All NMR assignments in PosAC and dehydro-PflAD were corroborated in the fully modified PflACD product by high-resolution tandem electron ionization mass spectrometry (HR-ESI-MS/MS) analysis of the LysC-digested PflACD fragment. The fragmentation patterns are consistent with hydroxylations of all five Gln residues, a C-terminal amide and removal of the ARMD sequence, and loss of 2 Da at Phe7 (Figure 3D). These MS/MS data further showed a 15.99 Da gain at Asp8 (Figure 3D). Based on the data described above in which mutation of His73 in the MNIO domain resulted in formation of dehydro-PflA and abolishment of the +14 Da product (Figure S6), dehydroPhe formation (−2 Da) is catalyzed by the nitroreductase domain of PflD whereas hydroxylation of Asp8 (+16 Da) is catalyzed by the MNIO domain. These two PTMs explain the net +14 Da change in mass in the minor product of hydroxy-PflAD.

### Stereochemistry of Hydroxylation by PflC and PflD

Enantiopure β-hydroxylated glutamine (or glutamic acid) is not commercially available. Therefore, we constructed a variant carrying a Q90N mutation in the PosA peptide (numbering here based on full length precursor peptide). When co-expressed with PosC, this mutant underwent the same modifications as the wild type (WT) PosA peptide, resulting in a −410 Da mass change (Figure S13). Marfey’s analysis of this mutant product alongside authentic standards assigned the hydroxylated Asn residue as having the (2*S*,3*R*) configuration (Figure S14A). While we cannot completely exclude the possibility that the WT substrate may result in a different stereochemical outcome for Gln hydroxylation, the close structural and biosynthetic parallels between the WT and Q90N variant suggest that they likely share the same stereoselectivity. Marfey’s analysis of the PflD-modified PflA alongside authentic standards assigned the hydroxylated Asp8 residue as having the (2*S*,3*S*) configuration (Figure S14B), which is different from the HEXXH PosC-hydroxylated Asn.

### In vitro Activity of PflC and PosC

To further investigate the unexpectedly complicated modification caused by the HEXXH enzymes, we reconstituted their activity in vitro. We first heterologously expressed His_6_-tagged PflC and PosC, and the corresponding precursor peptides (PflA and PosA) in *E. coli*, followed by IMAC purification. When incubated with their cognate modification enzymes (peptide to enzyme ratio 10:1) in the presence of Fe(II), ascorbic acid, and αKG, the fully modified peptides PflAC and PosAC were observed (Figure 4A). Control experiments showed that αKG was essential for the activity, and ascorbic acid enhanced the catalytic efficiency (Figure 4A). Omission of Fe(II) resulted in minor activity, likely due to some trace metal in the protein during purification (Figure 4A). We also investigated cross-activity by incubating PflA with PosC or PosA and PflC under standard conditions, and in both cases, incomplete hydroxylation and deletion of the ARMD motif were observed (Figure S15). The reconstituted in vitro activity also allowed identification of the second product from peptide backbone cleavage. Based on other oxidative peptide backbone cleavage events,^36^ we anticipated production of a ketone generated from the ARMD sequence. Therefore, we treated the in vitro reaction products with *O*-benzylhydroxylamine and observed the anticipated mixture of *E* and *Z* oxime products for both PflC and PosC reactions (Figure S16).

**Figure 4.**
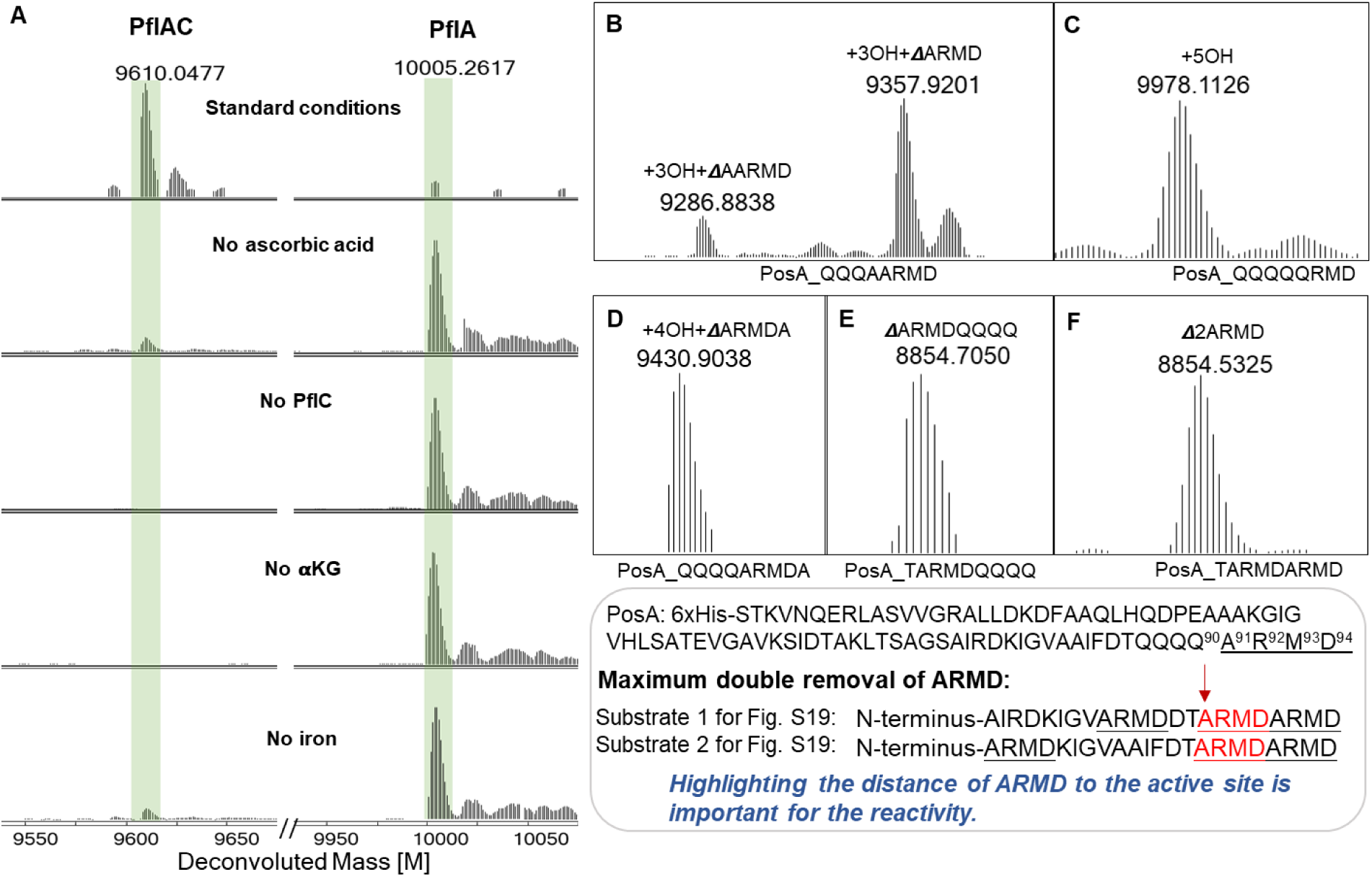
Peptide sequence tolerance test of PflC and PosC. (A) Experiments with full length PflA to determine the requirements for in vitro activity. (B) ESI-MS of PosA-Q90A modified in *E. coli* by PosC. (C) ESI-MS of PosA-A91Q modified in *E. coli* by PosC. (D) ESI-MS of PosA-QARMDA modified in *E. coli* by PosC. (E) ESI-MS of PosA-ARMDQQQQ modified in *E. coli* by PosC. (F) ESI-MS of PosA-2ARMD modified in *E. coli* by PosC. The sequences of the substrate are shown below the spectra, which display deconvoluted masses.

### Substrate Selectivity of PosC

We next evaluated the sequence selectivity of PosC. To probe the role of the C-terminal ARMD sequence, we performed an alanine scan at each position (PosA-R92A, PosA-M93A, and PosA-D94A). Co-expression of these precursor variants with PosC demonstrated that the Arg, Met, and Asp residues are individually dispensable, as all three mutants were fully modified to a product bearing four hydroxylations of the consecutive Gln residues with concomitant oxidative removal of the last four residues (AXXX; Figure S17). We also generated the PosA-Q90A variant to form a C-terminal QQQ-AARMD sequence; co-expression with PosC yielded a major product corresponding to hydroxylations of the three Gln residues with deletion of ARMD and a minor product corresponding to hydroxylation of the three Gln residues with deletion of AARMD. These observations indicate that the enzyme preferentially acts on Ala in the ARMD motif rather than an Ala after a series of (hydroxylated) Gln residues (Figure 4B). Substituting Ala91 with Gln (PosA-A91Q) abolished peptide scission and resulted in a product with hydroxylation of all five Gln residues in the variant peptide (Figure 4C). Likewise, the double mutant A91Q/R92A afforded the same five-hydroxylated species without AMD cleavage (Figure S18). When the C-terminal ARMD was deleted (PosA_ΔARMD), hydroxylation of two Gln residues was observed as the major product, along with a minor triple hydroxylation product (Figure S19), suggesting the C-terminal ARMD sequence enhanced Gln hydroxylation efficiency.

To test whether ARMD must reside at the C-terminus for peptide backbone cleavage, we inserted an alanine after ARMD (PosA-ARMDA) and in another variant, swapped the positions of ARMD and the four Gln residues (PosA-ARMDQQQQ). In both cases, PosC consistently cleaved immediately upstream of the ARMD sequence (Figure 4D and E), illustrating that the ARMD motif does not need to be at the C-terminus for oxidative cleavage. We also prepared a variant harboring two consecutive ARMD motifs by mutating the four consecutive Gln residues to ARMD (PosA-ARMDARMD). Coexpression of this variant with PosC produced exclusively the double-deletion product (Figure 4F). Conversely, two variants that contained an additional third ARMD motif further upstream did not show any cleavage at the first (most N-terminal) ARMD motif. These variants underwent cleavage at either the second or third ARMD sequence or both (Figure 4, bottom box and Figure S20). Collectively, these results suggest that productive cleavage requires an ARMD motif that does not need positioning at a specific location within the peptide as long as the motif is a certain distance of the N-terminal leader peptide. Cleavage also does not appear to be dependent on the residue before the cleavage site because the ARMD motif can be preceded by a hydroxylated Gln, a hydroxylated Asn, an Ala, or a Thr (Figures 4B,C,E,F and Figure S13).

Amidation by oxidative enzymes is a process that has been observed previously including for the peptidylglycine α-amidating monooxygenase (PAM)^37^ and the MNIO MovX.^38^ These enzymes usually catalyze hydrogen atom transfer (HAT)^39^ from the α-carbon of the amino acid where backbone cleavage takes place. A β-HAT pathway is also possible for PosC/PflC, and would be consistent with the observed β-hydroxylation of the Gln residues. Therefore, we prepared PosA-A91S as a possible intermediate of Ala oxidation. Upon co-expression with PosC, the peptide displayed hydroxylation at the four Gln residues and an additional −2 Da mass shift, while the backbone remained intact. LysC digestion and HR-MS/MS localized the −2 Da shift to Ser91, consistent with oxidation of the Ser side-chain primary alcohol to an aldehyde (Figure S21 and S22). Although the outcome with the PosA-A91S variant suggests that PosC can abstract a hydrogen atom from the β-position at position 91, as with the Gln hydroxylations, the observation that this did not result in backbone cleavage for both A91S and A91Q variants suggests that amide bond cleavage is achieved by hydrogen atom abstraction from the α-carbon, as in other α-amidating enzymes, and that an Ala residue is required for this process. The identification of the ketone products originating from the ARMD motif (Figure S16) strongly supports this mechanistic proposal.

To further investigate the requirements for backbone cleavage we generated two more variants, PosA-A91G and A91V. These variants were chosen to investigate two proteinogenic residues that are closest in size and structure to Ala, with a smaller amino acid (Gly) and a larger aliphatic amino acid (Val). After coexpression with PosC, four hydroxylations with removal of GRMD was observed for PosA-A91G, along with four and five hydroxylations without cleavage of the backbone (Figure S23). Given the structure of Gly, the product with five hydroxylations without cleavage must be hydroxylated on the α-carbon of Gly91. This hemiaminal product is reminiscent of the thiohemiaminal product produced by the MNIO TglHI that was stable.^40^ We also note that amidation of peptide hormones in mammals and insects requires a two step process, with the peptidyl α-hydroxylating monooxygenase (PHM) first catalyzing formation of an α-hydroxylated Gly, and then peptidylamidoglycolate lyase (PAL), catalyzing the hydrolysis of this intermediate to generate an α-amidated peptide and glyoxylate.^36^ These precedents illustrate the relatively high stability of hemiaminals in peptides. The observation of complete backbone cleavage with the natural substrate, but only partial cleavage of 5-fold hydroxylated PosA-A91G, suggests that PosC may catalyze the hydrolysis of the hemiaminal product for the wild type PosA peptide, akin to PAL catalysis, and that the presence of the methyl group on Ala91 is important for this step. The stability of the five-fold hydroxylated peptide is not unlimited, however, because it was mostly consumed upon endoproteinase LysC digestion (Figure S24), and attempts at tandem MS analysis were unsuccessful.

In contrast, no backbone cleavage was observed with PosA-A91V. Instead, mostly four-and five-fold hydroxylated peptides were formed (Figure S23). Digestion with LysC did not decrease the amount of the five-fold hydroxylated peptide (Figure S24), and analysis of this peptide by tandem MS showed that one of the five hydroxylations is clearly on Val91 (Figure S25). The observation of the lack of backbone cleavage, the stability of the five-fold hydroxylated product, and the fragmentation pattern is most consistent with hydroxylation taking place on the β-carbon of Val91, like the hydroxylations on the Gln and Asn residues described above. From the collective observed outcome with the mutants analyzed (Gln, Ser, Gly and Val), we infer that residues with small side chains, such as Ala and Gly, favor α-hydroxylation and backbone cleavage, while larger side chains (Gln, Val, Ser) result in β-hydroxylation without backbone cleavage.

### PflC Can Catalyze Oxidative and Hydrolytic Peptide Backbone Cleavage

Considering the apparently high sequence specificity of the HEXXH enzymes PosC and PflC for peptide backbone fragmentation, we turned to investigate the minimal sequence requirement of PflC to assess whether the N-terminal part of the substrate functions as leader peptide. We used solid-phase-peptide synthesis (SPPS) to prepare several truncated *N*-acetylated PflA precursor peptides (Figure S26). PflC was incubated with the truncated substrates in the presence of Fe(II), ascorbic acid and αKG. Hydroxylation activity was not observed, but unexpectedly, C-terminal hydrolytic release of ARMD was detected (Figures S27, S28). This process left a C-terminal carboxylic acid instead of amide, as confirmed by HR-MS/MS (Figure 5 and Figure S28). When the reaction was performed in H_2_^18^O-containing buffer, a +2 Da mass increase in the product confirmed that the oxygen atom originated from water (Figure 5). This observation established peptidase activity of PflC in the absence of the N-terminal sequence of PflA. These observations suggest that in this pathway, like for most RiPPs, the N-terminus of the precursor peptides serves as a leader peptide that ensures proper biocatalytic transformations. The hydrolytic activity was unexpected, but given the similarity of the PflC enzyme to the zincin protease family, we probed the metal dependency of the hydrolytic activity. We first generated apo PflC (see Experimental Section), reconstituted the protein with Zn(II), Co(II), Fe(II), or Fe(III) in individual experiments, and treated these enzymes with the truncated PflA peptides. Hydrolytic product was observed in all cases as with WT PflC (Figure S29), while no products were detected without adding any transition metal ions to the apo protein. These results demonstrate that the enzyme can use different metal ions, presumably as a Lewis acid, to facilitate peptide hydrolysis. We also investigated if hydrolysis can occur with full length PflA when oxygen is not present. Incubation of PflA with PflC in an anaerobic box for 20 h did not show any hydrolysis, suggesting that hydrolysis occurs from a distinct conformation that is attained only when the leader peptide is absent.

**Figure 5.**
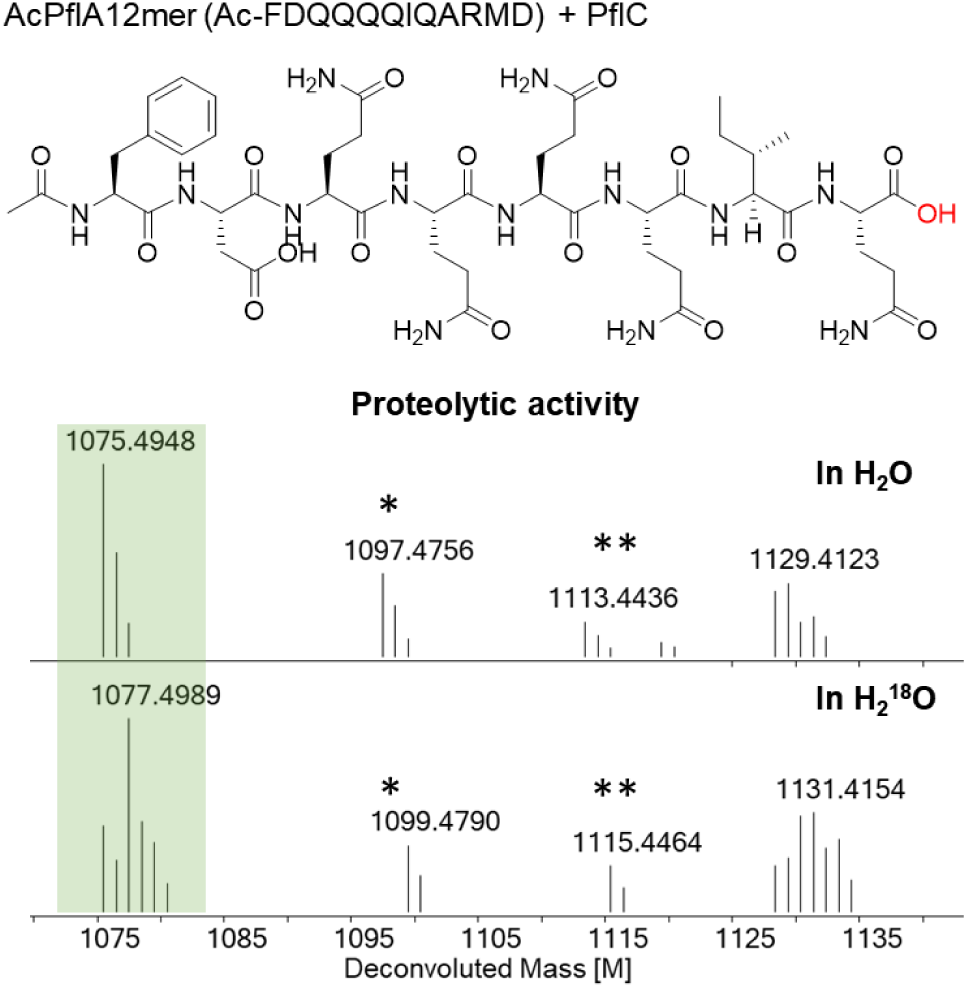
Proteolytic product obtained from a synthetic truncated PflA peptide shown at the top (also see Figures S26 and S27). An isotopic labeling experiment in H_2_^18^O labeled buffer indicated the source of the oxygen in the product is water. Calculated [M] 1075.4934 in H_2_O, [M] 1077.4977 in H_2_^18^O. * Na^+^ form of the ion; ** K^+^ form of the ion.

## Discussion

The investigation of the *pfl* and *pos* BGCs in this study revealed unexpected catalytic activities that are unprecedented in RiPP biochemistry. Similar BGCs are found in other *Pseudomonas* strains (Figure S1). Given the lack of a protease gene in the BGCs, we were not able to investigate the function of the final natural product of the pathways. Although it is possible that the full length products of PflA and PosA are the mature products, in all previously characterized RiPPs removal of a leader/follower peptide is involved in formation of the final bioactive product.^1,26^ The presence of a gene encoding an acetyl transferase PflB, which did not display any activity on full length peptides, also suggests that it will act after removal of an N-terminal leader peptide; N-terminal acylation by a GNAT-like enzyme has been reported for several RiPPs.^21,41–43^ N-terminal acylation, as opposed to acylation of Lys side chains,^21,44^ is supported by the observation that the sequence logo of the precursor peptides (Figure 2B) does not reveal a highly conserved Lys residues in the precursor peptides. We cannot rule out that a non-specific aminopeptidase encoded elsewhere in the genome may act on the modified peptides, as observed for several RiPPs.^25^ In that scenario, the aminopeptidase digestion could possibly terminate at the *Z*-dehydroPhe, resulting in formation of an N-terminal ketone upon hydrolysis (Figure S30), similar to the known generation of N-terminal 2-oxobutyryl and 2-oxopropionyl groups from dehydrobutyrine and dehydroalanine, respectively.^45^ If this were the case, the GNAT might acylate one of the introduced hydroxyl groups similar to the recently reported acylation of the hydroxyl group of a Tyr residue in a lasso peptide.^46^

The HEXXH enzymes PflC and PosC both catalyze hydroxylation of the β-carbon of a string of Gln residues, as well as oxidative removal of a C-terminal ARMD tetrapeptide. These two transformations appear mostly independent because hydroxylation of Gln was still observed when the ARMD motif was deleted, and backbone cleavage was still observed when the Gln residues were removed. Because we only see a product with four Gln hydroxylations and backbone cleavage with the wild type PosA substrate, whereas a peptide lacking the ARMD motif produced mostly a product with two hydroxylations (Figure S19), we infer that Gln hydroxylations precede cleavage of the backbone. However, we cannot rule out that the amide-terminated backbone-cleaved peptide is a more efficient substrate for Gln hydroxylation than the carboxylate-terminated peptide. The length of the tetrapeptide appears not critical as extension at the C-terminus still resulted in backbone cleavage. Other than the Ala residue, the individual residues of the ARMD motif also did not appear critical, despite their complete conservation (Figure 2B). A possible mechanism for backbone scission in which an initial radical generated on the β-carbon of the last Gln would abstract a hydrogen atom from the Ala residue of the ARMD motif to initiate cleavage also seems unlikely because for such a mechanism, insertion of an additional Ala (AARMD) would have been expected to result in cleavage of AARMD, but removal of ARMD was still observed. These observations further corroborate that modification of these Gln residues is not required for oxidative backbone cleavage. Furthermore, the variant in which all Gln residues were replaced by another ARMD sequence was still cleaved.

To rationalize the site-specific backbone cleavage and amide formation, we propose that Ala91 undergoes α-C–H abstraction by a high-valent iron–oxo species to generate an α-amino carbon radical stabilized by the neighboring amide nitrogen. This intermediate could either (i) undergo oxygen rebound to form an α-hydroxylated species that eliminates water to an iminium, followed by hydrolytic cleavage of the peptide backbone, or (ii) be oxidized by one electron directly to the same iminium intermediate (Figure S31). The observed hydrolysis at the exact same site as the oxidative backbone cleavage when a truncated substrate was used suggests that the truncated peptide binds in the exact same position near the metal ion, but that without the N-terminal substrate sequence, oxygen activation and/or αKG binding does not occur. Substrate gating has been shown to control canonical αKG-dependent enzymes,^47^ where productive oxygen activation by Fe(II) only occurs when both substrate and αKG are bound in the active site. Our observations with PflC/PosC suggest that the αKG-dependent HEXXH enzymes studied here are similarly substrate-gated for oxidative chemistry. Furthermore, the data suggest that binding of the N-terminal sequence of the substrate appears critical for oxygen activation, consistent with this segment of the substrate acting as a leader peptide, as in other RiPP biosynthetic pathways.^48^ Other observations are also suggestive of an important role of the N-terminal sequence for the regioselectivity of backbone cleavage. When three ARMD motifs were introduced into the precursor peptides, backbone cleavage was only observed at positions that are located towards the C-terminus of the substrates. Installation of the ARMD motif at positions that were more towards the center of the peptide did not result in cleavage (Figure S20). One plausible explanation is that an N-terminal leader peptide positions the C-terminal part of the substrate near the Fe(II) site and that ARMD motifs that are closer to the leader peptide cannot reach the active site. The observation that PosA was not a good substrate for PflC, and PflA was not a good substrate for PosC, could also be explained by differences in their N-terminal sequences.

An AlphaFold3 model of PosC in complex with PosA or PflC with PflA places the iron center right next to the Ala residue in the C-terminal ARMD motif (Figures S32, S33), consistent with the observed site-selective cleavage. The AlphaFold3 model also predicts a well-folded N-terminal region of PosA that resembles the recently reported NMR structures of nitrile hydratase-like leader peptides (Figure S33).^49,50^ These NHLPs are involved in enzyme recognition, but the manner of substrate recognition as well as the region on the long leader peptides used for enzyme engagement differs significantly for different pathway enzymes. By analogy, we anticipate that the PflA/PosA leader contributes to enzyme recognition, but the precise interaction requires further experiments. The AlphaFold3-predicted structure is also very similar to the crystal structure that was reported for the leader peptide of PosA in *Pseudomonas* sp. Os17.^30^ The PTMs catalyzed by the biosynthetic enzymes were not characterized in this prior study.

The nitroreductase domain of PflD catalyzes an atypical dehydrogenation of phenylalanine. Dehydroamino acids (dhAAs) play central roles in the biological activity of both RiPPs and non-ribosomal peptide natural products.^51,52^ In bacterial RiPPs, dhAAs are overwhelmingly represented by dehydroalanine (Dha) and dehydrobutyrine (Dhb), typically generated by dehydration/dehydrothiolation of serine or cysteine residues (for Dha), and threonine residues (for Dhb).^1^ To the best of our knowledge, this study reports the first dehydrophenylalanine formation in RiPPs. Other nitroreductases in RiPP BGCs that are not associated with YcaO enzymes could be involved in the formation of other dehydroamino acids.

The MNIO domain of PflD catalyzes a reaction that has been observed in several other RiPPs, the hydroxylation of the β-carbon of an Asp residue. In other RiPPs, this reaction is usually catalyzed by either canonical αKG dependent enzymes,^16,18^ or by recently discovered αKG-dependent HEXXH enzymes.^12^ The observation that an MNIO catalyzes this reaction was unexpected because most enzymes in this family catalyze four-electron oxidations, which is thought to involve first hydrogen abstraction from a weak C-H bond adjacent to a heteroatom by a ferric-superoxo species, and then a second oxidation involving an iron(IV)-oxo intermediate.^7^ We previously proposed that the MNIO ApyH catalyzed a four-electron oxidation with a transiently formed β-hydroxylation of an Asp in a peptide.^53^ The prior examples of (potential) two-electron oxidation that are catalyzed by MNIO enzymes involved substrates that either can eliminate hydrogen peroxide,^38^ or that have electron rich π-type electrons.^8^ However, for hydroxylation of Asp, hydrogen atom abstraction is required from a non-activated C-H bond with a relatively high bond strength, a step that appears unlikely for a ferric-superoxo intermediate.^39^ Hence, we propose that the reaction catalyzed by the MNIO domain of PflD likely involves a cellular reductant that allows formation of a Fe(IV)-oxo intermediate.

### Conclusions

Two αKG-dependent HEXXH enzymes (PflC and PosC) were characterized in the biosynthesis of a RiPP family from Pseudomonas (Figure 6). Both enzymes catalyze hydroxylation of consecutive Gln residues and specifically recognize a C-terminal tetrapeptide ARMD to promote oxidative peptide backbone cleavage and generation of a C-terminal amide and a ketone containing by-product originating from the tetrapeptide. The PflC enzyme also exhibits proteolytic activity in the absence of the leader peptide, indicating that leader peptide–enzyme interactions can regulate catalytic outcomes. Further investigation is required to elucidate the mechanisms underlying these distinct catalytic selectivities. In addition, we identified a unique MNIO–nitroreductase fusion enzyme that produces a rare dehydrophenylalanine and hydroxylates aspartate. Together, these findings describe new catalytic activities that expand the catalog of post-translational modifications and set up studies to characterize the function of the product of the BGCs.

**Figure 6.**
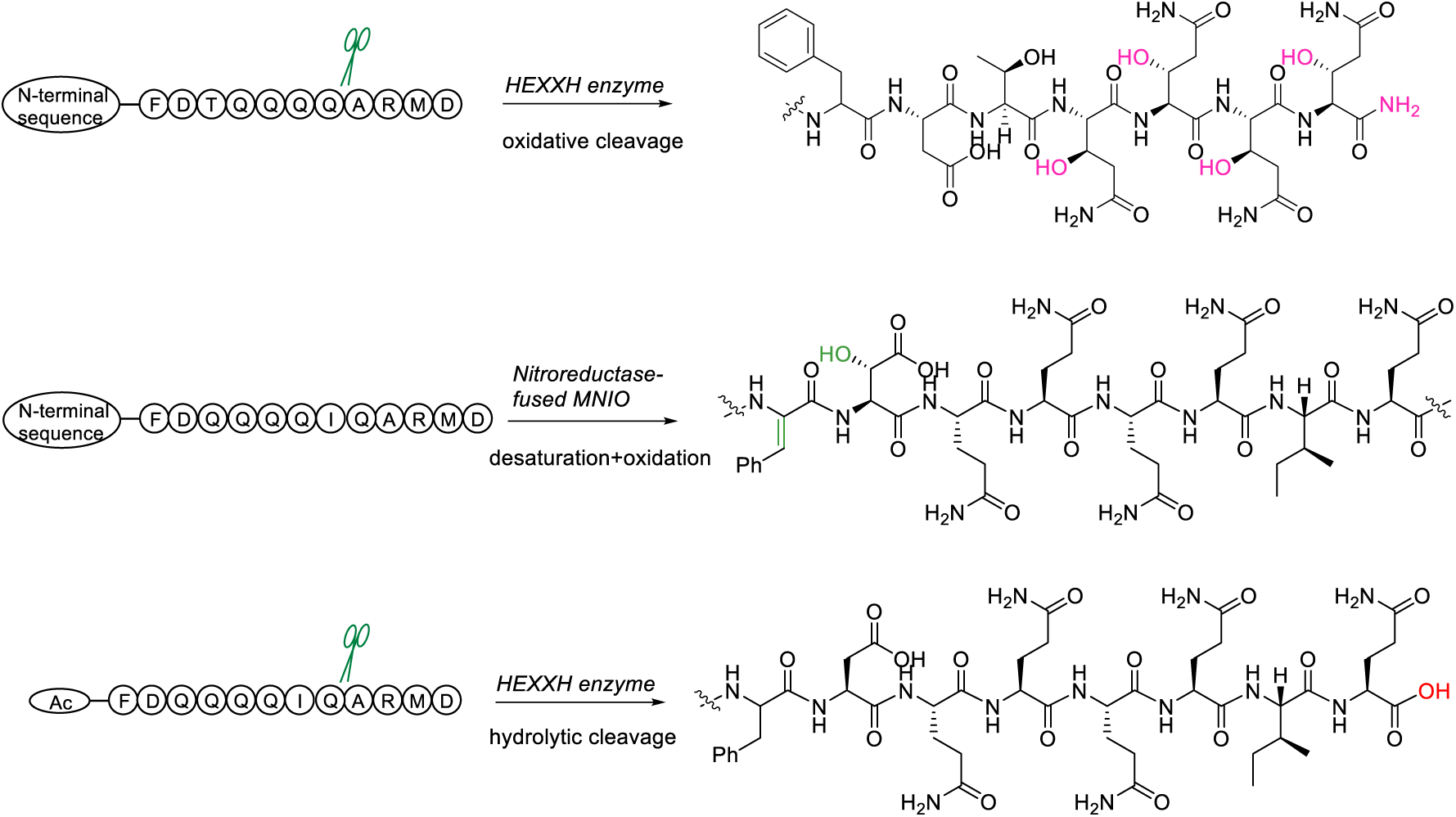
Summary of enzymatic activities of the HEXXH enzymes and nitro-reductase-fused MNIO.

## ASSOCIATED CONTENT

### Supporting Information

The Supporting Information is available free of charge at https://pubs.acs.org/doi/

Experimental procedures, Figures S1-S29 showing BGC information, spectroscopic data, NMR annotations, and AlphaFold3 modeling, and Tables S1-S4 with gene sequences and primers (PDF).

## AUTHOR INFORMATION

### Corresponding Author

Wilfred A. van der Donk −Department of Chemistry and Howard Hughes Medical Institute, University of Illinois at Urbana-Champaign, Urbana, IL, United States.

### Authors

Yao Ouyang – Department of Chemistry and Howard Hughes Medical Institute, University of Illinois at Urbana-Champaign, Urbana, IL, United States.

Yue Yu – Department of Chemistry and the Howard Hughes Medical Institute at the University of Illinois at Urbana-Champaign, 1206 W Gregory Drive Urbana, IL 61801, United States.

Lingyang Zhu – School of Chemical Sciences NMR Laboratory, University of Illinois at Urbana-Champaign, Urbana, IL, United States.

Dinh Nguyen – Department of Chemistry and the Howard Hughes Medical Institute at the University of Illinois at Urbana-Champaign, 1206 W Gregory Drive Urbana, IL 61801, United States.

### Notes

The authors declare no competing financial interests.

## ACKNOWLEDGMENTS

We thank Dr. Chandrashekhar Padhi for assistance with bioinformatic analysis, Dr. Lide Cha for providing standards for Marfey’s analysis, and Youran Luo for assistance with Marfey’s analysis. We also thank Dr. Chandrashekhar Padhi, Ceilia Leso, and Pelham Kogut for assistance with mass spectrometry.

---

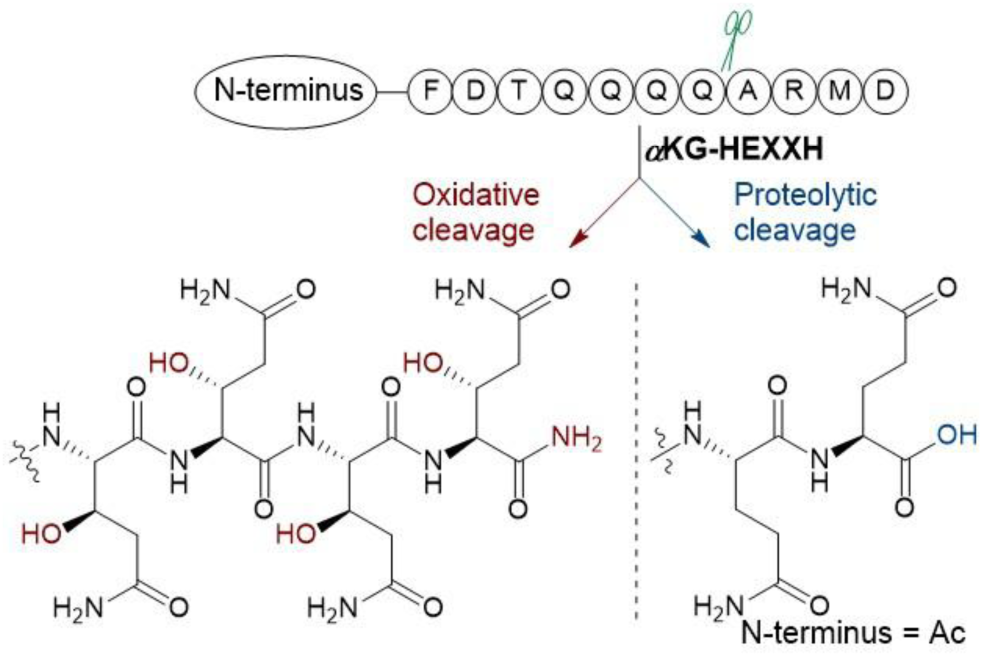
TOC graphic.

