## Supplemental figures and tables for "Oxidative Peptide Backbone Cleavage by a HEXXH Enzyme During RiPP Biosynthesis"

### Materials and methods

Molecular biology experiments were performed using reagents from New England Biolabs, Thermo Fisher Scientific, or Sigma-Aldrich. Plasmid maintenance and protein overexpression were conducted using *Escherichia coli* DH10β and BL21 (DE3), respectively. MALDI-TOF-MS data acquisition was performed using a Bruker UltrafleXtreme mass spectrometer (Bruker Daltonics) in reflector positive mode at the University of Illinois School of Chemical Sciences Mass Spectrometry Laboratory. A commercial mixture of 9:1 2,5-dihydroxybenzoic acid (DHB) and 2-hydroxy-5-methoxybenzoic acid (Super-DHB or SDHB) solution with a stock concentration of 25 mg/mL was routinely mixed with samples in a 1 to 1 ratio and air-dried before MALDI-TOF-MS analysis. MS1 and MS2 data acquisition was performed using an Agilent qTOF instrument equipped with an UPLC system. 2D NMR experiments were recorded using a Bruker 600 MHz spectrometer in d6-DMSO with or without 0.2% d-formic acid. Ni-NTA resin for peptide and protein purifications was obtained from ThermoFisher.

Genome mining and the generation of a sequence similarity network (SSN) of $\alpha$-KG-HEXXH proteins was created using the EFI-EST webtool[1] and using the Interpro family IPR026337 as input, accounting for ca. 3058 entries as of September 2025. An alignment score of 30 was applied for network generation. Cluster coloring was conducted using the EFI-EST's Color SSNs functionality and the resultant representative node (rep-node) 40, where sequences over 40% identity threshold were grouped into a meta-node, was visualized in Cytoscape v3.10 (see additional Supporting Information file). Uniprot IDs of the HEXXH hits were extracted from the SSN and analyzed utilizing the RODEO webtool.[2] Sequence alignment and sequence logo generation were conducted by PHI-blast using PosA as query and aligned with the Geneious tool using the Muscle 5.1 parallel perturbed probcons (PPP) algorithm through 100 hidden Markov model (HMM) perturbations.[3]

### Plasmid construction

Codon-optimized (for *E. coli*) synthetic genes encoding precursor peptides were ordered as gBlocksTM from Integrated DNA Technologies (IDT) or adapter-removed gene fragments from Twist Bioscience with sequences homologous to the 5' and 3' ends of the plasmid vector insertion site.

Plasmid DNA was prepared using a MiniPrep kit (Qiagen) following the manufacturer's instructions from *E.*

*coli* DH10β cells. Plasmids were manually designed and prepared by Gibson assembly using NEBuilder HiFi DNA Assembly Master Mix. The vector primer design usually incorporated overhangs complementary to the gene of interest. PCR amplification was performed using Q5 polymerase, dNTPs and Q5 reaction buffer from New England Biolabs. Designed primers for cloning and in vitro transcription were acquired from Integrated DNA Technologies.

**Heterologous expression and purification of peptides/proteins**

Bacterial growth and protein production were conducted with 50 µg/mL kanamycin in Luria Bertani Broth supplemented with 1x final concentration of trace metal mix (Teknova). A 10 mL overnight starter culture was used to inoculate a 1 L culture and grown at 37 ºC until the optical density at 600 nm (OD600) reached ~0.8. The cells were then transferred to ice for 15 min followed by the addition of 0.2 mM IPTG to induce gene expression. Induced cells were further cultivated at 18 ºC overnight at 200 rpm. Cells were then harvested by centrifugation at 4500 ×g for 10 min. Cell pellets were stored in a −80 °C freezer for future use or resuspended in denaturation buffer for peptides (50 mM Tris-HCl buffer containing 300 mM NaCl, 10% glycerol, and 6 M guanidine hydrochloride at pH 7.5). For proteins, the cell pellets were resuspended in start buffer (50 mM HEPES buffer containing 300 mM NaCl, 10% glycerol, pH 7.5). The resuspended cells were disrupted by sonication using repeated cycles of 2 s on and 5 s off. After centrifugation at 50,000 ×g, clarified lysate was incubated with Ni-NTA resin at 4 ºC for 1 h before loading on to a column. The column was washed with 10 column volumes (CV) of wash buffer (denaturing buffer with 30 mM imidazole). The column was washed with the start buffer supplemented with 30 mM and 50 mM imidazole. The His-tagged peptide/protein was then eluted with 5 CV of elution buffer composed of 300 mM imidazole, 300 mM NaCl, and 50 mM HEPES at pH 7.5.

**Apo PflC/PosC production**

The protein purification step is the same as the general procedure described above. After elution from the Ni-NTA resin, the resulting protein was dialyzed against 50 mM HEPES buffer containing 300 mM NaCl, 10% glycerol, and 15 mM EDTA. After overnight incubation at 4 °C, dialysis with EDTA-free buffer was performed three times (buffer change every 3 h). The resulting protein was concentrated, aliquoted, flash-frozen, and stored at −80 °C. This enzyme was devoid of any activity.

**Isolation of modified peptide core fragments**

Following IMAC purification, the peptide eluent was desalted and the buffer was exchanged to a new buffer (50 mM Tris-HCl, 100 mM NaCl at pH 8.0) using a Sephadex PD10 column. Fully length peptides were directly subjected to HR-LC-MS analysis using a C4 column. For peptides characterized by NMR spectroscopy, proteolysis was conducted at 37 °C overnight using LysC endopeptidase (NEB) at 1:500 enzyme:substrate ratio.

PflD-modified PflA was acidified with a 10% trifluoroacetic acid (TFA) solution to a final concentration of 0.1 to 2% TFA. The sample was then passed through a 0.45 µm syringe filter and injected directly into an Agilent 1260 Infinity II HPLC instrument for purification. The peptide was purified using a preparative scale column, VP NUCLEODUR C18 HTec, 5 µm, 250x10 mm with the following gradient: 0-5 min isocratic 2% acetonitrile (ACN), 5-40 min 2%-60% ACN, 40-42 min 60%-100% ACN; mobile phase buffered with 0.1%

TFA. Fractions were collected manually and analyzed by MALDI-TOF MS by combining 1 to 1 with a 25 mg/mL solution of SDHB. Fractions containing the desired peptide were combined and lyophilized. The dried solids were then used for NMR analysis.

For LysC-digested PosC-modified PosA peptide, the sample was centrifuged at 4000 g for 10 min and the supernatant was discarded. The pellet was washed three times with $NaHCO_3$ (100 mM, pH 8) to remove any fragments that were dissolved in solution. After drying with a lyophilizer, the modified peptide was obtained as a white powder and dissolved in d6-DMSO for NMR analysis. Using this procedure, we isolated 2-3 mg modified peptide from 1 L cell culture.

**High-resolution tandem mass spectrometry**

The desalted protease-digested peptides were injected on to an Agilent 1290 LC-MS QToF for ESI-HRMS and MS/MS analysis. LC separation was conducted at 50 ºC on a 5%-95% gradient of ACN/water (+0.1% formic acid) over 10 min at 0.4 mL/min flow rate on a Phenomenex Aeris 2.6 µm PEPTIDE XB-C18 LC column (part nr. 00F-4505-E0). Mass spectra were collected in positive mode at 10 spectra/s and 100 ms/spectrum. Tandem-MS Fragmentation was collected at normalized collision energies of 20, 25 and 30 eV. HR-MS/MS analysis was performed using the Interactive Peptide Spectral Annotator (IPSA) tool,[4] and verified manually.

**Advanced Marfey's analysis**

The absolute configuration of the hydroxylated Asn90 (full length) was determined as previously described, with slight modifications.[5] Briefly, ~80 µg of dried peptide product from coexpression of PosA-Q90N with PosC (coPosA-Q90N) was hydrolyzed in 2 mL of 6 M DCl in $D_2O$ in a 10 mL pressured glass vial. The mixture was heated at 120 ºC overnight with stirring. The solvent of the resulting solution was removed under reduced pressure followed by resuspension in 1 mL of deionized water and dried again to remove any remaining acid. Next, 0.6 mL of 0.8 M $NaHCO_3$ solution and 0.4 mL of a 10 mg/mL solution of N-α-(5-fluoro-2,4-dinitrophenyl)-L-alanine amide (L-FDAA) in acetonitrile was added, followed by stirring in the dark (amber glass vial) at 67 °C for 6 h. Post-derivatization, 100 µL of 6 M HCl was added to neutralize the reaction, and the mixture was vortexed. The mixture was lyophilized and subsequently resuspended in 500 µL of acetonitrile by sonication. The suspension was centrifuged at 12,000g for 10 min, and the supernatant was analyzed by LC–MS on an Agilent 6545 LC/Q-TOF instrument. Chromatographic separation was obtained on a Kinetex 2.6 µm XB-C18 100 Å, LC column (Phenomenex; part no.: 00F-4496-Y0). The column oven was maintained at 50 °C, and the mobile phase used was A: water with 0.1% formic acid (FA) and B: ACN with 0.1% FA. At a constant flow rate of 0.4 mL/min, a gradient of 2–10% B over 2.5 min, and 10–80% B over the next 7.5 min was maintained, followed by a wash step of 95% B for 3 min and a post-run equilibration stage of 2% B for 5 min. MS spectra were collected in the negative ion polarity mode, and 0.5–1 µL injections were performed for each sample. For co-injection experiments, the data collected from the individual runs were used to roughly normalize the amount of each sample and standard compounds and each sample:standard compound was combined in 1:1 ratio, followed by LC-MS analysis.

**In vitro assays**

PflA or PosA (50 µM) was incubated with PflC or PosC (5 µM) in Tris buffer (100 mM, pH 8), with ascorbic acid (1 mM), ammonium iron(II) sulfate (0.5 mM), and alpha-KG (2 mM). The mixture was incubated at room temperature overnight. The reaction was quenched by adding 2x volume of ACN and a final concentration of 0.1% FA to precipitate any protein, followed by centrifuging at 15000 g for 10 min. The resulting supernatant was subjected to LC-MS analysis.

Primary data associated with the figures in this study have been deposited in: Ouyang, Yao; Zhu, L; van der Donk, Wilfred (2025), “Data associated with "Oxidative Peptide Backbone Cleavage by a HEXXH Enzyme During RiPP Biosynthesis"”, Mendeley Data, V1, doi: 10.17632/df2xckv3fr.1

**Table S1:** DNA sequences for constructs used in this study.

| Construct name | DNA sequence |
|---|---|
| His6-PflA | ATGGGCAGCAGCCATCACCATCATCACCACTCTACTAAAGTAAATCAGGAGCGACTC<br>GCGAGCGTCGTGGGTAGAGCCCTGCTCGACAAGGATTTTGCCGCCCAGCTCCACC<br>AGGATCCGGAGGCTGCGGCTAAAGGTATAGGCGTGCATTTAAGTGCGACGGAGGTG<br>GGTGCGGTCAAAAGTATCGACACTGCTAAACTCACATCAGCAGGCTCCGCAATCCG<br>CGATAAGATTGGAGTAGCTGCGATTTTCGACACTCAGCAGCAACAAGCTCGAATGGA<br>CTGA |
| Translated sequence | MGSSHHHHHHPAKVNQERLATVVGRALIDKDFAEQLHKDAEGAAKGIGVHLSATELSAV<br>KNIDVAKLGTAGAGIRDK LGTAAIFDQQQQIQARMD |
| His-TEV-PflB | actctagatttcagtgcaatttatctcttcaaatgtagcacctgaagtcagccccatacgatataagttgtaattctcatgttagtc<br>atgccccgcgcccaccggaaggagctgactgggttgaaggctctcaagggcatcggtcgagatcccggtgcctaatgagt<br>gagctaacttacattaattgcgttgcgctcactgcccgctttccagtcgggaaacctgtcgtgccagctgcattaatgaatcgg<br>ccaacgcgcggggagaggcggtttgcgtattgggcgccagggtggtttttcttttcaccagtgagacgggcaacagctgatt<br>gcccttcaccgcctggccctgagagagttgcagcaagcggtccacgctggtttgccccagcaggcgaaaatcctgtttgat<br>ggtggttaacggcgggatataacatgagctgtcttcggtatcgtcgtatcccactaccgagatgtccgcaccaacgcgcagc<br>ccggactcggtaatggcgcgcattgcgcccagcgccatctgatcgttggcaaccagcatcgcagtgggaacgatgccctc<br>attcagcatttgcatggtttgttgaaaaccggacatggcactccagtcgccttcccgttccgctatcggctgaatttgattgcga<br>gtgagatatttatgccagccagccagacgcagacgcgccgagacagaacttaatgggcccgctaacagcgcgatttgct<br>ggtgacccaatgcgaccagatgctccacgcccagtcgcgtaccgtcttcatgggagaaaataatactgttgatgggtgtctg<br>gtcagagacatcaagaaataacgccggaacattagtgcaggcagcttccacagcaatggcatcctggtcatccagcgga<br>tagttaatgatcagcccactgacgcgttgcgcgagaagattgtgcaccgccgctttacaggcttcgacgccgcttcgttctac<br>catcgacaccaccacgctggcacccagttgatcggcgcgagatttaatcgccgcgacaatttgcgacggcgcgtgcagg<br>gccagactggaggtggcaacgccaatcagcaacgactgtttgcccgccagttgttgtgccacgcggttgggaatgtaattc<br>agctccgccatcgccgcttccactttttcccgcgttttcgcagaaacgtggctggcctggttcaccacgcgggaaacggtctg<br>ataagagacaccggcatactctgcgacatcgtataacgttactggtttcacattcaccaccctgaattgactctcttccgggcg<br>ctatcatgccataccgcgaaaggttttgcgccattcgatggtgtccgggatctcgacgctctcccttatgcgactcctgcattag<br>gaaattaatacgactcactataggggaattgtgagcggataacaattcccctgtagaaataattttgtttaactttaataagga<br>gatataccatgGGCAGCAGCCATCATCATCATCATCACAGCAGCGGCGAGAACCTGTACT<br>TCCAAAGTatgagtgttacactgatcccgatggatgatgcggattttgttacttttaccgaacgggcagtggatgaatacg<br>ctcaaggtatggttgccagcggtgaatggagccaggcggaagcgtcacatcgctcagccacagcattcgctcaactcctg<br>gataaagggcgtctgacggaaaatcaccagttatgggtcattagagataatcagcgtcgaatcggagaattatggatcgcg<br>cgccggcatgttgggactaaaccgatagctttcatactggatatttatatagaaccgggtgaacgccgccaaggtcatgccc<br>gtcagtctatgttggcggcagaaacatgggcacgacagaatgactgcgaagaaatgcgactgcatgtatttggccgtaatc<br>tgcctgcgcgtggcctttatgaacagttaggttatgacatagcaagtcttactatggctaagccactgccacccgcataaagc<br>ttgcggccgcataatgcttaagtcgaacagaaagtaatcgtattgtacacggccgcataatcgaaattaatacgactcactat<br>aggggaattgtgagcggataacaattccccatcttagtatattagttaagtataagaaggagatatacatatggcagatctca<br>attggatatcggccggccacgcgatcgctgacgtcggtaccctcgagtctggtaaagaaaccgctgctgcgaaatttgaac<br>gccagcacatggactcgtctactagcgcagcttaattaacctaggctgctgccaccgctgagcaataactagcataacccc<br>ttggggcctctaaacgggtcttgaggggttttttgctgaaacctcaggcatttgagaagcacacggtcacactgcttccggtag |

| | |
|---|---|
| | tcaataaaccggtaaaccagcaatagacataagcggctatttaacgaccctgccctgaaccgacgacaagctgacgacc<br>gggtctccgcaagtggcacttttcggggaaatgtgcgcggaacccctatttgtttatttttctaaatacattcaaatatgtatccgc<br>tcatgaattaattcttagaaaaactcatcgagcatcaaatgaaactgcaatttattcatatcaggattatcaataccatatttttga<br>aaaagccgtttctgtaatgaaggagaaaactcaccgaggcagttccataggatggcaagatcctggtatcggtctgcgatt<br>ccgactcgtccaacatcaatacaacctattaatttcccctcgtcaaaaataaggttatcaagtgagaaatcaccatgagtga<br>cgactgaatccggtgagaatggcaaaagtttatgcatttctttccagacttgttcaacaggccagccattacgctcgtcatcaa<br>aatcactcgcatcaaccaaaccgttattcattcgtgattgcgcctgagcgagacgaaatacgcggtcgctgttaaaaggac<br>aattacaaacaggaatcgaatgcaaccggcgcaggaacactgccagcgcatcaacaatattttcacctgaatcaggatat<br>tcttctaatacctggaatgctgttttcccggggatcgcagtggtgagtaaccatgcatcatcaggagtacggataaaatgcttg<br>atggtcggaagaggcataaattccgtcagccagtttagtctgaccatctcatctgtaacatcattggcaacgctacctttgcca<br>tgtttcagaaacaactctggcgcatcgggcttcccatacaatcgatagattgtcgcacctgattgcccgacattatcgcgagc<br>ccatttatacccatataaatcagcatccatgttggaatttaatcgcggcctagagcaagacgtttcccgttgaatatggctcata<br>ctcttcctttttcaatattattgaagcatttatcagggttattgtctcatgagcggatacatatttgaatgtatttagaaaaataaaca<br>aataggcatgcagcgctcttccgcttcctcgctcactgactcgctacgctcggtcgttcgactgcggcgagcggtgtcagctc<br>actcaaaagcggtaatacggttatccacagaatcaggggataaagccggaaagaacatgtgagcaaaaagcaaagca<br>ccggaagaagccaacgccgcaggcgtttttccataggctccgcccccctgacgagcatcacaaaaatcgacgctcaagc<br>cagaggtggcgaaacccgacaggactataaagataccaggcgtttccccctggaagctccctcgtgcgctctcctgttccg<br>accctgccgcttaccggatacctgtccgcctttctcccttcgggaagcgtggcgctttctcatagctcacgctgttggtatctcag<br>ttcggtgtaggtcgttcgctccaagctgggctgtgtgcacgaaccccccgttcagcccgaccgctgcgccttatccggtaact<br>atcgtcttgagtccaacccggtaagacacgacttatcgccactggcagcagccattggtaactgatttagaggactttgtcttg<br>aagttatgcacctgttaaggctaaactgaaagaacagattttggtgagtgcggtcctccaacccacttaccttggttcaaaga<br>gttggtagctcagcgaaccttgagaaaaccaccgttggtagcggtggtttttctttatttatgagatgatgaatcaatcggtctat<br>caagtcaacgaacagctattccgtt |
| Translated sequence | MGSSHHHHHHSSGENLYFQSSVTLIPMDDADFVTFTERAVDEYAQGMVASGEWSQAE<br>ASHRSATAFAQLLDKGRLTENHQLWVIRDNQRRIGELWIARRHVGTKPIAFILDIYIEPGE<br>RRQGHARQSMLAAETWARQNDCEEMRLHVFGRNLPARGLYEQLGYDIASLTMAKPLP<br>PA |
| His-TEV-PfIC | ggggaattgtgagcggataacaattcccctgtagaaataattttgtttaactttaataaggagatataccatgggcagcagcc<br>atcaccatcatcaccacgagaacctgtacttccaaagcgatattggaaatcacggagaatttgatagcactgtggacgattt<br>attcaagggaaatacattcctggccgatgcaagatactccagatttgaaattctcttagaacatgtgagtcgaaatcgattca<br>ccattttcatggttctgtacgccgaactgttcaagctgttcagtggagagagtagatttagcgtctttttcgaccaggcagtcgag<br>cttgttttactgcctccccgtgagcaagcctcaataattaatgaccctgtgtttcagatttggaccggtcaggcgtttcgattgac<br>gaatcatttttggttggtaaatcagaaaatacagcagaactcgagaagaccctgtttgagttcccggaaatgttggatcgtgt<br>caaagcacgcagtaaatcggtggtggtacaaaatgacccaccggtttatcgctttgacgcagaccccttataatgcaggc<br>taccccacccagctatgagtttccggaagacgaggctaccagaaaacgtcttgaacgcggcggagtctcctcacgcttttc<br>gctgacgtcgttaaaattgcattgcttcgtatcgaacatacctggccggcgtgcagagaacaaatgaaaagtctgataaagt<br>ccatttgctatctccccgatggaacattccgtagctgttcggcctcccgttacaccggagtgattctgctggcaagccgtgatg<br>actccattctggatctggaagaatccctggtgcacgaagccacgcatcagttgctgtaccatgttgtagaggtctcagcagtg<br>gtcgaccccgatgcttcgcaagaggcctctttctcgctgccgtggtcaggtcaaaagcgggatctgtacggatactttcacgc<br>cttctatgtttatattgcattggtaaaatatatggaacgcgttcgttcacgtagtgacgaagagctccagcgcgcccagaatcgt<br>ttattatttatacttcgaggtctgattcgtgcccttccagacctgcaggggtctgaagacttcacgcctcacggcagacaactgc |

| | |
|---|---|
| | tcgaaaatctcgcagcagaagttcgtgccatggaaagtaaatacgccggtcgattggcggcagcagcgccccttacgact<br>actccgcctgtgcttggggctacggcctaaagccaggatccgaattcgagctcggcgcgcctgcaggtcgacaagcttgc<br>ggccgcataatgcttaagtcgaacagaaagtaatcgtattgtacacggccgcataatcgaaattaatacgactcactatag<br>gggaattgtgagcggataacaattccccatcttagtatattagttaagtataagaaggagatatacatatggcagatctcaatt<br>ggatatcggccggccacgcgatcgctgacgtcggtaccctcgagtctggtaaagaaaccgctgctgcgaaatttgaacgc<br>cagcacatggactcgtctactagcgcagcttaattaacctaggctgctgccaccgctgagcaataactagcataaccccttg<br>gggcctctaaacgggtcttgaggggttttttgctgaaacctcaggcatttgagaagcacacggtcacactgcttccggtagtc<br>aataaaccggtaaaccagcaatagacataagcggctatttaacgaccctgccctgaaccgacgacaagctgacgaccg<br>ggtctccgcaagtggcacttttcggggaaatgtgcgcggaacccctatttgtttatttttctaaatacattcaaatatgtatccgct<br>catgaattaattcttagaaaaactcatcgagcatcaaatgaaactgcaatttattcatatcaggattatcaataccatatttttga<br>aaaagccgtttctgtaatgaaggagaaaactcaccgaggcagttccataggatggcaagatcctggtatcggtctgcgatt<br>ccgactcgtccaacatcaatacaacctattaatttcccctcgtcaaaaataaggttatcaagtgagaaatcaccatgagtga<br>cgactgaatccggtgagaatggcaaaagtttatgcatttctttccagacttgttcaacaggccagccattacgctcgtcatcaa<br>aatcactcgcatcaaccaaaccgttattcattcgtgattgcgcctgagcgagacgaaatacgcggtcgctgttaaaaggac<br>aattacaaacaggaatcgaatgcaaccggcgcaggaacactgccagcgcatcaacaatattttcacctgaatcaggatat<br>tcttctaatacctggaatgctgttttcccggggatcgcagtggtgagtaaccatgcatcatcaggagtacggataaaatgcttg<br>atggtcggaagaggcataaattccgtcagccagtttagtctgaccatctcatctgtaacatcattggcaacgctacctttgcca<br>tgtttcagaaacaactctggcgcatcgggcttcccatacaatcgatagattgtcgcacctgattgcccgacattatcgcgagc<br>ccatttatacccatataaatcagcatccatgttggaatttaatcgcggcctagagcaagacgtttcccgttgaatatggctcata<br>ctcttcctttttcaatattattgaagcatttatcagggttattgtctcatgagcggatacatatttgaatgtatttagaaaaataaaca<br>aataggcatgcagcgctcttccgcttcctcgctcactgactcgctacgctcggtcgttcgactgcggcgagcggtgtcagctc<br>actcaaaagcggtaatacggttatccacagaatcaggggataaagccggaaagaacatgtgagcaaaaagcaaagca<br>ccggaagaagccaacgccgcaggcgtttttccataggctccgcccccctgacgagcatcacaaaaatcgacgctcaagc<br>cagaggtggcgaaacccgacaggactataaagataccaggcgtttccccctggaagctccctcgtgcgctctcctgttccg<br>accctgccgcttaccggatacctgtccgcctttctcccttcgggaagcgtggcgctttctcatagctcacgctgttggtatctcag<br>ttcggtgtaggtcgttcgctccaagctgggctgtgtgcacgaaccccccgttcagcccgaccgctgcgccttatccggtaact<br>atcgtcttgagtccaacccggtaagacacgacttatcgccactggcagcagccattggtaactgatttagaggactttgtcttg<br>aagttatgcacctgttaaggctaaactgaaagaacagattttggtgagtgcggtcctccaacccacttaccttggttcaaaga<br>gttggtagctcagcgaaccttgagaaaaccaccgttggtagcggtggtttttctttatttatgagatgatgaatcaatcggtctat<br>caagtcaacgaacagctattccgttactctagatttcagtgcaatttatctcttcaaatgtagcacctgaagtcagccccatacg<br>atataagttgtaattctcatgttagtcatgccccgcgcccaccggaaggagctgactgggttgaaggctctcaagggcatcg<br>gtcgagatcccggtgcctaatgagtgagctaacttacattaattgcgttgcgctcactgcccgctttccagtcgggaaacctgt<br>cgtgccagctgcattaatgaatcggccaacgcgcggggagaggcggtttgcgtattgggcgccagggtggtttttcttttcac<br>cagtgagacgggcaacagctgattgcccttcaccgcctggccctgagagagttgcagcaagcggtccacgctggtttgcc<br>ccagcaggcgaaaatcctgtttgatggtggttaacggcgggatataacatgagctgtcttcggtatcgtcgtatcccactacc<br>gagatgtccgcaccaacgcgcagcccggactcggtaatggcgcgcattgcgcccagcgccatctgatcgttggcaacca<br>gcatcgcagtgggaacgatgccctcattcagcatttgcatggtttgttgaaaaccggacatggcactccagtcgccttcccgtt<br>ccgctatcggctgaatttgattgcgagtgagatatttatgccagccagccagacgcagacgcgccgagacagaacttaatg<br>ggcccgctaacagcgcgatttgctggtgacccaatgcgaccagatgctccacgcccagtcgcgtaccgtcttcatgggag<br>aaaataatactgttgatgggtgtctggtcagagacatcaagaaataacgccggaacattagtgcaggcagcttccacagc<br>aatggcatcctggtcatccagcggatagttaatgatcagcccactgacgcgttgcgcgagaagattgtgcaccgccgcttta |

| | |
|---|---|
| | caggcttcgacgccgcttcgttctaccatcgacaccaccacgctggcacccagttgatcggcgcgagatttaatcgccgcg<br>acaatttgcgacggcgcgtgcagggccagactggaggtggcaacgccaatcagcaacgactgtttgcccgccagttgttg<br>tgccacgcggttgggaatgtaattcagctccgccatcgccgcttccactttttcccgcgttttcgcagaaacgtggctggcctg<br>gttcaccacgcgggaaacggtctgataagagacaccggcatactctgcgacatcgtataacgttactggtttcacattcacc<br>accctgaattgactctcttccgggcgctatcatgccataccgcgaaaggttttgcgccattcgatggtgtccgggatctcgacg<br>ctctcccttatgcgactcctgcattaggaaattaatacgactcactata |
| Translated sequence | MGSSHHHHHHENLYFQSDIGNHGEFDSTVDDLFKGNTFLADARYSRFEILLEHVSRNR<br>FTIFMVLYAELFKLFSGESRFSVFFDQAVELVLLPPREQASIINDPVFQIWTGQAFRLTNH<br>FLVGKSENTAELEKTLFEFPEMLDRVKARSKSVVVQNDPPVYRFDADPLIMQATPPSYE<br>FPEDEATRKRLERGGVSSRFFADVVKIALLRIEHTWPACREQMKSLIKSICYLPDGTFRS<br>CSASRYTGVILLASRDDSILDLEESLVHEATHQLLYHVVEVSAVVDPDASQEASFSLPWS<br>GQKRDLYGYFHAFYVYIALVKYMERVRSRSDEELQRAQNRLLFILRGLIRALPDLQGSE<br>DFTPHGRQLLENLAAEVRAMESKYAGRLAAAAPLTTTPPVLGATA |
| His-TEV-PflD | actctagatttcagtgcaatttatctcttcaaatgtagcacctgaagtcagccccatacgatataagttgtaattctcatgttagtc<br>atgccccgcgcccaccggaaggagctgactgggttgaaggctctcaagggcatcggtcgagatcccggtgcctaatgagt<br>gagctaacttacattaattgcgttgcgctcactgcccgctttccagtcgggaaacctgtcgtgccagctgcattaatgaatcgg<br>ccaacgcgcggggagaggcggtttgcgtattgggcgccagggtggtttttcttttcaccagtgagacgggcaacagctgatt<br>gcccttcaccgcctggccctgagagagttgcagcaagcggtccacgctggtttgccccagcaggcgaaaatcctgtttgat<br>ggtggttaacggcgggatataacatgagctgtcttcggtatcgtcgtatcccactaccgagatgtccgcaccaacgcgcagc<br>ccggactcggtaatggcgcgcattgcgcccagcgccatctgatcgttggcaaccagcatcgcagtgggaacgatgccctc<br>attcagcatttgcatggtttgttgaaaaccggacatggcactccagtcgccttcccgttccgctatcggctgaatttgattgcga<br>gtgagatatttatgccagccagccagacgcagacgcgccgagacagaacttaatgggcccgctaacagcgcgatttgct<br>ggtgacccaatgcgaccagatgctccacgcccagtcgcgtaccgtcttcatgggagaaaataatactgttgatgggtgtctg<br>gtcagagacatcaagaaataacgccggaacattagtgcaggcagcttccacagcaatggcatcctggtcatccagcgga<br>tagttaatgatcagcccactgacgcgttgcgcgagaagattgtgcaccgccgctttacaggcttcgacgccgcttcgttctac<br>catcgacaccaccacgctggcacccagttgatcggcgcgagatttaatcgccgcgacaatttgcgacggcgcgtgcagg<br>gccagactggaggtggcaacgccaatcagcaacgactgtttgcccgccagttgttgtgccacgcggttgggaatgtaattc<br>agctccgccatcgccgcttccactttttcccgcgttttcgcagaaacgtggctggcctggttcaccacgcgggaaacggtctg<br>ataagagacaccggcatactctgcgacatcgtataacgttactggtttcacattcaccaccctgaattgactctcttccgggcg<br>ctatcatgccataccgcgaaaggttttgcgccattcgatggtgtccgggatctcgacgctctcccttatgcgactcctgcattag<br>gaaattaatacgactcactataggggaattgtgagcggataacaattcccctgtagaaataattttgtttaactttaataagga<br>gatataccatgGGCAGCAGCCATCATCATCATCATCACAGCAGCGGCGAGAACCTGTACT<br>TCCAAAGTatgaccatgttcgacgccagcagatttcctgaggcgggggttgggttagaatatcatttaccacggcaccg<br>cgtagctgatcacgccgtagacccgactttggagcggatcctggcagaggggctgccgtactttgactacttggaatttcag<br>ccgacgcattccattcttgaaccccggttattagaggttggcgaacagacgcctagtctgctgcatagtagctcactcagtctg<br>gggagcgtgggaatcgccatggatcgggaatttctccagatgactcgaagactctgcgaccggacccgaagtccgtggct<br>cgctgagcatatcagctggtcgcggttccacggaggggatacccaacatttcatccttccaacactggctgcagaggtcgct<br>gatacggtagtggctaacgctctggaattgcaagccttaactgcgacgccgttggttttagaaaacgccccacgcctttttctt<br>tagcggatgccccggagcaaagcgaaggcgaattcatcagttccgtggtgcaacggagtggcgcgggatttctgttagac<br>cttgattcagcaatcaccacagcgaaagcattggggtatgactttaaagattatcttcgctctctccctctggatagactgatcg<br>agatacataccggccacccgagacgtgattgggatttgttagctcagctgtttgccgtcagcccggtgagagccgtgacgct |

| | |
|---|---|
| | tgagtgggacatcgcagatagaacggatgacgcgcagcttgaggtactgataagagatataaaacgccttaaaccccgt |
| | gatatgttctggcaagggcgggagccacccccgcacccgatacccctgctctggagccgggcagcctccttaagttaag |
| | agaatcagtttggtttagcgttgggagctctagtttcgcattacgtgatcgtcagtcgggcctgtcccttgatttctgtctgactttatt |
| | accgttgcttaatcattttatgacgccgcacagcctcgaatcggcactgatgctgcccggtgtcttaaatagtccagaacaag |
| | gctctcatctcgcttttctgcaagcgctggttagtcacggaatcgttcaacctgtggcgggaccgcgtgaccgtgtccatcggc |
| | agccactgaaactgtggagccggtgggaggcggccctggagttttatctgagtacgagaacgggactgcagaccccttac |
| | gtatcagttgtcgaactggaagcagaattagaacaaaaagccagccagcaaagacagccgtcgagtttcaaagattatc |
| | attctcatcctttcatcgcactggagaacccattacttgtgcccggggaaacacttgctgaaacaacgctcctcgattctttgtg |
| | cgcacggcgtacaagccgtgcatttagtggcaaaccattgacaccgactcagctgtccctgttgttgtattacacctggggcg |
| | tgacggcaatggaaccaaacggtatgggagactactttcttaaaaaaacaagcccgagtggaggatctttgcaagccacc |
| | gaggtatatgccgtcttaatgaatgtccagggattcgagcgcggcctctaccactactctgtacgaagacatggtctggaact |
| | gctgagtcgtgaggatccccgtacatggattagtgaggcctcaggcggtcagccttgggtgaaagatgcggcagccgtgtt |
| | cgtaagtactgcccgcgtggaacgtttgagttggaagtatgaatttagcagagcgcttcgtgtggccctgatggatgcaggc |
| | catctgtcccagaccttctctctggtcgctactgcattgaacttaggttgctttacaacagccgcattacgcgatgaaatgtttga |
| | gaatcgcctggggctggactatctcgaagaaccggtttttctcttgaacggcgtcggcggttaaagcttgcggccgcataatg |
| | cttaagtcgaacagaaagtaatcgtattgtacacggccgcataatcgaaattaatacgactcactataggggaattgtgagc |
| | ggataacaattccccatcttagtatattagttaagtataagaaggagatatacatatggcagatctcaattggatatcggccgg |
| | ccacgcgatcgctgacgtcggtaccctcgagtctggtaaagaaaccgctgctgcgaaatttgaacgccagcacatggact |
| | cgtctactagcgcagcttaattaacctaggctgctgccaccgctgagcaataactagcataaccccttggggcctctaaacg |
| | ggtcttgaggggttttttgctgaaacctcaggcatttgagaagcacacggtcacactgcttccggtagtcaataaaccggtaa |
| | accagcaatagacataagcggctatttaacgaccctgccctgaaccgacgacaagctgacgaccgggtctccgcaagtg |
| | gcacttttcggggaaatgtgcgcggaacccctatttgtttatttttctaaatacattcaaatatgtatccgctcatgaattaattctta |
| | gaaaaactcatcgagcatcaaatgaaactgcaatttattcatatcaggattatcaataccatatttttgaaaaagccgtttctgt |
| | aatgaaggagaaaactcaccgaggcagttccataggatggcaagatcctggtatcggtctgcgattccgactcgtccaac |
| | atcaatacaacctattaatttcccctcgtcaaaaataaggttatcaagtgagaaatcaccatgagtgacgactgaatccggtg |
| | agaatggcaaaagtttatgcatttctttccagacttgttcaacaggccagccattacgctcgtcatcaaaatcactcgcatcaa |
| | ccaaaccgttattcattcgtgattgcgcctgagcgagacgaaatacgcggtcgctgttaaaaggacaattacaaacaggaa |
| | tcgaatgcaaccggcgcaggaacactgccagcgcatcaacaatattttcacctgaatcaggatattcttctaatacctggaa |
| | tgctgttttcccggggatcgcagtggtgagtaaccatgcatcatcaggagtacggataaaatgcttgatggtcggaagaggc |
| | ataaattccgtcagccagtttagtctgaccatctcatctgtaacatcattggcaacgctacctttgccatgtttcagaaacaactc |
| | tggcgcatcgggcttcccatacaatcgatagattgtcgcacctgattgcccgacattatcgcgagcccatttatacccataa |
| | atcagcatccatgttggaatttaatcgcggcctagagcaagacgtttcccgttgaatatggctcatactcttcctttttcaatattat |
| | tgaagcatttatcagggttattgtctcatgagcggatacatatttgaatgtatttagaaaaataaacaaataggcatgcagcgc |
| | tcttccgcttcctcgctcactgactcgctacgctcggtcgttcgactgcggcgagcggtgtcagctcactcaaaagcggtaata |
| | cggttatccacagaatcaggggataaagccggaaagaacatgtgagcaaaaagcaaagcaccggaagaagccaac |
| | gccgcaggcgtttttccataggctccgcccccctgacgagcatcacaaaaatcgacgctcaagccagaggtggcgaaac |
| | ccgacaggactataaagataccaggcgtttccccctggaagctccctcgtgcgctctcctgttccgaccctgccgcttaccg |
| | gatacctgtccgcctttctcccttcgggaagcgtggcgctttctcatagctcacgctgttggtatctcagttcggtgtaggtcgttc |
| | gctccaagctgggctgtgtgcacgaaccccccgttcagcccgaccgctgcgccttatccggtaactatcgtcttgagtccaa |
| | cccggtaagacacgacttatcgccactggcagcagccattggtaactgatttagaggactttgtcttgaagttatgcacctgtt |
| | aaggctaaactgaaagaacagattttggtgagtgcggtcctccaacccacttaccttggttcaaagagttggtagctcagcg |

| | |
|---|---|
| | aaccttgagaaaaccaccgttggtagcggtggtttttctttatttatgagatgatgaatcaatcggtctatcaagtcaacgaaca<br>gctattccgtt |
| Translated sequence | MGSSHHHHHHSSGENLYFQSMTMFDASRFPEAGVGLEYHLPRHRVADHAVDPTLERIL<br>AEGLPYFDYLEFQPTHSILEPRLLEVGEQTPSLLHSSSLSLGSVGIAMDREFLQMTRRLC<br>DRTRSPWLAEHISWSRFHGGDTQHFILPTLAAEVADTVVANALELQALTATPLVLENAPR<br>LFSLADAPEQSEGEFISSVVQRSGAGFLLDLDSAITTAKALGYDFKDYLRSLPLDRLIEIH<br>TGHPRRDWDLLAQLFAVSPVRAVTLEWDIADRTDDAQLEVLIRDIKRLKPRDMFWQGR<br>EPPPAPDTPALEPGSLLKLRESVWFSVGSSSFALRDRQSGLSLDFCLTLLPLLNHFMTP<br>HSLESALMLPGVLNSPEQGSHLAFLQALVSHGIVQPVAGPRDRVHRQPLKLWSRWEAA<br>LEFYLSTRTGLQTPYVSVVELEAELEQKASQQRQPSSFKDYHSHPFIALENPLLVPGETL<br>AETTLLDSLCARRTSRAFSGKPLTPTQLSLLLYYTWGVTAMEPNGMGDYFLKKTSPSG<br>GSLQATEVYAVLMNVQGFERGLYHYSVRRHGLELLSREDPRTWISEASGGQPWVKDA<br>AAVFVSTARVERLSWKYEFSRALRVALMDAGHLSQTFSLVATALNLGCFTTAALRDEMF<br>ENRLGLDYLEEPVFLLNGVGG |
| PflACD | atgggcagcagccatcaccatcatcaccaccccgcgaaagtgaatcaagaacgtctggcgaccgtggtgggtcgagcg<br>ctgattgataaagatttcgcagaacagttgcataaagacgccgagggtgctgctaagggtattggtgtacatctgagtgcga<br>cagagctctcggcagtaaaaaacatagatgttgccaaacttggaaccgctggtgcaggtattcgcgataaactgggtacg<br>gcggctatcttcgaccagcaacaacaaatacaggcgcggatggactaaAAGGAGATATACAATGACCATG<br>TTCGACGCCAGCAGATTTCCTGAGGCGGGGGTTGGGTTAGAATATCATTTACCACGG<br>CACCGCGTAGCTGATCACGCCGTAGACCCGACTTTGGAGCGGATCCTGGCAGAGG<br>GGCTGCCGTACTTTGACTACTTGGAATTTCAGCCGACGCATTCCATTCTTGAACCCC<br>GGTTATTAGAGGTTGGCGAACAGACGCCTAGTCTGCTGCATAGTAGCTCACTCAGTC<br>TGGGGAGCGTGGGAATCGCCATGGATCGGGAATTTCTCCAGATGACTCGAAGACTC<br>TGCGACCGGACCCGAAGTCCGTGGCTCGCTGAGCATATCAGCTGGTCGCGGTTCC<br>ACGGAGGGGATACCCAACATTTCATCCTTCCAACACTGGCTGCAGAGGTCGCTGAT<br>ACGGTAGTGGCTAACGCTCTGGAATTGCAAGCCTTAACTGCGACGCCGTTGGTTTTA<br>GAAAACGCCCCACGCCTTTTTTCTTTAGCGGATGCCCCGGAGCAAAGCGAAGGCGA<br>ATTCATCAGTTCCGTGGTGCAACGGAGTGGCGCGGGATTTCTGTTAGACCTTGATTC<br>AGCAATCACCACAGCGAAAGCATTGGGGTATGACTTTAAAGATTATCTTCGCTCTCTC<br>CCTCTGGATAGACTGATCGAGATACATACCGGCCACCCGAGACGTGATTGGGATTTG<br>TTAGCTCAGCTGTTTGCCGTCAGCCCGGTGAGAGCCGTGACGCTTGAGTGGGACAT<br>CGCAGATAGAACGGATGACGCGCAGCTTGAGGTACTGATAAGAGATATAAAACGCCT<br>TAAACCCCGTGATATGTTCTGGCAAGGGCGGGAGCCACCCCCCGCACCCGATACCC<br>CTGCTCTGGAGCCGGGCAGCCTCCTTAAGTTAAGAGAATCAGTTTGGTTTAGCGTTG<br>GGAGCTCTAGTTTCGCATTACGTGATCGTCAGTCGGGCCTGTCCCTTGATTTCTGTC<br>TGACTTTATTACCGTTGCTTAATCATTTTATGACGCCGCACAGCCTCGAATCGGCACT<br>GATGCTGCCCGGTGTCTTAAATAGTCCAGAACAAGGCTCTCATCTCGCTTTTCTGCA<br>AGCGCTGGTTAGTCACGGAATCGTTCAACCTGTGGCGGGACCGCGTGACCGTGTC<br>CATCGGCAGCCACTGAAACTGTGGAGCCGGTGGGAGGCGGCCCTGGAGTTTTATC<br>TGAGTACGAGAACGGGACTGCAGACCCCTTACGTATCAGTTGTCGAACTGGAAGCA<br>GAATTAGAACAAAAAGCCAGCCAGCAAAGACAGCCGTCGAGTTTCAAAGATTATCAT |

| | |
|---|---|
| | TCTCATCCTTTCATCGCACTGGAGAACCCATTACTTGTGCCCGGGGAAACACTTGCT GAAACAACGCTCCTCGATTCTTTGTGCGCACGGCGTACAAGCCGTGCATTTAGTGG CAAACCATTGACACCGACTCAGCTGTCCCTGTTGTTGTATTACACCTGGGGCGTGAC GGCAATGGAACCAAACGGTATGGGAGACTACTTTCTTAAAAAAACAAGCCCGAGTG GAGGATCTTTGCAAGCCACCGAGGTATATGCCGTCTTAATGAATGTCCAGGGATTCG AGCGCGGCCTCTACCACTACTCTGTACGAAGACATGGTCTGGAACTGCTGAGTCGT GAGGATCCCCGTACATGGATTAGTGAGGCCTCAGGCGGTCAGCCTTGGGTGAAAGA TGCGGCAGCCGTGTTCGTAAGTACTGCCCGCGTGGAACGTTTGAGTTGGAAGTATG AATTTAGCAGAGCGCTTCGTGTGGCCCTGATGGATGCAGGCCATCTGTCCCAGACC TTCTCTCTGGTCGCTACTGCATTGAACTTAGGTTGCTTTACAACAGCCGCATTACGC GATGAAATGTTTGAGAATCGCCTGGGGCTGGACTATCTCGAAGAACCGGTTTTTCTC TTGAACGGCGTCGGCGGTTAAAGCCAGGATCCGAATTCGAGCTCGGCGCGCCTGC AGGTCGACAAGCTTGCGGCCGCATAATGCTTAAGTCGAACAGAAAGTAATCGTATTG TACACGGCCGCATAATCGAAATTAATACGACTCACTATAGGGGAATTGTGAGCGGATA ACAATTCCCCATCTTAGTATATTAGTTAAGTATAAGAAGGAGATATACATATGGATATTG GAAATCACGGAGAATTTGATAGCACTGTGGACGATTTATTCAAGGGAAATACATTCCT GGCCGATGCAAGATACTCCAGATTTGAAATTCTCTTAGAACATGTGAGTCGAAATCGA TTCACCATTTTCATGGTTCTGTACGCCGAACTGTTCAAGCTGTTCAGTGGAGAGAGT AGATTTAGCGTCTTTTTCGACCAGGCAGTCGAGCTTGTTTTACTGCCTCCCCGTGAG CAAGCCTCAATAATTAATGACCCTGTGTTTCAGATTTGGACCGGTCAGGCGTTTCGAT TGACGAATCATTTTTTGGTTGGTAAATCAGAAAATACAGCAGAACTCGAGAAGACCCT GTTTGAGTTCCCGGAAATGTTGGATCGTGTCAAAGCACGCAGTAAATCGGTGGTGG TACAAAATGACCCACCGGTTTATCGCTTTGACGCAGACCCCCTTATAATGCAGGCTA CCCCACCCAGCTATGAGTTTCCGGAAGACGAGGCTACCAGAAAACGTCTTGAACGC GGCGGAGTCTCCTCACGCTTTTTCGCTGACGTCGTTAAAATTGCATTGCTTCGTATC GAACATACCTGGCCGGCGTGCAGAGAACAAATGAAAAGTCTGATAAAGTCCATTTGC TATCTCCCCGATGGAACATTCCGTAGCTGTTCGGCCTCCCGTTACACCGGAGTGATT CTGCTGGCAAGCCGTGATGACTCCATTCTGGATCTGGAAGAATCCCTGGTGCACGA AGCCACGCATCAGTTGCTGTACCATGTTGTAGAGGTCTCAGCAGTGGTCGACCCCG ATGCTTCGCAAGAGGCCTCTTTCTCGCTGCCGTGGTCAGGTCAAAAGCGGGATCTG TACGGATACTTTCACGCCTTCTATGTTTATATTGCATTGGTAAAATATATGGAACGCGTT CGTTCACGTAGTGACGAAGAGCTCCAGCGCGCCCAGAATCGTTTATTATTTATACTTC GAGGTCTGATTCGTGCCCTTCCAGACCTGCAGGGGTCTGAAGACTTCACGCCTCAC GGCAGACAACTGCTCGAAAATCTCGCAGCAGAAGTTCGTGCCATGGAAAGTAAATA CGCCGGTCGATTGGCGGCAGCAGCGCCCCTTACGACTACTCCGCCTGTGCTTGGG GCTACGGCCTAA |
| Color code | PflA linker PflD linker PflC |
| PflABD | atgggcagcagccatcaccatcatcaccaccccgcgaaagtgaatcaagaacgtctggcgaccgtggtgggtcgagcg ctgattgataaagatttcgcagaacagttgcataaagacgccgagggtgctgctaagggtattggtgtacatctgagtgcga cagagctctcggcagtaaaaaacatagatgttgccaaacttggaaccgctggtgcaggtattcgcgataaactgggtacg gcggctatcttcgaccagcaacaacaaatacaggcgcggatggactaaaagaaggagatataccatgagtgttacactg |

| | |
|---|---|
| | atcccgatggatgatgcggattttgttacttttaccgaacgggcagtggatgaatacgctcaaggtatggttgccagcggtga<br>atggagccaggcggaagcgtcacatcgctcagccacagcattcgctcaactcctggataaagggcgtctgacggaaaat<br>caccagttatgggtcattagagataatcagcgtcgaatcggagaattatggatcgcgcgccggcatgttgggactaaaccg<br>atagctttcatactggatatttatatagaaccgggtgaacgccgccaaggtcatgcccgtcagtctatgttggcggcagaaac<br>atgggcacgacagaatgactgcgaagaaatgcgactgcatgtatttggccgtaatctgcctgcgcgtggcctttatgaacag<br>ttaggttatgacatagcaagtcttactatggctaagccactgccacccgcataaaaggagatatacaatgaccatgttcgac<br>gccagcagatttcctgaggcgggggttgggttagaatatcatttaccacggcaccgcgtagctgatcacgccgtagacccg<br>actttggagcggatcctggcagaggggctgccgtactttgactacttggaatttcagccgacgcattccattcttgaaccccgg<br>ttattagaggttggcgaacagacgcctagtctgctgcatagtagctcactcagtctggggagcgtgggaatcgccatggatc<br>gggaatttctccagatgactcgaagactctgcgaccggacccgaagtccgtggctcgctgagcatatcagctggtcgcggtt<br>ccacggaggggatacccaacatttcatccttccaacactggctgcagaggtcgctgatacggtagtggctaacgctctgga<br>attgcaagccttaactgcgacgccgttggttttagaaaacgccccacgcctttttctttagcggatgccccggagcaaagcg<br>aaggcgaattcatcagttccgtggtgcaacggagtggcgcgggatttctgttagaccttgattcagcaatcaccacagcgaa<br>agcattggggtatgactttaaagattatcttcgctctctccctctggatagactgatcgagatacataccggccacccgagac<br>gtgattgggatttgttagctcagctgtttgccgtcagcccggtgagagccgtgacgcttgagtgggacatcgcagatagaac<br>ggatgacgcgcagcttgaggtactgataagagatataaaacgccttaaaccccgtgatatgttctggcaagggcgggagc<br>caccccccgcacccgataccccctgctctggagccgggcagcctccttaagttaagagaatcagtttggtttagcgttgggag<br>ctctagtttcgcattacgtgatcgtcagtcgggcctgtcccttgatttctgtctgactttattaccgttgcttaatcattttatgacgccg<br>cacagcctcgaatcggcactgatgctgcccggtgtcttaaatagtccagaacaaggctctcatctcgcttttctgcaagcgct<br>ggttagtcacggaatcgttcaacctgtggcgggaccgcgtgaccgtgtccatcggcagccactgaaactgtggagccggt<br>gggaggcggccctggagttttatctgagtacgagaacgggactgcagaccccttacgtatcagttgtcgaactggaagcag<br>aattagaacaaaaagccagccagcaaagacagccgtcgagtttcaaagattatcattctcatcctttcatcgcactggaga<br>acccattacttgtgcccggggaaacacttgctgaaacaacgctcctcgattctttgtgcgcacggcgtacaagccgtgcattt<br>agtggcaaaccattgacaccgactcagctgtccctgttgttgtattacacctggggcgtgacggcaatggaaccaaacggt<br>atgggagactactttcttaaaaaaacaagcccgagtggaggatctttgcaagccaccgaggtatatgccgtcttaatgaatg<br>tccagggattcgagcgcggcctctaccactactctgtacgaagacatggtctggaactgctgagtcgtgaggatccccgtac<br>atggattagtgaggcctcaggcggtcagccttgggtgaaagatgcggcagccgtgttcgtaagtactgcccgcgtggaacg<br>tttgagttggaagtatgaatttagcagagcgcttcgtgtggccctgatggatgcaggccatctgtcccagaccttctctctggtc<br>gctactgcattgaacttaggttgctttacaacagccgcattacgcgatgaaatgtttgagaatcgcctggggctggactatctc<br>gaagaaccggtttttctcttgaacggcgtcggcggttaa |
| Color code | PflA linker PflB linker PflD |
| His6-PosA | ATGGGCAGCAGCCATCACCATCATCACCACTCTACTAAAGTAAATCAGGAGCGACTC<br>GCGAGCGTCGTGGGTAGAGCCCTGCTCGACAAGGATTTTGCCGCCCAGCTCCACC<br>AGGATCCGGAGGCTGCGGCTAAAGGTATAGGCGTGCATTTAAGTGCGACGGAGGTG<br>GGTGCGGTCAAAAGTATCGACACTGCTAAACTCACATCAGCAGGCTCCGCAATCCG<br>CGATAAGATTGGAGTAGCTGCGATTTTCGACACTCAGCAGCAACAAGCTCGAATGGA<br>CTGA |
| Translated sequence | MGSSHHHHHHSTKVNQERLASVVGRALLDKDFAAQLHQDPEAAAK<br>GIGVHLSATEVGAVK SIDTAK LTSAGSAIRDK IGVAAIFDTQQQQARMD |
| His-TEV-PosC | ggggaattgtgagcggataacaattcccctgtagaaataattttgtttaactttaataaggagatataccatgggcagcagcc<br>atcaccatcatcaccacgagaacctgtacttccaaagcgattcggtcaactcaaagcaagctgaactgagtatcgatgaac |

| | |
|---|---|
| | tgttcaaaggaacaacttttctcgcggactcagagccgcagcgttttctgcaattacaagaacaggttgccaagaacagata<br>taccatgttcgttgccttgtacgctgaattaagtaaactgtttgcggcaaattctgctctgcgtgttttctttaagcaagcactcgac<br>atcattctttctccggagtcagtgcgaaatccactggttgcacatcccgtatttcagatctggtccgtgcttacctttcgtgatgtga<br>actatttgctgacgggcaaaaccctggatgattcagaggtgatcgcacgcctgttggaatttccgcaggtactgcaaagaat<br>cgaagccgcccaacaggcgcgtccagatgagcagtgccccctgtttatagattcgacgtcgaccctctgattactcaggt<br>gacaccaccgtcatacgattaccctccggatgaagccacacggcgccagttagaaagagccggctactccaaggcatttt<br>ttcgggatgttatgcaactggcactccagagaattaaacacacgtggcccgcatgccacgaacagtggcaagttctggtta<br>aagcagtgtgctatctccccgatggctcattccgtagctgttctgccagcagatatacaggtgtgattctgttgagctctcgcga<br>caacagtattcttgatttagaagaatcactggtccacgaggcaacacatcagttattatataatgtagtagaagtggcagccg<br>tggtagagccgcaggcaagccgggaagttttgtataccttaccatggagcgggcaacagcgcgatctgtacggttattttcat<br>gcattttacgtttatatcgctttagttaagtacttggaacgtgttcgcgatcgaccggctcaggagatgcgtcgtgcggagcaac<br>gcctcctgtttatccttcgcggtttatcaaaagcccaagcggattttgctgcagcgcccggcttcaccgctcagggccgtgagt<br>tgttaggtaacctgctgcaggaagttcatcgccttgaacgacgacacgcaggttccctgagccgtggggaaggtgcctccg<br>cggtagccacagtgcttcacatgactgcatgaagccaggatccgaattcgagctcggcgcgcctgcaggtcgacaagctt<br>gcggccgcataatgcttaagtcgaacagaaagtaatcgtattgtacacggccgcataatcgaaattaatacgactcactata<br>ggggaattgtgagcggataacaattccccatcttagtatattagttaagtataagaaggagatatacatatggcagatctcaa<br>ttggatatcggccggccacgcgatcgctgacgtcggtaccctcgagtctggtaaagaaaccgctgctgcgaaatttgaacg<br>ccagcacatggactcgtctactagcgcagcttaattaacctaggctgctgccaccgctgagcaataactagcataacccctt<br>ggggcctctaaacgggtcttgaggggttttttgctgaaacctcaggcatttgagaagcacacggtcacactgcttccggtagt<br>caataaaccggtaaaccagcaatagacataagcggctatttaacgaccctgccctgaaccgacgacaagctgacgacc<br>gggtctccgcaagtggcacttttcggggaaatgtgcgcggaacccctatttgtttatttttctaaatacattcaaatatgtatccgc<br>tcatgaattaattcttagaaaaactcatcgagcatcaaatgaaactgcaatttattcatatcaggattatcaataccatatttttga<br>aaaagccgtttctgtaatgaaggagaaaactcaccgaggcagttccataggatggcaagatcctggtatcggtctgcgatt<br>ccgactcgtccaacatcaatacaacctattaatttcccctcgtcaaaaataaggttatcaagtgagaaatcaccatgagtga<br>cgactgaatccggtgagaatggcaaaagtttatgcatttctttccagacttgttcaacaggccagccattacgctcgtcatcaa<br>aatcactcgcatcaaccaaaccgttattcattcgtgattgcgcctgagcgagacgaaatacgcggtcgctgttaaaaggac<br>aattacaaacaggaatcgaatgcaaccggcgcaggaacactgccagcgcatcaacaatattttcacctgaatcaggatat<br>tcttctaatacctggaatgctgttttcccggggatcgcagtggtgagtaaccatgcatcatcaggagtacggataaaatgcttg<br>atggtcggaagaggcataaattccgtcagccagtttagtctgaccatctcatctgtaacatcattggcaacgctacctttgcca<br>tgtttcagaaacaactctggcgcatcgggcttcccatacaatcgatagattgtcgcacctgattgcccgacattatcgcgagc<br>ccatttatacccatataaatcagcatccatgttggaatttaatcgcggcctagagcaagacgtttcccgttgaatatggctcata<br>ctcttcctttttcaatattattgaagcatttatcagggttattgtctcatgagcggatacatatttgaatgtatttagaaaaataaaca<br>aataggcatgcagcgctcttccgcttcctcgctcactgactcgctacgctcggtcgttcgactgcggcgagcggtgtcagctc<br>actcaaaagcggtaatacggttatccacagaatcaggggataaagccggaaagaacatgtgagcaaaaagcaaagca<br>ccggaagaagccaacgccgcaggcgtttttccataggctccgcccccctgacgagcatcacaaaaatcgacgctcaagc<br>cagaggtggcgaaacccgacaggactataaagataccaggcgtttccccctggaagctccctcgtgcgctctcctgttccg<br>accctgccgcttaccggatacctgtccgcctttctcccttcgggaagcgtggcgctttctcatagctcacgctgttggtatctcag<br>ttcggtgtaggtcgttcgctccaagctgggctgtgtgcacgaaccccccgttcagcccgaccgctgcgccttatccggtaact<br>atcgtcttgagtccaacccggtaagacacgacttatcgccactggcagcagccattggtaactgatttagaggactttgtcttg<br>aagttatgcacctgttaaggctaaactgaaagaacagattttggtgagtgcggtcctccaacccacttaccttggttcaaaga<br>gttggtagctcagcgaaccttgagaaaaccaccgttggtagcggtggtttttctttatttatgagatgatgaatcaatcggtctat |

| | |
|---|---|
| | caagtcaacgaacagctattccgttactctagatttcagtgcaatttatctcttcaaatgtagcacctgaagtcagccccatacg<br>atataagttgtaattctcatgttagtcatgccccgcgcccaccggaaggagctgactgggttgaaggctctcaagggcatcg<br>gtcgagatcccggtgcctaatgagtgagctaacttacattaattgcgttgcgctcactgcccgctttccagtcgggaaacctgt<br>cgtgccagctgcattaatgaatcggccaacgcgcggggagaggcggtttgcgtattgggcgccagggtggtttttcttttcac<br>cagtgagacgggcaacagctgattgcccttcaccgcctggccctgagagagttgcagcaagcggtccacgctggtttgcc<br>ccagcaggcgaaaatcctgtttgatggtggttaacggcgggatataacatgagctgtcttcggtatcgtcgtatcccactacc<br>gagatgtccgcaccaacgcgcagcccggactcggtaatggcgcgcattgcgcccagcgccatctgatcgttggcaacca<br>gcatcgcagtgggaacgatgccctcattcagcatttgcatggtttgttgaaaaccggacatggcactccagtcgccttcccgtt<br>ccgctatcggctgaatttgattgcgagtgagatatttatgccagccagccagacgcagacgcgccgagacagaacttaatg<br>ggcccgctaacagcgcgatttgctggtgacccaatgcgaccagatgctccacgcccagtcgcgtaccgtcttcatgggag<br>aaaataatactgttgatgggtgtctggtcagagacatcaagaaataacgccggaacattagtgcaggcagcttccacagc<br>aatggcatcctggtcatccagcggatagttaatgatcagcccactgacgcgttgcgcgagaagattgtgcaccgccgcttta<br>caggcttcgacgccgcttcgttctaccatcgacaccaccacgctggcacccagttgatcggcgcgagatttaatcgccgcg<br>acaatttgcgacggcgcgtgcagggccagactggaggtggcaacgccaatcagcaacgactgtttgcccgccagttgttg<br>tgccacgcggttgggaatgtaattcagctccgccatcgccgcttccactttttcccgcgttttcgcagaaacgtggctggcctg<br>gttcaccacgcgggaaacggtctgataagagacaccggcatactctgcgacatcgtataacgttactggtttcacattcacc<br>accctgaattgactctcttccgggcgctatcatgccataccgcgaaaggttttgcgccattcgatggtgtccgggatctcgacg<br>ctctcccttatgcgactcctgcattaggaaattaatacgactcactata |
| Translated sequence | MGSSHHHHHHENLYFQSDSVNSKQAELSIDELFKGTTFLADSEPQRFLQLQEQVAKNR<br>YTMFVALYAELSKLFAANSALRVFFKQALDIILSPESVRNPLVAHPVFQIWSVLTFRDVNY<br>LLTGKTLDDSEVIARLLEFPQVLQRIEAAQQARPDEQCPPVYRFDVDPLITQVTPPSYDY<br>PPDEATRRQLERAGYSKAFFRDVMQLALQRIKHTWPACHEQWQVLVKAVCYLPDGSF<br>RSCSASRYTGVILLSSRDNSILDLEESLVHEATHQLLYNVVEVAAVVEPQASREVLYTLP<br>WSGQQRDLYGYFHAFYVYIALVKYLERVRDRPAQEMRRAEQRLLFILRGLSKAQADFAA<br>APGFTAQGRELLGNLLQEVHRLERRHAGSLSRGEGASAVATVLHMTA |
| PosABCD | **TAATACGACTCACTATAGGGGAATTGTGAGCGGATAACAATTCCCCTGTAGAAATAA<br>TTTTGTTTAACTTTAATAAGGAGATATACC**ATGGGCAGCAGCCATCACCATCATCACC<br>ACTCTACTAAAGTAAATCAGGAGCGACTCGCGAGCGTCGTGGGTAGAGCCCTGCTC<br>GACAAGGATTTTGCCGCCCAGCTCCACCAGGATCCGGAGGCTGCGGCTAAAGGTAT<br>AGGCGTGCATTTAAGTGCGACGGAGGTGGGTGCGGTCAAAAGTATCGACACTGCTA<br>AACTCACATCAGCAGGCTCCGCAATCCGCGATAAGATTGGAGTAGCTGCGATTTTCG<br>ACACTCAGCAGCAACAAGCTCGAATGGACTGAAAGAAGGAGATATACCATGAGCATC<br>AGCTTAGTTTCTATGCAGGAAGCTGACTTTGAACGTTTTGCACGCCGCGCACTTCAA<br>GAATATGCTGCCGGAATGGTTGCAGCGGGCGAATGGCCACCGGAACAGGCCTCTTT<br>TCAGGCGGCCGAGGTTTTTCGACAGCTGCTTCCTCAAGGACGTTTGAGCCCGCATA<br>ATCATTTGTGGCGTGCATTACACGCTGGACGTGCCGTGGGAGAATTGTGGATCGCA<br>GAACGTCAGGCAGGCAGTCGCCGGATAGCCTTTATTCTCGATCTGTATATTGAACCC<br>ATTGAACGTCGTCAGGGGTACGCCCGTCAGACGCTGCTGGCTGGTGAAAGTCTGG<br>CGCGCGAGTGGGGTTGTGACGAAATGCGTCTTCATGTCTTTGGTCGCAACTTAGAA<br>GCACGCCGCTTGTATGAATTACTTGGCTATGAAGTCGCATCATTAACGATGTCCAAGC<br>CGTTGCCGTAAAAGGAGATATACAATGCAGGACTTTGATGCATCCCGTTTCCCCGAT |

| | |
|---|---|
| | GCAGGGGTCGGGTTAGAGTATCATCTCCCCCCGGGTGCGGGCAGCGGTGCTGTTG<br>GGTTAGACCCGAGCCTTGAGCGCATCCAAGCTGAGGGCCTGCCGTACTTCGACTAC<br>TTAGAATTTCAACCAACACATTGTATCCTTCAACCGCGCCTTAACCAATTGGGTGGAC<br>AGGTGCCGGTGCTGCTTCACAGTTCATCACTGTCTCTTGGTTCGGTGGGAATTGAG<br>ATGGACCGCGAATTCCTGCACATGACGAGACGGCTGTGCCAGCGCACCGCAAGCC<br>CTTGGTTAGCAGAGCATATTTCTTGGAGCAGATTTCAGGGGGGGGATACTCAGCATT<br>TCATTCTGCCGTCCCTGGCCAGAGAAGTAGCAGACACAGTAGTTGCCAATGCCCTT<br>GAGTTGCAAGCATTTACTGGCACCCCCCTGGTGCTTGAAAATGCCCCCCGCTTATTT<br>GCAATGGATATTGGTCAGGAGTTATCCGAAGGCGGATTCATCTGCTCGGTGGTTGAA<br>CGTTCACACTCCGGGTTTCTCCTTGACTTAGATTCAGCAATCTGCACAGCACGCGCC<br>CTGGGATACGATCTTCGAGAATATTTGCGTTCATTGCCTCTGAAACGCCTGATCGAG<br>GTTCATGTTGGCGACCCGCGTCGCGATTGGGACATACTCCGACAGCTGTTTGGTCA<br>CGCACCCCTGAAGGCAGTTACAATAGAATGGGATATCGCCGACCGAGCGGGAGATC<br>CGGAACTGGCAGCTCTGATTCGGGAAATTAAGGCCTTACGCCCGGAACCCCTTTTTT<br>GGCAGGCGGGCGCGCTCGCGAAGCCTGCACCGCCAGCAGCGGGCCCCCTGGATG<br>GCCAGACCCTGCTTCAGCTGCGGCCAGGGACTTGGTTGGGTATTGATCGCGATCGA<br>TTTACTCTGCGGGACCGCCGCCGAGCGCTGGATCTGGAATTAGCCCTGCAGTTCCT<br>GCCCCTTGTACGCCATTTCCTTTGCCCGCGGACGCTCGACTCTGCCCTGCTGCTGC<br>CCGGCGCCTTAGAGAATCCTTCTCTCGCTCTCCTCCAGGACCTGATTGCTCGTGGT<br>CTCCTGCAACCCGTGGATGGCCAGGGGTCCGGCCCATCGGAACCGGAAGCTGCC<br>GGAGAAATATGGTCGCATTGGCAGGTTGCTCTCGATTTTTATTTGGGTACCCGTACG<br>CAGACGGCAACTCCATATATCTCGGTTGCTCAGATGGAGGAGCAACTGGCCGGCAA<br>AGCACTGCAGCAACGTCAGCCTTCGGCTTTTAAAGACTATTTTAGTCATCCGTTCCA<br>AGCCTTAGATAACCCACTGCTCGCATCGCCGTCCCTGTTTGAACAGCCACAATTGCT<br>GGAGGTCTTGGCCCGTCGGCAGACTTGTCGAACGTTTGACGGGGGTCCGGTCACT<br>GCCCAGCAGTTATCCGCGCTCTTGTATTACACATGGGGAGCCAGCCAGGTGCGTCG<br>AAACCCTATGGGTGATTTTTTTTTAAAAAAAACCAGTGCCTCAGGCGGCTCGTTGCAT<br>GCGATCGAGGTTTATCCAATCCTGCTCAATGTTCAAGGTCTTGAAGCAGGCTTGTAC<br>CATTACAGTGTGCGCCGTCATGGGCTGGAGTTGCTGTCGCGGGAAGACCCGCGTT<br>CGTGGATAGGTGCGGCTTGTGGTGATCAGAGTTGGATCGAACAGGCTTCTGTGTTG<br>TTTCTGAGCACTGCGTGCCTCCAGCGTCTTGCATGGAAGTATCCTGGTTCCCGTGC<br>CCTGCGCGTTGCCTTAATGGACTGTGGGCACTTGTCCCAATCTTTTGCTCTGGTAGC<br>AACAGCGTTAGGTTTGCGTTCGTGTACTACGGCGGCCCTCCGTGATGAGACCTTTG<br>AGACGCGTCTCGGTTTGGATTATCTTCGGGAACCTGTATTCCTGCTGAATGGTGCAG<br>GAGGCTGA<br>AAGCCAGGATCCGAATTCGAGCTCGGCGCGCCTGCAGGTCGACAAGCTTGCGGCC<br>GCATAATGCTTAAGTCGAACAGAAAGTAATCGTATTGTACACGGCCGCATAATCGAAA<br>T**TAATACGACTCACTATAGGGGAATTGTGAGCGGATAACAATTCCCCATCTTAGTAT**<br>**ATTAGTTAAGTATAAGAAGGAGATATACAT**ATGGATTCGGTCAACTCAAAGCAAGCT<br>GAACTGAGTATCGATGAACTGTTCAAAGGAACAACTTTTCTCGCGGACTCAGAGCC<br>GCAGCGTTTTCTGCAATTACAAGAACAGGTTGCCAAGAACAGATATACCATGTTCGTT |

| | |
|---|---|
| | GCCTTGTACGCTGAATTAAGTAAACTGTTTGCGGCAAATTCTGCTCTGCGTGTTTTCT<br>TTAAGCAAGCACTCGACATCATTCTTTCTCCGGAGTCAGTGCGAAATCCACTGGTTG<br>CACATCCCGTATTTCAGATCTGGTCCGTGCTTACCTTTCGTGATGTGAACTATTTGCT<br>GACGGGCAAAACCCTGGATGATTCAGAGGTGATCGCACGCCTGTTGGAATTTCCGC<br>AGGTACTGCAAAGAATCGAAGCCGCCCAACAGGCGCGTCCAGATGAGCAGTGCCC<br>CCCTGTTTATAGATTCGACGTCGACCCTCTGATTACTCAGGTGACACCACCGTCATA<br>CGATTACCCTCCGGATGAAGCCACACGGCGCCAGTTAGAAAGAGCCGGCTACTCCA<br>AGGCATTTTTTCGGGATGTTATGCAACTGGCACTCCAGAGAATTAAACACACGTGGC<br>CCGCATGCCACGAACAGTGGCAAGTTCTGGTTAAAGCAGTGTGCTATCTCCCCGAT<br>GGCTCATTCCGTAGCTGTTCTGCCAGCAGATATACAGGTGTGATTCTGTTGAGCTCT<br>CGCGACAACAGTATTCTTGATTTAGAAGAATCACTGGTCCACGAGGCAACACATCAG<br>TTATTATATAATGTAGTAGAAGTGGCAGCCGTGGTAGAGCCGCAGGCAAGCCGGGAA<br>GTTTTGTATACCTTACCATGGAGCGGGCAACAGCGCGATCTGTACGGTTATTTTCATG<br>CATTTTACGTTTATATCGCTTTAGTTAAGTACTTGGAACGTGTTCGCGATCGACCGGC<br>TCAGGAGATGCGTCGTGCGGAGCAACGCCTCCTGTTTATCCTTCGCGGTTTATCAAA<br>AGCCCAAGCGGATTTTGCTGCAGCGCCCGGCTTCACCGCTCAGGGCCGTGAGTTG<br>TTAGGTAACCTGCTGCAGGAAGTTCATCGCCTTGAACGACGACACGCAGGTTCCCT<br>GAGCCGTGGGGAAGGTGCCTCCGCGGTAGCCACAGTGCTTCACATGACTGCATGA |
| Color code | PosA linker PosB linker PosD linker PosC (**bold sequence includes T7 promoter, lac operator, and RBS site**) |

**Table S2:** Accession IDs for proteins.

Proteins in the *pfl* cluster have been removed during database consolidation efforts and are no longer in the NCBI database.

| Protein | NCBI ID or gene bank ID |
|---|---|
| Pfl BGC | Biosample SAMN07703430 |
| PosC | WP_060837885.1 |
| PosD | WP_060837886.1 |
| PosA | WP_016965124.1 |

**Table S3**: $^{1}$H and $^{13}$C chemical shifts of the LysC-digested PosAC peptide in d6-DMSO at 25 $^{o}$C.

| number | AA | NH<br>N | αH<br>Cα | βH<br>Cβ | γH<br>Cγ | others |
|---|---|---|---|---|---|---|
| 1 | I | | 3.36<br>58.4 | 1.72<br>37.6 | 1.45, 1.10<br>24.3<br>0.84(t), 11.8 | 0.88 (d), 15.5 |
| 2 | G | 8.35<br>108.2 | 3.86, 3.78<br>42.3 | | | |
| 3 | V | 7.894<br>118.0 | 4.21<br>58.2 | 1.95<br>31.1 | 0.85, 19.6<br>0.82, 18.5 | |
| 4 | A | 8.16<br>122.7 | 4.27<br>48.5 | 1.19<br>18.4 | | |
| 5 | A | 7.892<br>116.7 | 4.27<br>48.5 | 1.14<br>18.5 | | |
| 6 | I | 7.67<br>113.0 | 4.10<br>57.2 | 1.64<br>37.3 | Hγ1:<br>1.29, 0.97<br>24.4<br>0.75(t), 11.5 | 0.70 (d), 15.7 |
| 7 | F | 7.96<br>117.8 | 4.58<br>53.6 | 2.96, 2.76<br>38.3 | 138.0 | 7.20, 128.5<br>7.20, 129.7<br>7.15, 126.6 |
| 8 | D | 8.18<br>117.2 | 4.65<br>50.7 | 2.56, 2.37<br>40.3 | | |
| 9 | T | 7.96<br>117.8 | 4.11<br>57.3 | 4.23<br>66.0 | 1.06<br>20.8 | OH: 4.86 |
| 10 | Q | 8.58 | 4.18<br>59.0 | 4.14, 69.0<br>OH: 5.39 or 5.56 | 2.33<br>39.7 | εNH$_2$: 7.51, 6.73<br>εN: 109.5 |
| 11 | Q | 7.87<br>114.0 | 4.19<br>59.5 | 4.17,69.0<br>OH: 5.56 or 5.39 | 2.28,2.18<br>39.7 | εNH$_2$: 7.62, 6.78<br>εN: 109.8 |
| 12 | Q | 7.85<br>114.4 | 4.25<br>58.0 | 4.11,68.8<br>OH: 5.25 | 2.33,2.23<br>39.7 | εNH$_2$: 7.34, 6.86<br>εN: 109.9 |
| 13 | Q | 7.87<br>114.0 | 4.18<br>59.5 | 4.12,68.8<br>OH: 5.14 | 2.25<br>39.7 | εNH$_2$: 7.28, 6.82<br>εN: 110.0<br>C-terminal NH$_2$:<br>7.19, 7.16<br>N: 106.3 |

**Table S4**: $^{1}$H and $^{13}$C chemical shifts of residues 5 to 8 of the LysC-digested PflAD peptide in d6-DMSO at 25 $^{o}$C.

| number | AA | NH | αH/C | βH/C | γH/C | δH/C and others |
|---|---|---|---|---|---|---|
| 5 | A | 8.06 | 4.37 | 1.22<br>18.6 | | |
| 6 | I | 7.91 | 4.26 | 1.83<br>36.7 | 1.48, 1.11<br>24.7 | 1.19 (d)<br>0.89 (g2) |
| 7 | F | 9.79 (s) | N/A | 6.96 (s)<br>127.9 | | 7.56 (d), 129.9<br>7.37 (t), 129.0<br>7.34 (t), 129.1 |
| 8 | D | 8.28 | 4.63<br>50.4 | 2.82, 2.64<br>36.1 | | |
| 9 | Q | 7.80 | 4.22 | 1.90, 1.82<br>28.0 | 2.16<br>30.7 | ε$NH_2$: 7.25, 6.80 |

The $^{1}$H and $^{13}$C chemical shifts are referenced to 2.50 ppm and 40.5 ppm of the DMSO-d6, respectively.

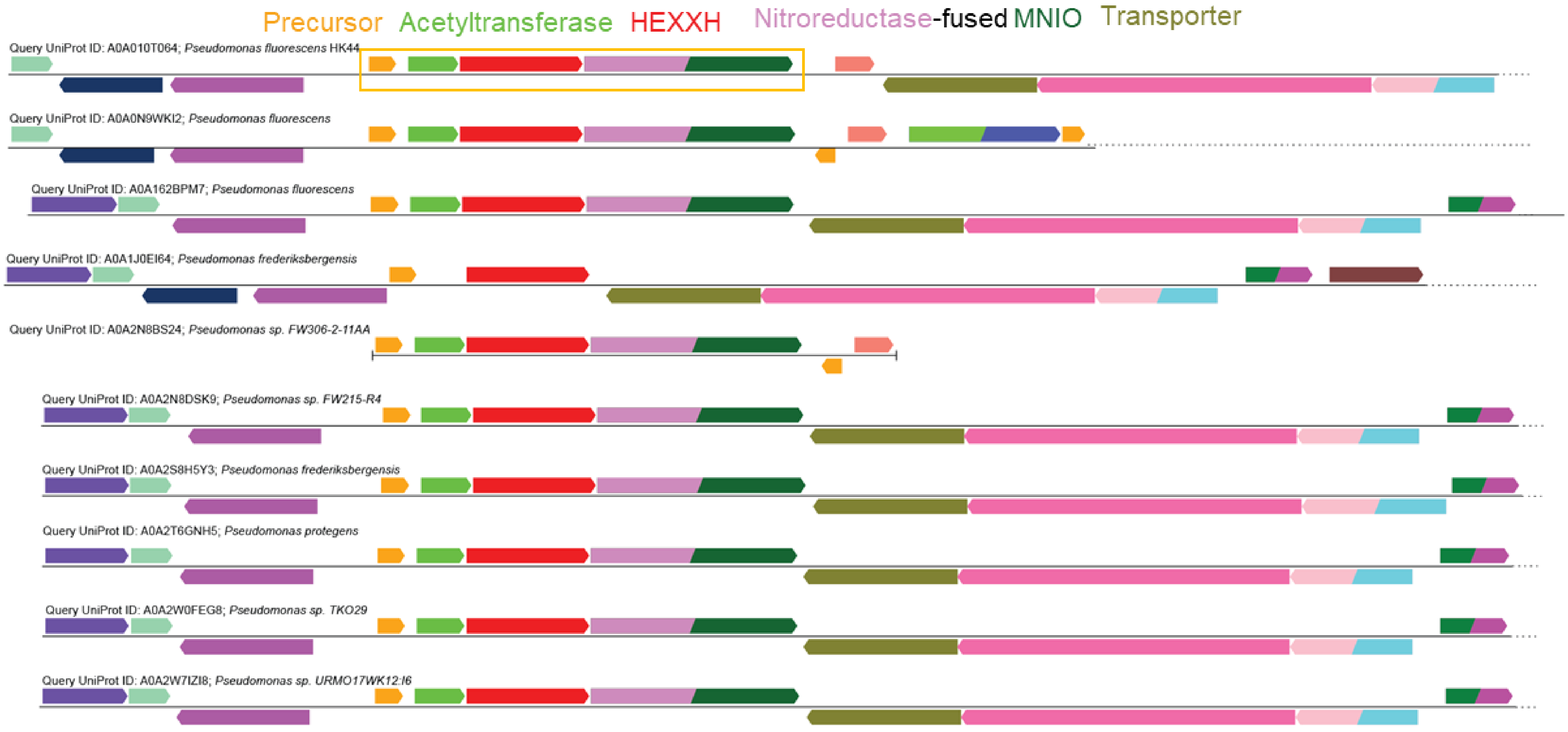
Precursor
Acetyltransferase
HEXXH
Nitroreductase-fused MNIO
Transporter
Query UniProt ID: A0A010T064; Pseudomonas fluorescens HK44
Query UniProt ID: A0A0N9WKI2; Pseudomonas fluorescens
Query UniProt ID: A0A162BPM7; Pseudomonas fluorescens
Query UniProt ID: A0A1J0EI64; Pseudomonas frederiksbergensis
Query UniProt ID: A0A2N8BS24; Pseudomonas sp. FW306-2-11AA
Query UniProt ID: A0A2N8DSK9; Pseudomonas sp. FW215-R4
Query UniProt ID: A0A2S8H5Y3; Pseudomonas frederiksbergensis
Query UniProt ID: A0A2T6GNH5; Pseudomonas protegens
Query UniProt ID: A0A2W0FEG8; Pseudomonas sp. TKO29
Query UniProt ID: A0A2W7IZI8; Pseudomonas sp. URMO17WK12:I6

**Figure S1**

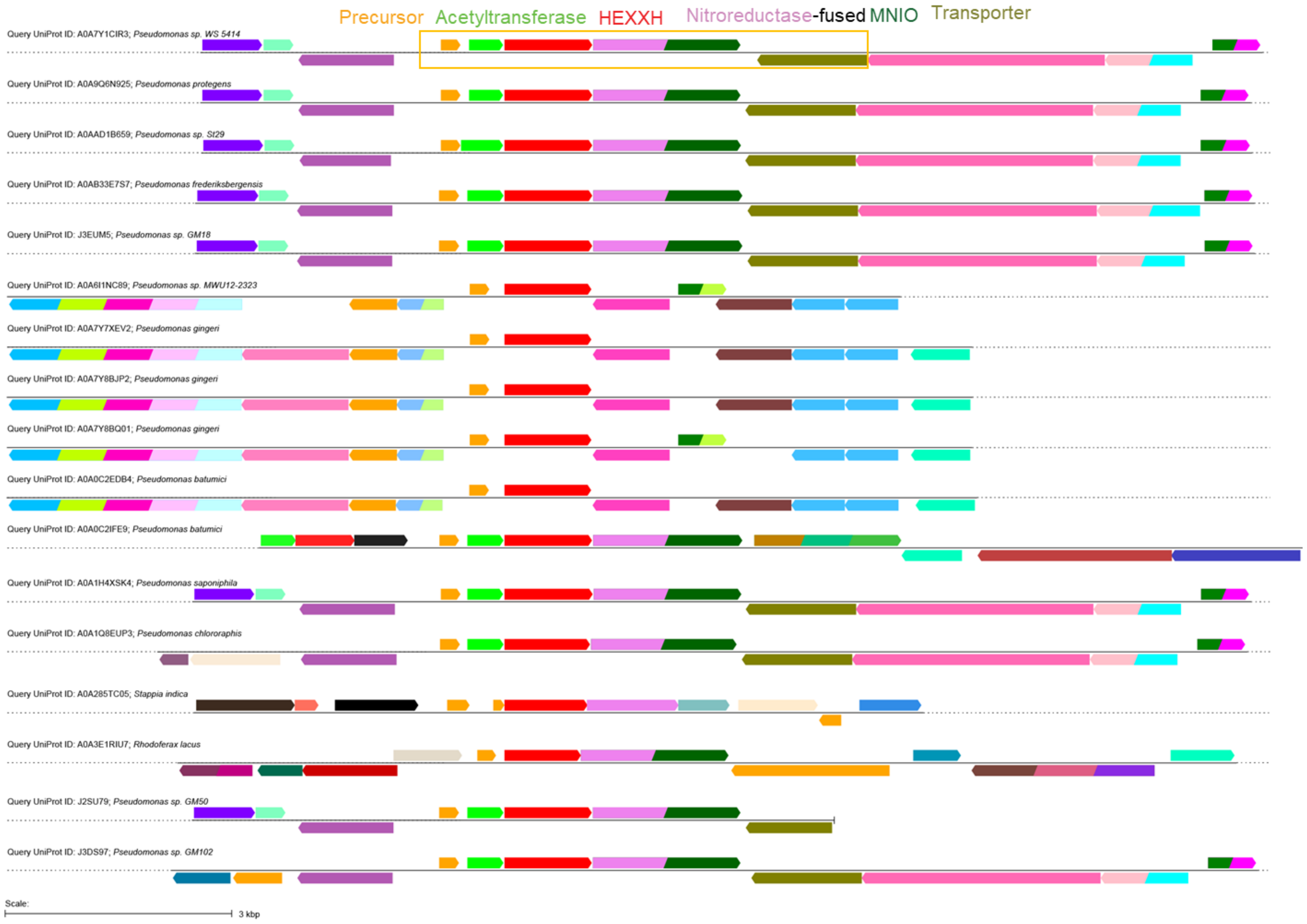

Precursor
Acetyltransferase
HEXXH
Nitroreductase-fused MNIO
Transporter
Query UniProt ID: A0A7Y1CIR3; *Pseudomonas sp. WS 5414*
Query UniProt ID: A0A9Q6N925; *Pseudomonas protegens*
Query UniProt ID: A0AAD1B659; *Pseudomonas sp. St29*
Query UniProt ID: A0AB33E7S7; *Pseudomonas frederiksbergensis*
Query UniProt ID: J3EUM5; *Pseudomonas sp. GM18*
Query UniProt ID: A0A6I1NC89; *Pseudomonas sp. MWU12-2323*
Query UniProt ID: A0A7Y7XEV2; *Pseudomonas gingeri*
Query UniProt ID: A0A7Y8BJP2; *Pseudomonas gingeri*
Query UniProt ID: A0A7Y8BQ01; *Pseudomonas gingeri*
Query UniProt ID: A0A0C2EDB4; *Pseudomonas batumici*
Query UniProt ID: A0A0C2IFE9; *Pseudomonas batumici*
Query UniProt ID: A0A1H4XSK4; *Pseudomonas saponiphila*
Query UniProt ID: A0A1Q8EUP3; *Pseudomonas chlororaphis*
Query UniProt ID: A0A285TC05; *Stappia indica*
Query UniProt ID: A0A3E1RIU7; *Rhodoferax lacus*
Query UniProt ID: J2SU79; *Pseudomonas sp. GM50*
Query UniProt ID: J3DS97; *Pseudomonas sp. GM102*
Scale:
3 kbp

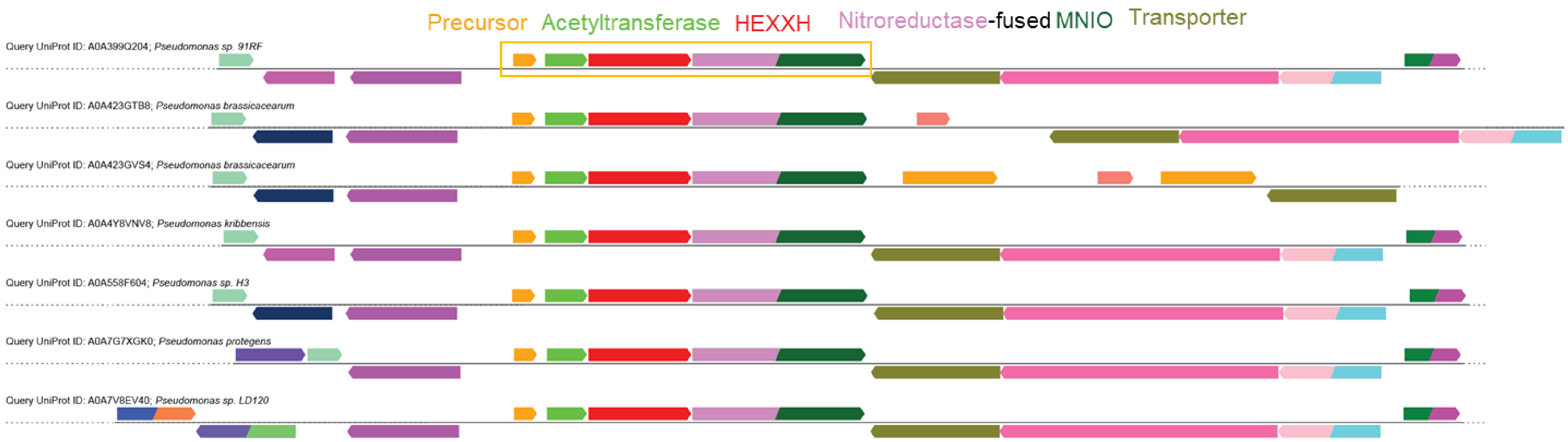


**Figure S1.** Biosynthetic clusters that are homologous to the *pfl* and *pos* BGCs are prevalent in *Pseudomonas*. The nitroreductase-MNIO fusion protein and the acetyltransferase are not always conserved. The Uniprot number A0A2T6GNH5 *in Pseudomonas protegens* corresponds to PosC from strain *Pseudomonas* sp. Os17.

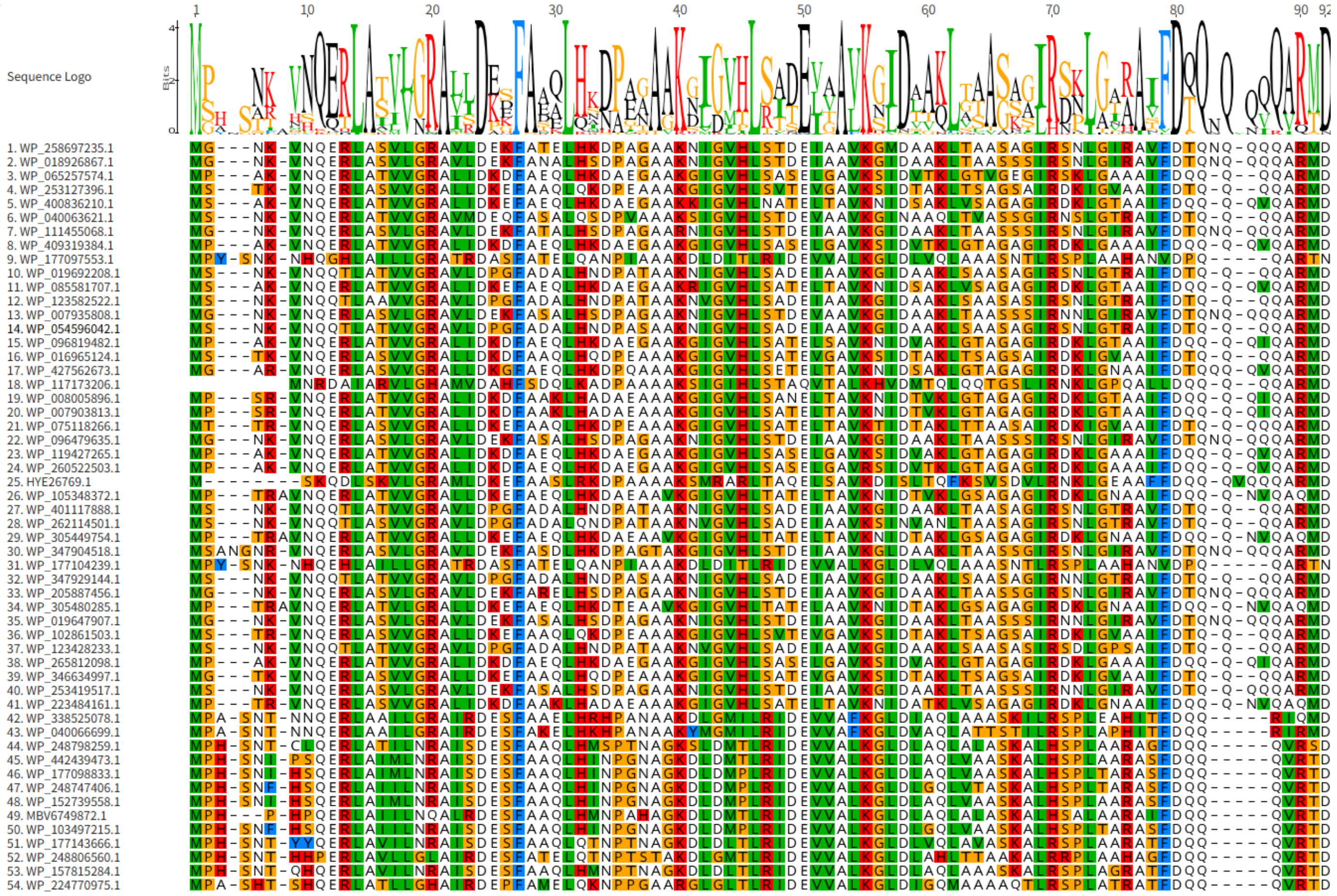


**Figure S2.** Sequences used to generate the logo in Figure 2B. Sequences were obtained by using PosA as the query sequence in a PHI-Blastp.

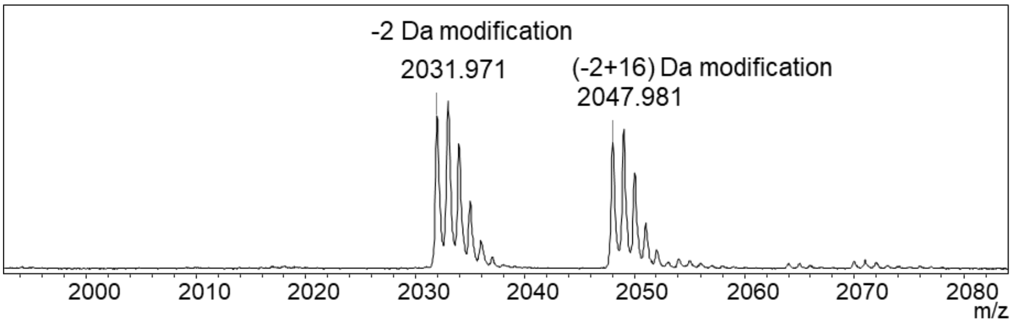


**Figure S3.** MALDI-TOF mass spectrum of LysC-digested PflD-modified PflA (calculated m/z 2031.993, 2047.988).

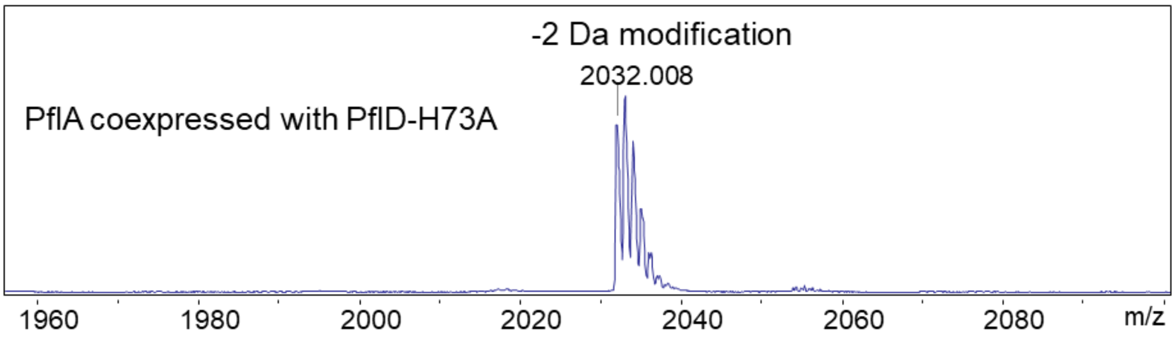


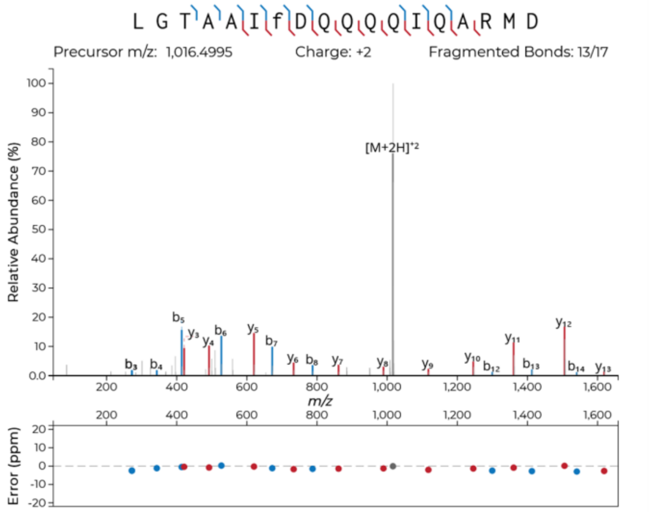


**Figure S4**. (Top) MALDI-TOF mass spectrum of LysC-digested PflD-H73A-modified PflA. (Bottom) qTOF-MS/MS spectrum (calculated m/z for PflA minus two hydrogens: 2031.9928). Fragment ion annotation and ppm errors were generated using the interactive peptide spectral annotator[4] with residues indicated in lower case f entered as dehydroPhe (-2 Da).

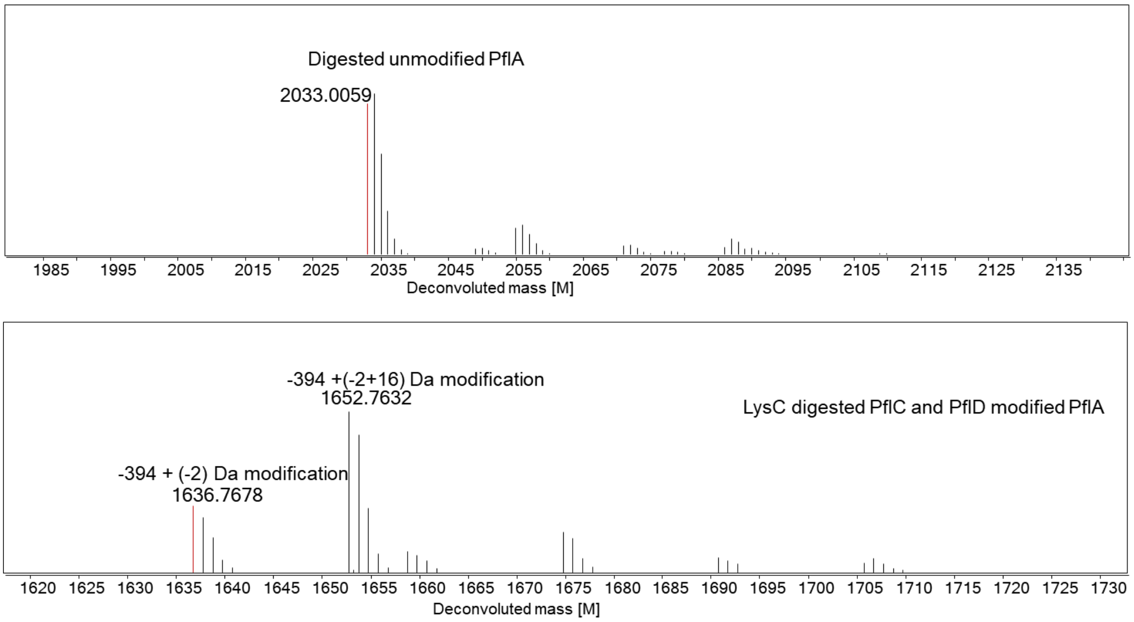


**Figure S5.** LC-HRMS spectra (ESI) of one of the fragments of LysC-digested PflA co-expressed with PflCD (digested unmodified sequence: LGTAAIFDQQQQIQARMD, calculated monoisotopic [M] 2033.000024). (Top) LC-MS spectrum of LysC-digested unmodified PflA. (Bottom) LC-MS spectrum of LysC-digested PflA modified by PflC and PflD (calculated [M] 1652.7425, 1636.7475). The modification of −394 Da was calculated by the mass difference between the mono isotopic masses, which are highlighted in red.

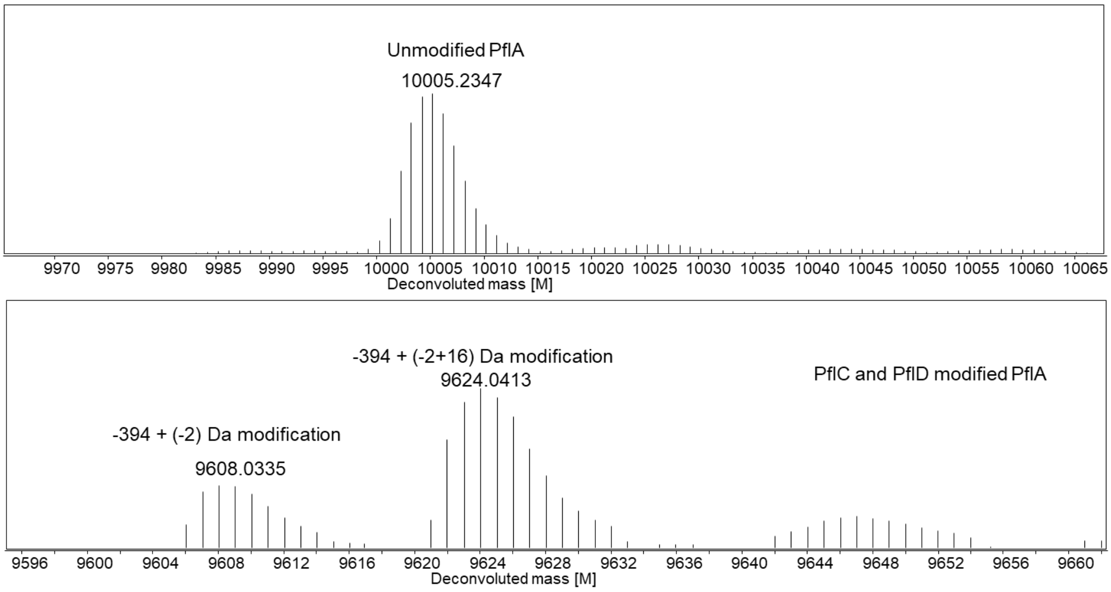


**Figure S6.** HR-LC-MS spectra of PflA and PflCD-modified PflA. (Top) LC-MS spectrum (ESI) of full length unmodified PflA (calculated [M] 10005.1055). (bottom) LC-MS spectrum of PflCD-modified PflA (calculated [M] 9624.5067, 9608.5117). Average masses are shown. For monoisotopic masses, see Fig. S5.

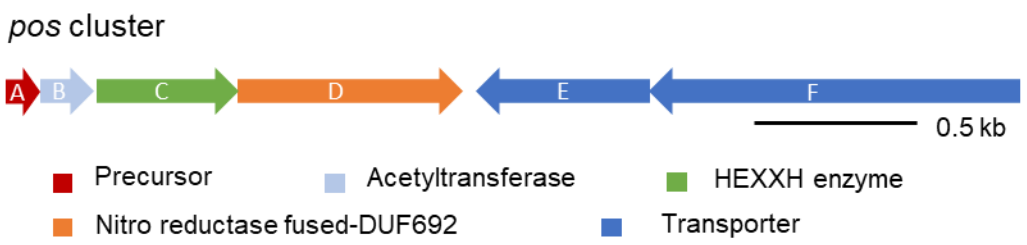


**PosA**:
MSTKVNQERLASVVGRALLDKDFAAQLHQDPEAAAKGIGVHLSATEV
GAVKSIDTAKLTSAGSAIRDK IGVAAIFDTQQQQARMD

**Figure S7.** *pos* BGC with precursor peptide sequence.

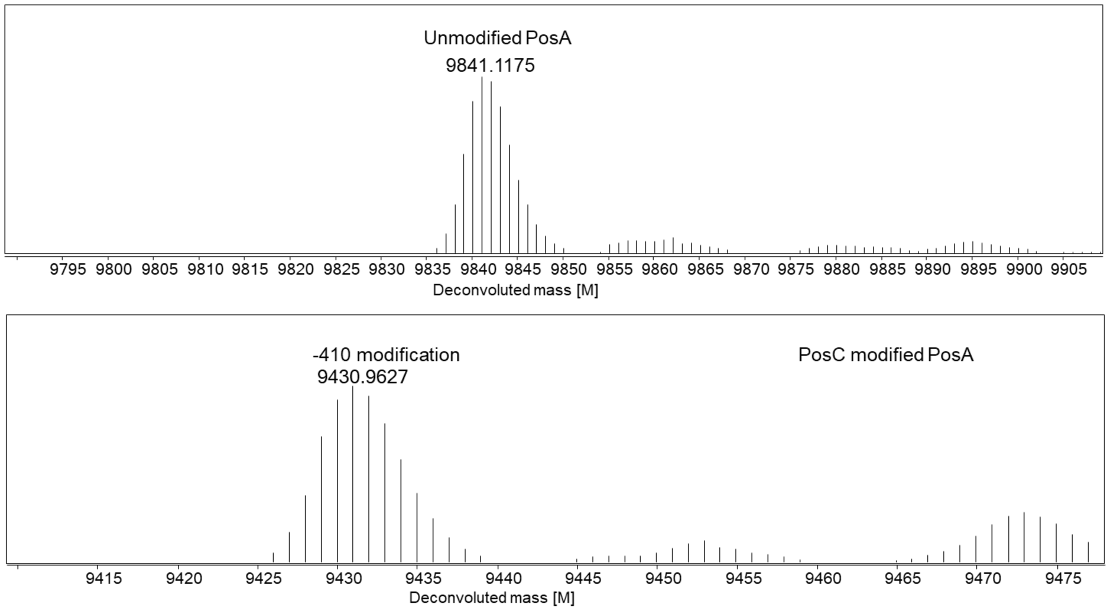


**Figure S8.** LC-HRMS spectra of PosA and PosC-modified PosA. (Top) LC-MS spectrum of full length unmodified PosA (calculated [M] 9841.8428). (Bottom) LC-MS spectrum of PosC-modified PosA (calculated [M] 9431.2685). Average mass shown.

**A**

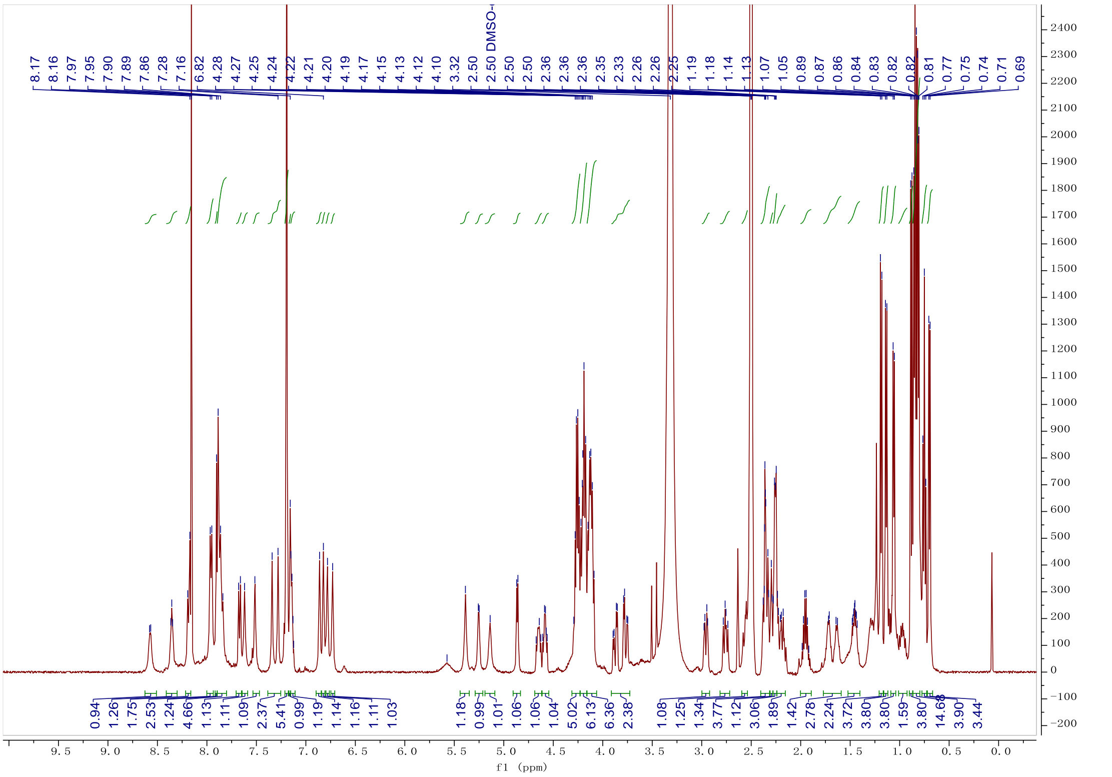

**B**

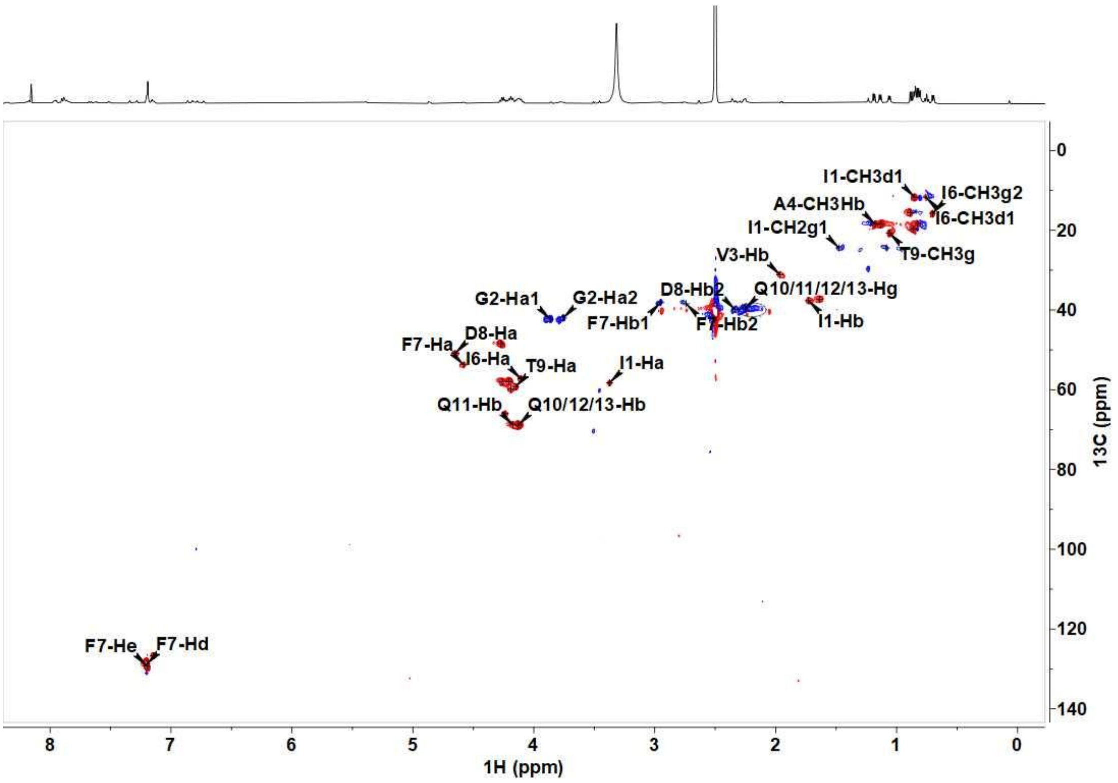
I1-CH3d1
I6-CH3g2
A4-CH3Hb
I6-CH3d1
I1-CH2g1
T9-CH3g
V3-Hb
D8-Hb2
Q10/11/12/13-Hg
G2-Ha1
G2-Ha2
F7-Hb1
F7-Hb2
I1-Hb
F7-Ha
D8-Ha
I6-Ha
T9-Ha
I1-Ha
Q11-Hb
Q10/12/13-Hb
F7-He
F7-Hd
13C (ppm)
1H (ppm)

**C**

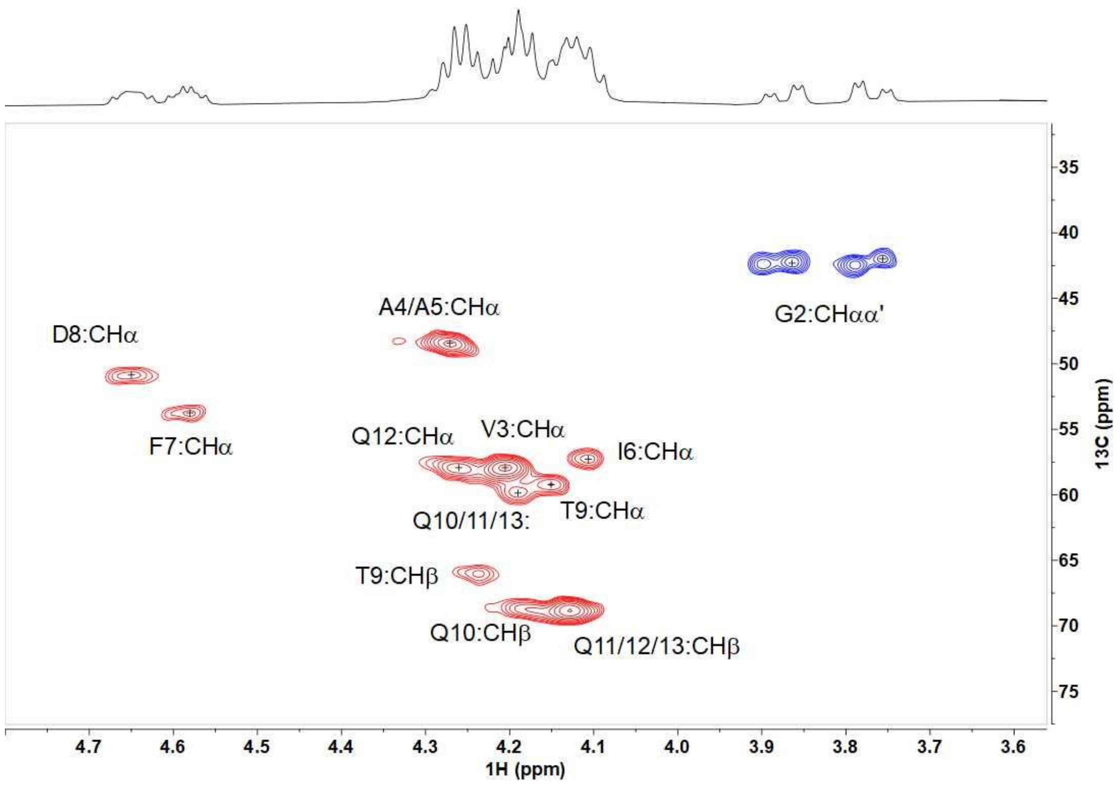


**Figure S9.** (A) The $^1H$ NMR spectrum of PosC-modified PosA. (B) 2D $^1H$-$^{13}C$ HSQC spectrum. The $^1H$ and $^{13}C$ cross peaks are labeled in the figure, and (C) 2D $^1H$-$^{13}C$ HSQC spectrum: ~ 4 ppm region. The $^1H$ and $^{13}C$ cross peaks are labeled in the figure. Hydroxylation occurred at the β-position of each of the four Gln residues, resulting in $^{13}C$ chemical shifts of the βCH groups near 69 ppm.

A

B

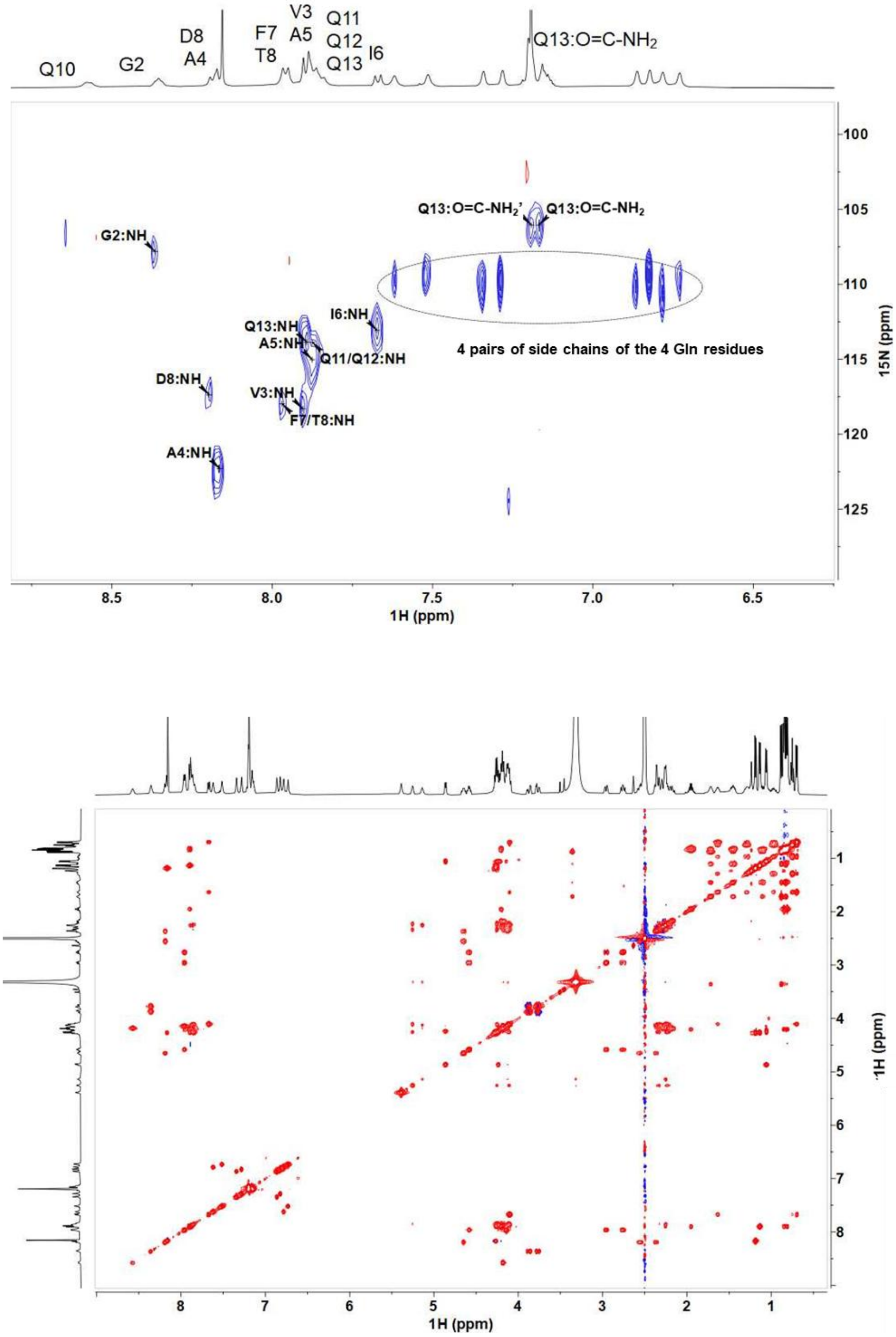


**Figure S10.** (A) 2D $^1$H and $^{15}$N HSQC spectrum of PosC-modified PosA. All $^1$H-$^{15}$N cross peaks are observed except the NH of Gln 10 residue. In the elipse are shown the side chain amide protons of the four Gln residues (8 protons). (B) The 2D $^1$H-$^1$H TOCSY spectrum in DMSO-d6 of PosC-modified PosA.

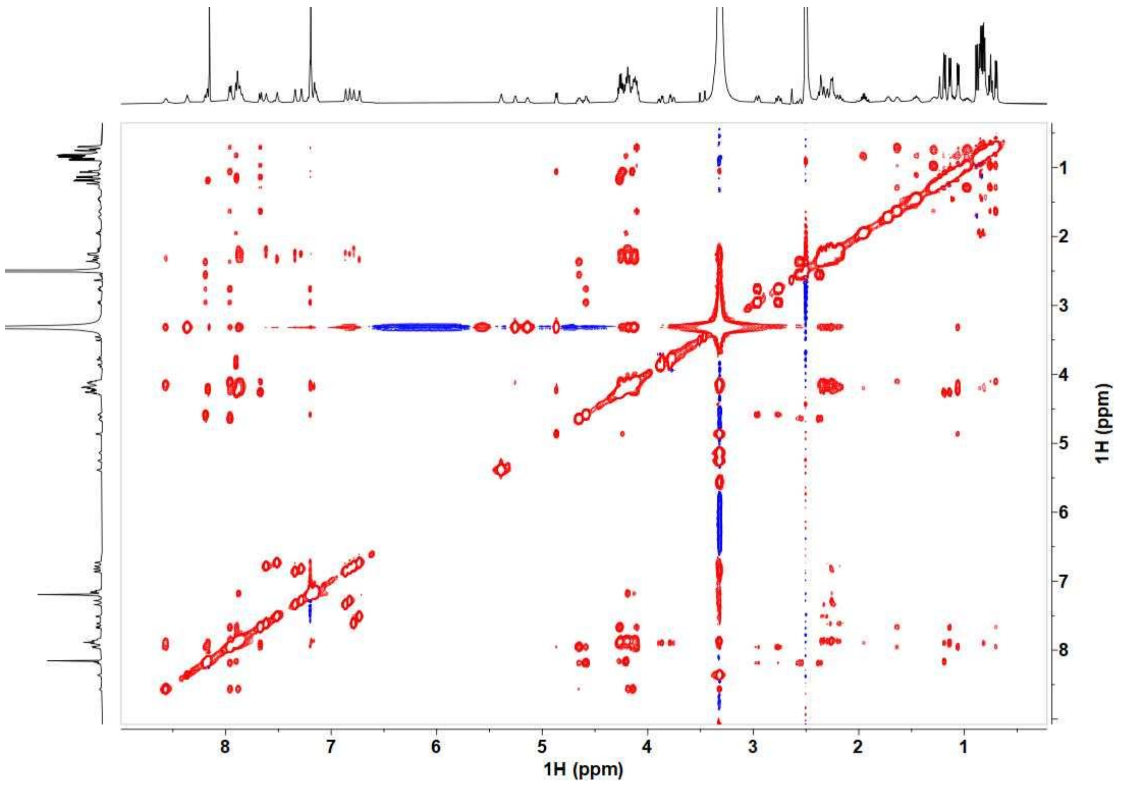


**Figure S11.** 2D $^1H$-$^1H$ NOESY spectrum in DMSO-d6 of PosC-modified PosA.

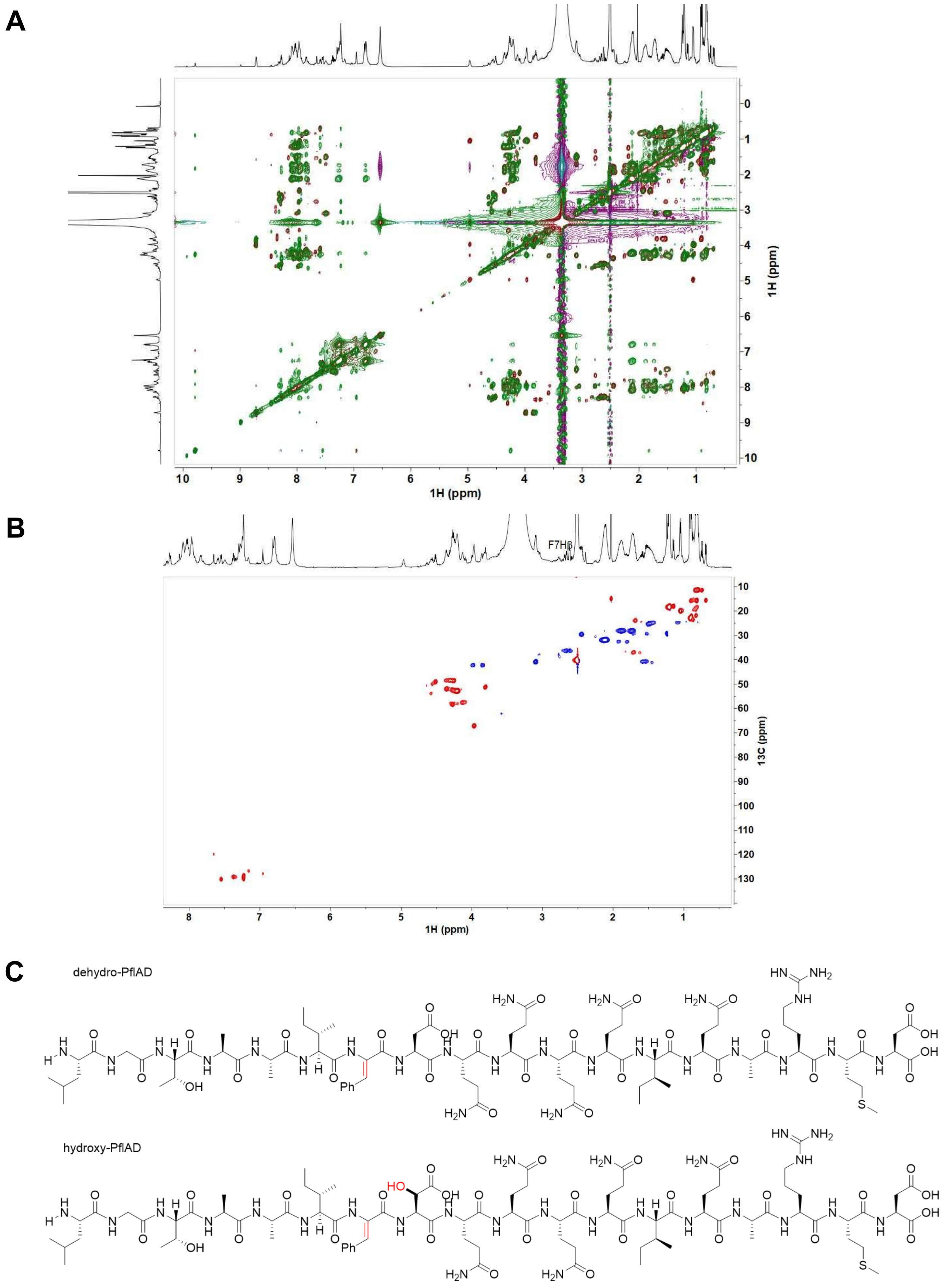


**Figure S12.** PflD-modified PflA digested with LysC generated a mixture of peptide **1** (PflAD1; isolated and characterized by NMR), peptide **2** (PflAD2; not isolated due to the low concentration), and unmodified PflA (~40% of the isolated peptide quantified by qNMR). (A) The TOCSY-NOESY spectra of **1** are overlayed, with the TOCSY spectrum in purple, and the NOESY spectrum in green. (B) 2D $^{1}H$-$^{13}C$ HSQC spectrum of **1**. The CH/$CH_3$ groups are in red and $CH_2$ groups are in blue. (C) The NMR experiments on dehydro-PflAD showed the dehydroPhe has the *Z*-configuration.

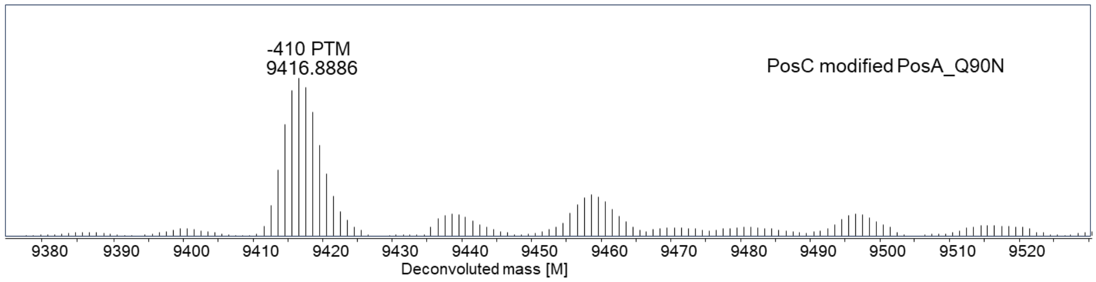


**Figure S13.** HR-LC-ESI mass spectrum of PosA-Q90N co-expressed with PosC (calculated [M] 9417.2419). Average mass shown.

PosA-Q90N sequence:

MGSSHHHHHHSTKVNQERLASVVGRALLDKDFAAQLHQDPEAAAKGIGVHLSATEVGAVKSIDTAK LTSAGSAIRDKIGVAAIFDTQQQ**N**ARMD.

PosC product:

MGSSHHHHHHSTKVNQERLASVVGRALLDKDFAAQLHQDPEAAAKGIGVHLSATEVGAVKSIDTAK LTSAGSAIRDKIGVAAIFDT-hQ-hQ-hQ-hN; hQ = 3-hydroxyGln; hN = 3-hydroxyAsn with amido terminus.

**A**

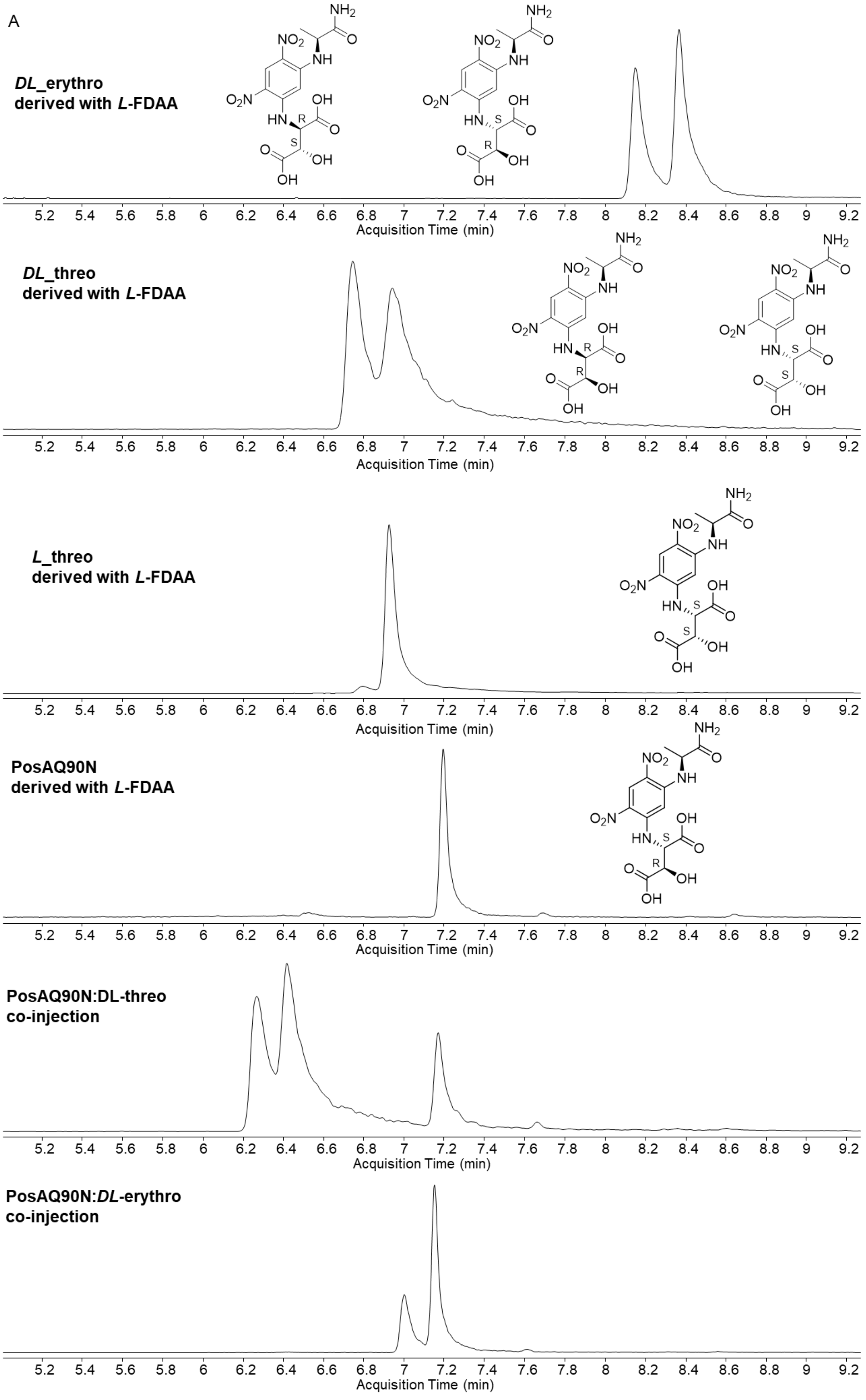
A
DL_erythro
derived with L-FDAA
DL_threo
derived with L-FDAA
L_threo
derived with L-FDAA
PosAQ90N
derived with L-FDAA
PosAQ90N:DL-threo
co-injection
PosAQ90N:DL-erythro
co-injection
Acquisition Time (min)
5.2 5.4 5.6 5.8 6 6.2 6.4 6.6 6.8 7 7.2 7.4 7.6 7.8 8 8.2 8.4 8.6 8.8 9 9.2
NH2
NO2
NH
O2N
HN
OH
O
R
S

**B**

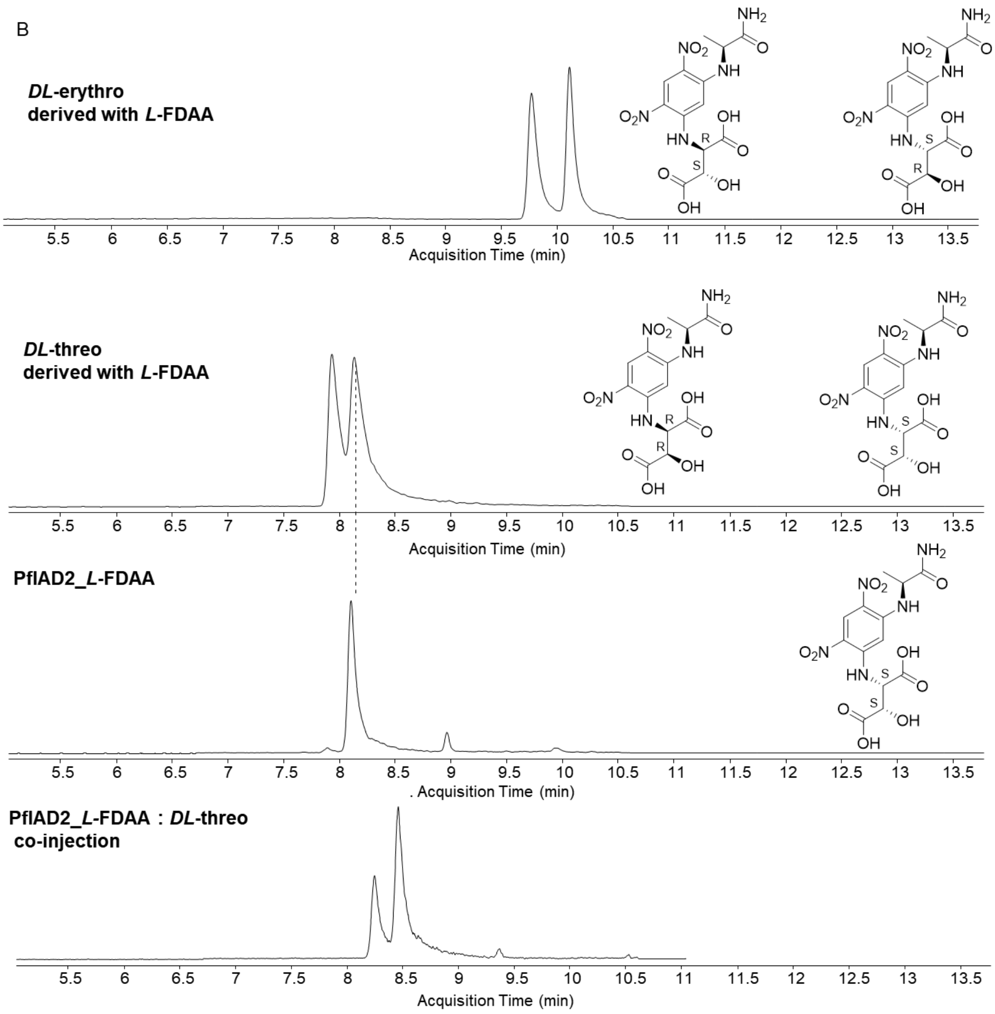


**Figure S14.** Marfey's analysis of (A) PosA-Q90N modified by PosC. Hydrolyzed PosA-Q90N and the standard samples were derivatized with *L*-FDAA. Note that the 3-hydroxy-Asn is hydrolyzed to 3-hydroxy-Asp under standard Marfey's analysis conditions. Because of retention time drift, the identity of the product from PosA-Q90N was determined by co-injections with derivatized *DL-threo* and *DL-erythro*-standards. The PosA-Q90N product matches one of the isomers in the DL-*erythro* 3-hydroxy-Asp standard, which is composed of two enantiomers with a configuration of (2*S*, 3*R*) and (2*R*, 3*S*). Because the ribosomally derived Asn has the *S* configuration at C2*,* the 3-hydroxy-Asn product was determined to have the (2*S*,3*R*) configuration. (B) Marfey's analysis of PflD-modified PflA that underwent a +14 Da change (hydroxy-PflAD). Hydrolyzed hydroxy-PflAD and the standards were derivatized with *L*-FDAA.. The PflD product matches one of the isomers in the DL-*threo* 3-hydroxy-Asp standard, which is composed of two enantiomers with a configuration of (2*S*, 3*S*) and (2*R*, 3*R*). Because the ribosomally derived Asp has the *S* configuration at C2*,* the 3-hydroxy-Asp product of PflD was determined to have the (2*S*,3*S*) configuration.

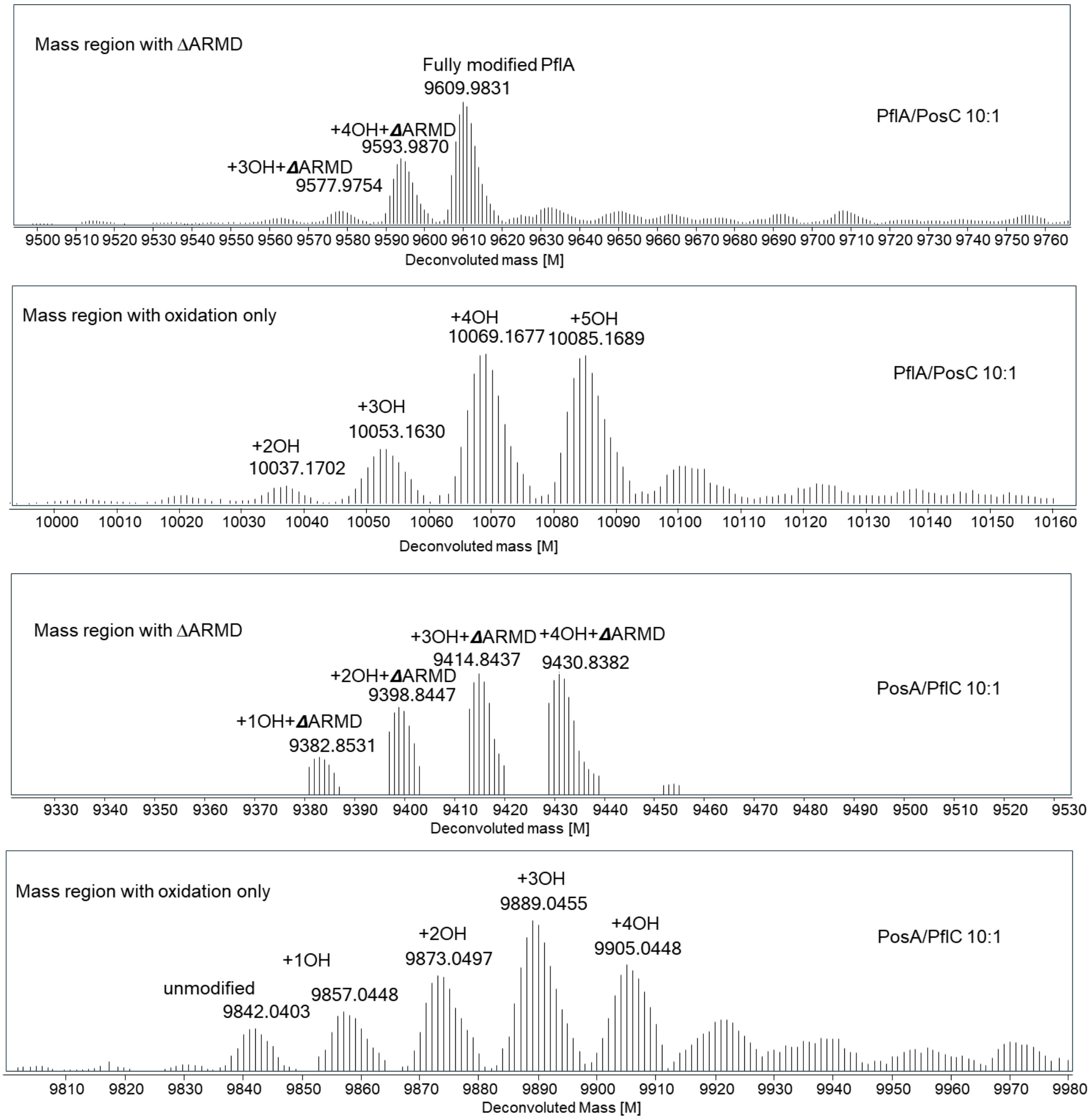


**Figure S15**. In vitro cross activity of HEXXH enzymes. PosC reacted with PflA (top) and PflC reacted with PosA (bottom) at 1:10 ratio (enzyme to peptide) under standard conditions. PosC-modified PflA, calculated [full modification] 9610.5262, [+4OH+∆ARMD] 9594.5312. [+3OH+∆ARMD] 9578.536164. [+5OH] 10085.0805, [+4OH] 10069.0855, [+3OH] 10053.0905, [+2OH] 10037.0955. PflC-modified PosA, calculated [+4OH+∆ARMD] 9431.2685, [+3OH+∆ARMD] 9415.2735, [+2OH+∆ARMD] 9399.2785, [+1OH+∆ARMD] 9383.2835. Unmodified PosA calculated [M] 9841.8428, [+4OH] 9905.8228, [+3OH], 9889.8278, [+2OH] 9873.8328, [+1OH] 9857.8378. Average mass values are shown.

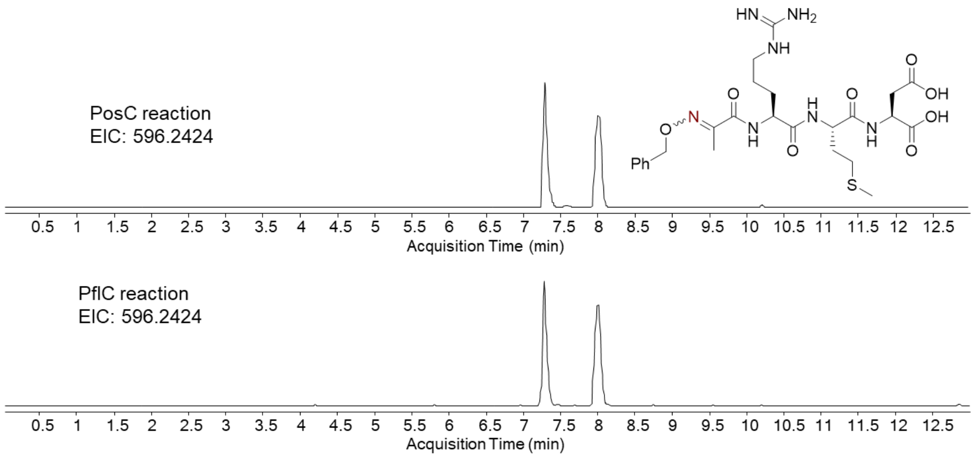


**Figure S16**. HR-LC-MS extracted ion chromatograms (EICs) of the products of the ARMD motif after PflC-catalyzed backbone cleavage. *O*-Benzylhydroxylamine was added to the in vitro reaction of PflC/PflA or PosC/PosA. Calculated m/z for the oxime products: 596.2497.

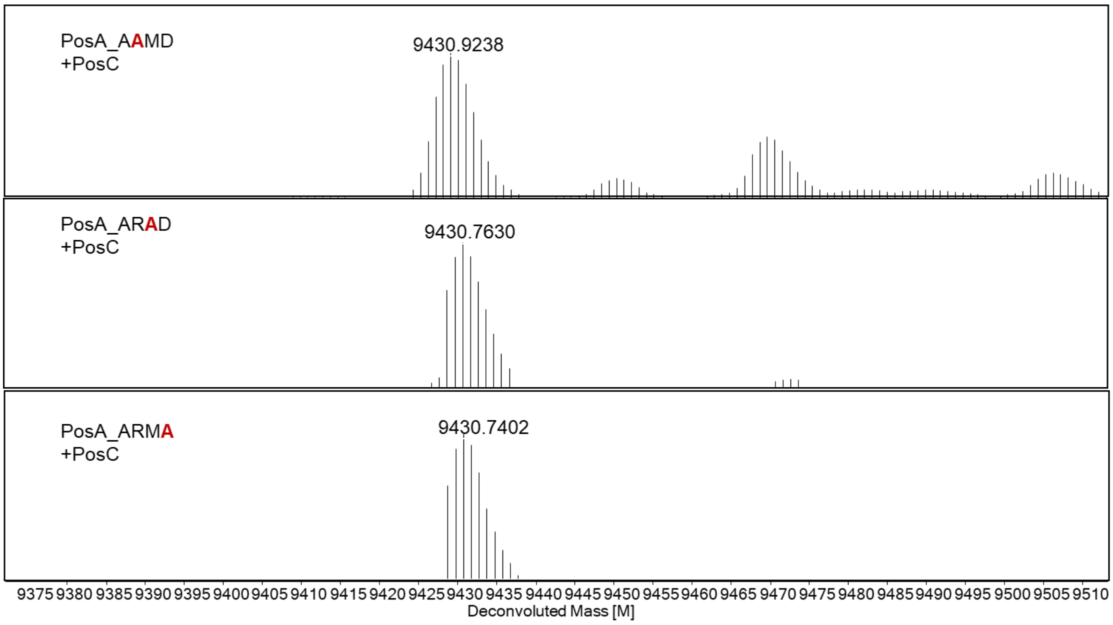


**Figure S17**. LC-HRMS spectra of PosA variants coexpressed with PosC. Alanine scan across the ARMD motif afforded the same modifications as observed with WT PosA with four hydroxylated Gln residues and removal of the ARMD sequence (calculated [M] 9431.2685). Average mass is shown.

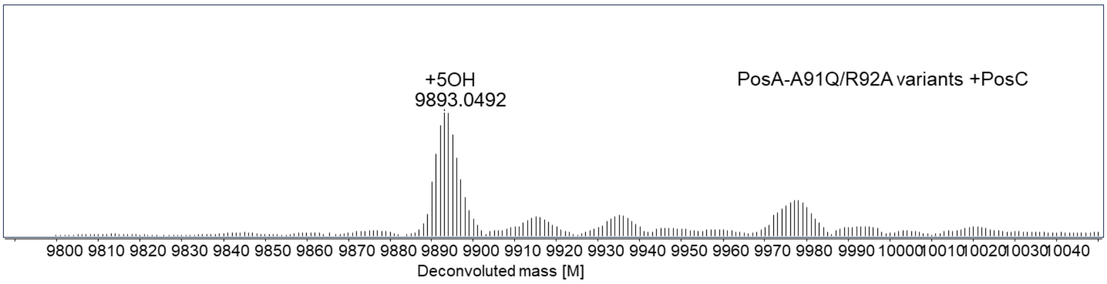


**Figure S18**. LC-HRMS spectrum of a PosA-A91Q/R92A variant coexpressed with PosC (calculated [M] 9893.7613). Average mass is shown. No backbone cleavage was observed.

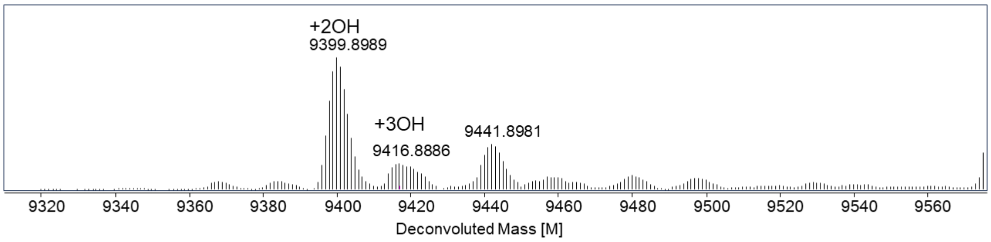


**Figure S19**. HR-LC-MS spectrum of PosA-ΔARMD mutant coexpressed with PosC, calculated [M] 9400.2858). Average mass is shown.

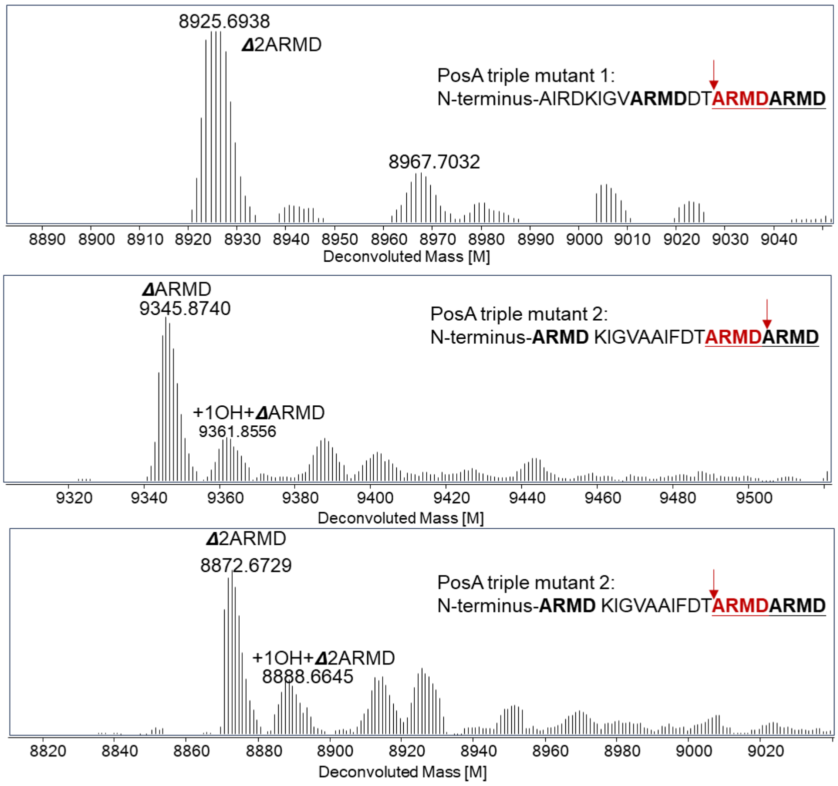


**Figure S20**. LC-HRMS spectra of PosA-triple ARMD variants coexpressed with PosC. Triple mutant 1 gave one major product as double cleavage of ARMD (calculated [M] 8925.8313), while mutant 2 gave two products, the major product was formed by mono-ARMD removal (calculated [M] 9346.3571) and the minor product was double ARMD cleavage (calculated [M] 8872.8100). Average mass shown.

Sequence of mutant sequences containing three ARMD motifs ("triple mutants") in comparison with WT (mutations in red font):

WT PosA: MGSSHHHHHHSTKVNQERLASVVGRALLDKDFAAQLHQDPEAAAKGIGVHLSATEVGAVKSIDTAK LTSAGSAIRDKIGVAAIFDTQQQQARMD

Triple mutant 1: MGSSHHHHHHSTKVNQERLASVVGRALLDKDFAAQLHQDPEAAAKGIGVHLSATEVGAVKSIDTAK LTSAGSAIRDKIGVARMDDTARMDARMD

Triple mutant 2: MGSSHHHHHHSTKVNQERLASVVGRALLDKDFAAQLHQDPEAAAKGIGVHLSATEVGAVKSIDTAK LTSAGSARMDKIGVAAIFDTARMDARMD

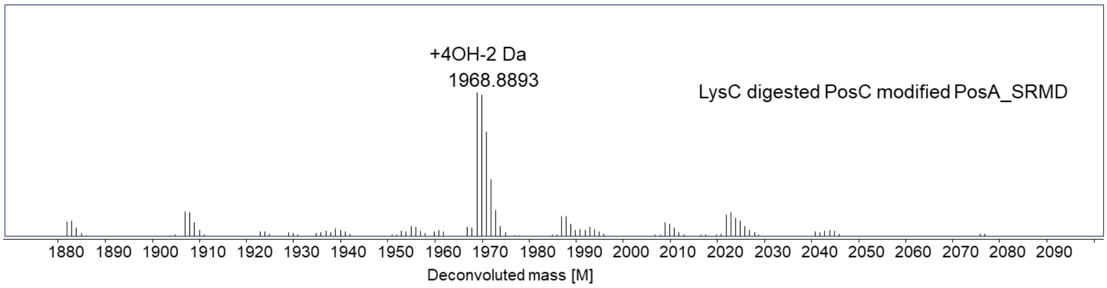


**Figure S21**. HR-LC-MS spectrum of LysC-digested PosC-modified PosA-SRMD mutant (digested sequence: IGVAAIFDTQQQQSRMD, calculated [M]1968.8862). Monoisotopic mass is shown.

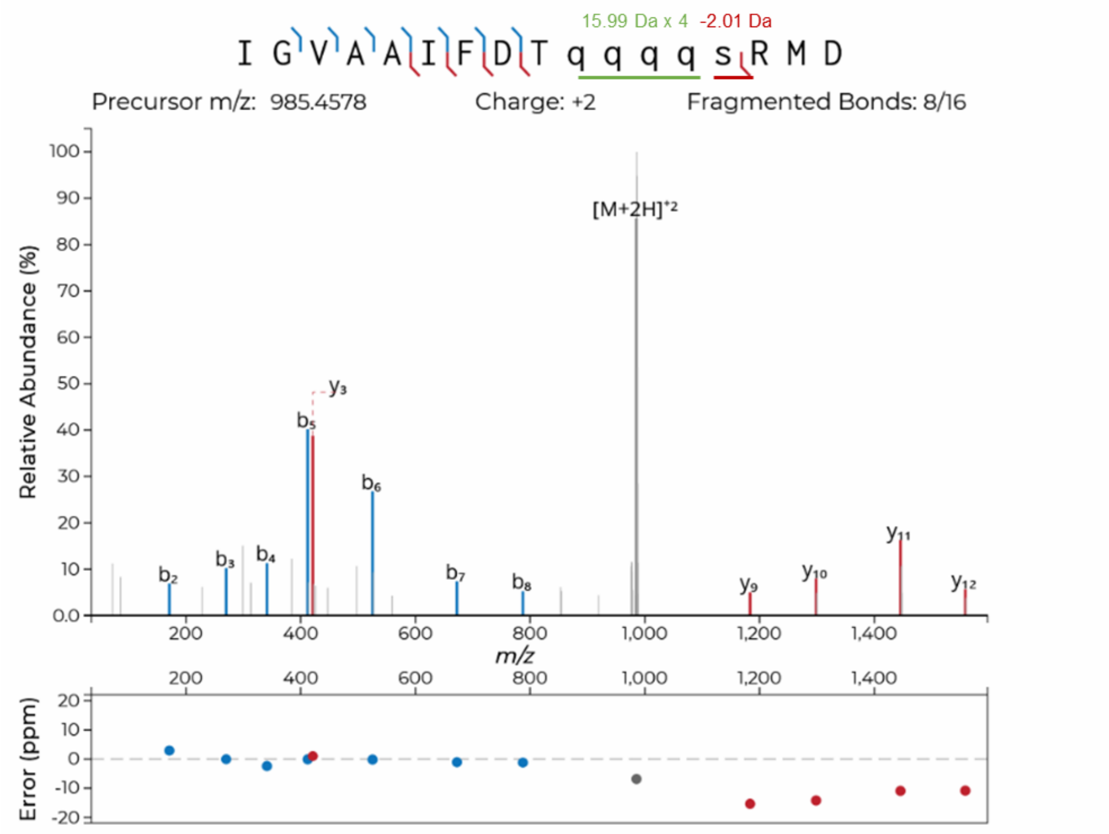


**Figure S22**. HR-MS/MS spectrum of LysC-digested PosC-modified PosA-SRMD (digested sequence: IGVAAIFDTQQQQSRMD). Fragment ion annotation and ppm errors were generated using the interactive peptide spectral annotator.[4]

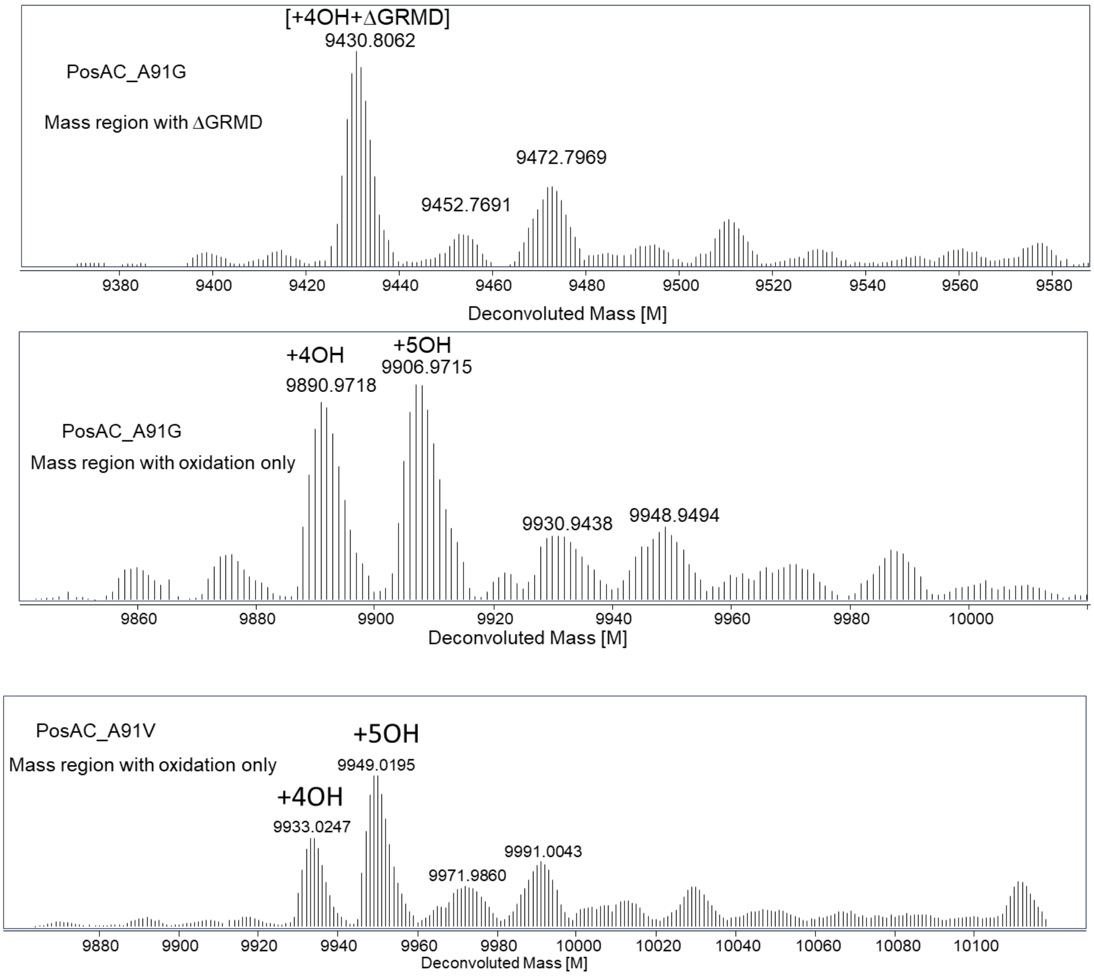


**Figure S23**. LC-HRMS spectra of PosA-A91G and A91V coexpressed with PosC in *E. coli*. (Top) PosC-modified PosA-A91G shows a major product in which four Gln residues are hydroxylated and the GRMD sequence is removed. (Middle) PosC-modified PosA-A91G also results in products of four and five hydroxylations in which the GRMD sequence is not removed. (Bottom) PosC-modified PosA-A91V was hydroxylated four and five times but no cleavage and removal of the VRMD sequence was observed. Calculated mass for PosAC_GRMD, [+4OH+ΔGRMD] 9431.2685, [+4OH] 9891.7962, [+5OH] 9907.7912. Calculated mass for PosAC_VRMD, [+4OH] 9933.8760, [+5OH] 9949.8710. Average mass values are shown.

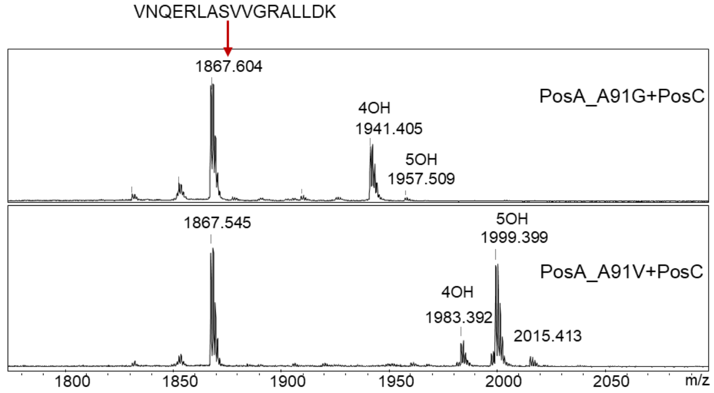


**Figure S24.** MALDI-ToF spectra of LysC digested PosA-A91G and A91V coexpressed with PosC. Calculated mass for PosAC_GRMD, [4OH] 1941.8974, [5OH] 1957.8924. Calculated mass for PosAC_VRMD, [4OH] 1983.9444, [5OH] 1999.9394. Monoisotopic mass values are shown. A different LysC-digestion product (residues 5-21 of PsA) is also observed as indicated.

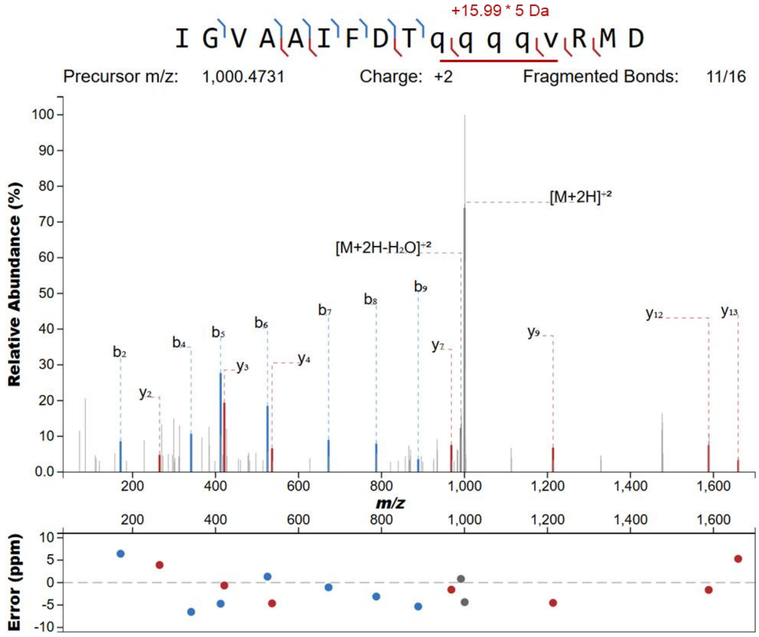


**Figure S25.** MS/MS spectrum of LysC-digested PosA-A91V coexpressed with PosC. We were unable to obtain a MS/MS spectrum for the five-fold hydroxylated LysC-digested A91G mutant.

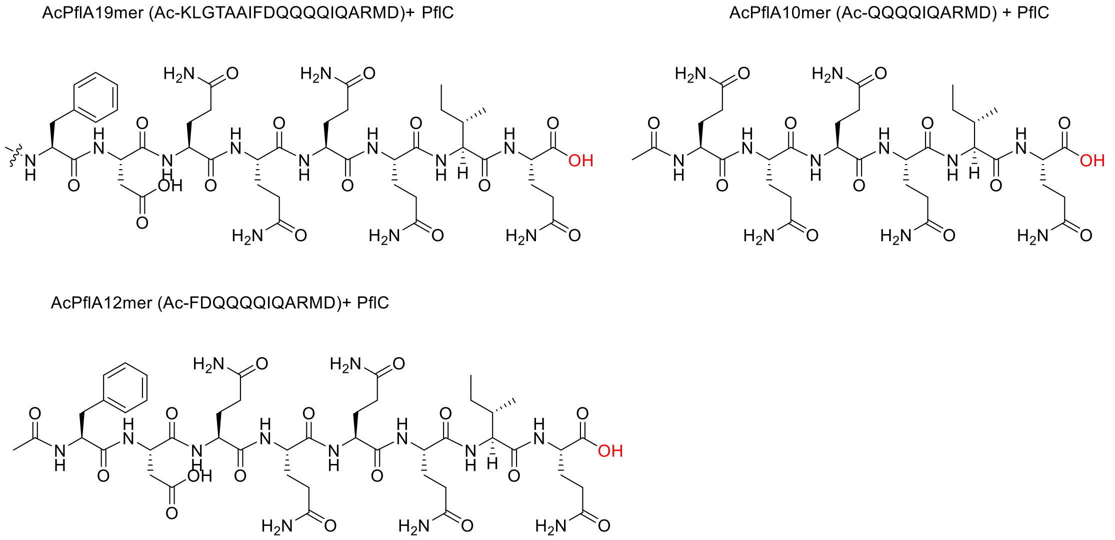


**Figure S26**. Hydrolytic products formed by incubation of PflC with the truncated peptides (shown above the products).

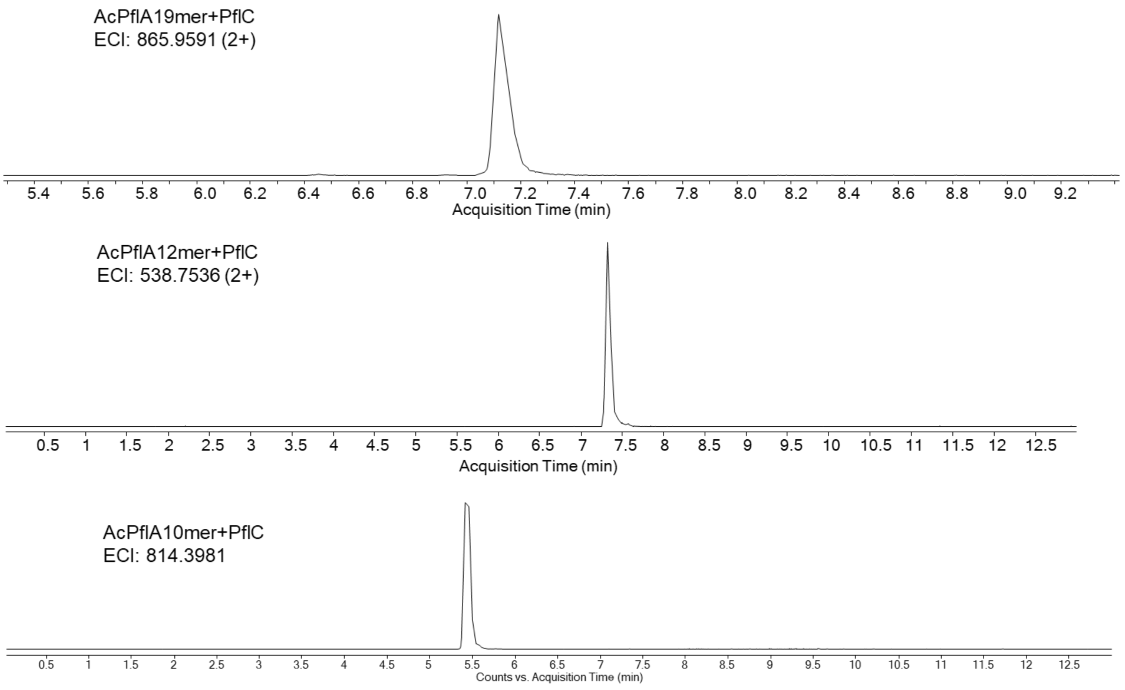


**Figure S27**. EICs of PflC-modified truncated peptides; extracted masses used are shown.

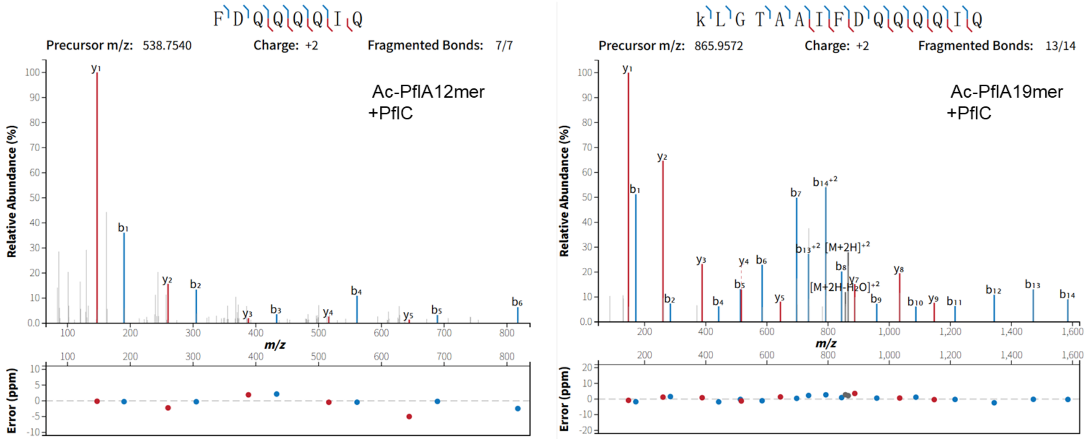


**Figure S28**. HR-MS/MS spectra of truncated synthetic peptides modified by PflC in vitro. The fragmentation data show a C-terminal carboxylic acid.

**A**

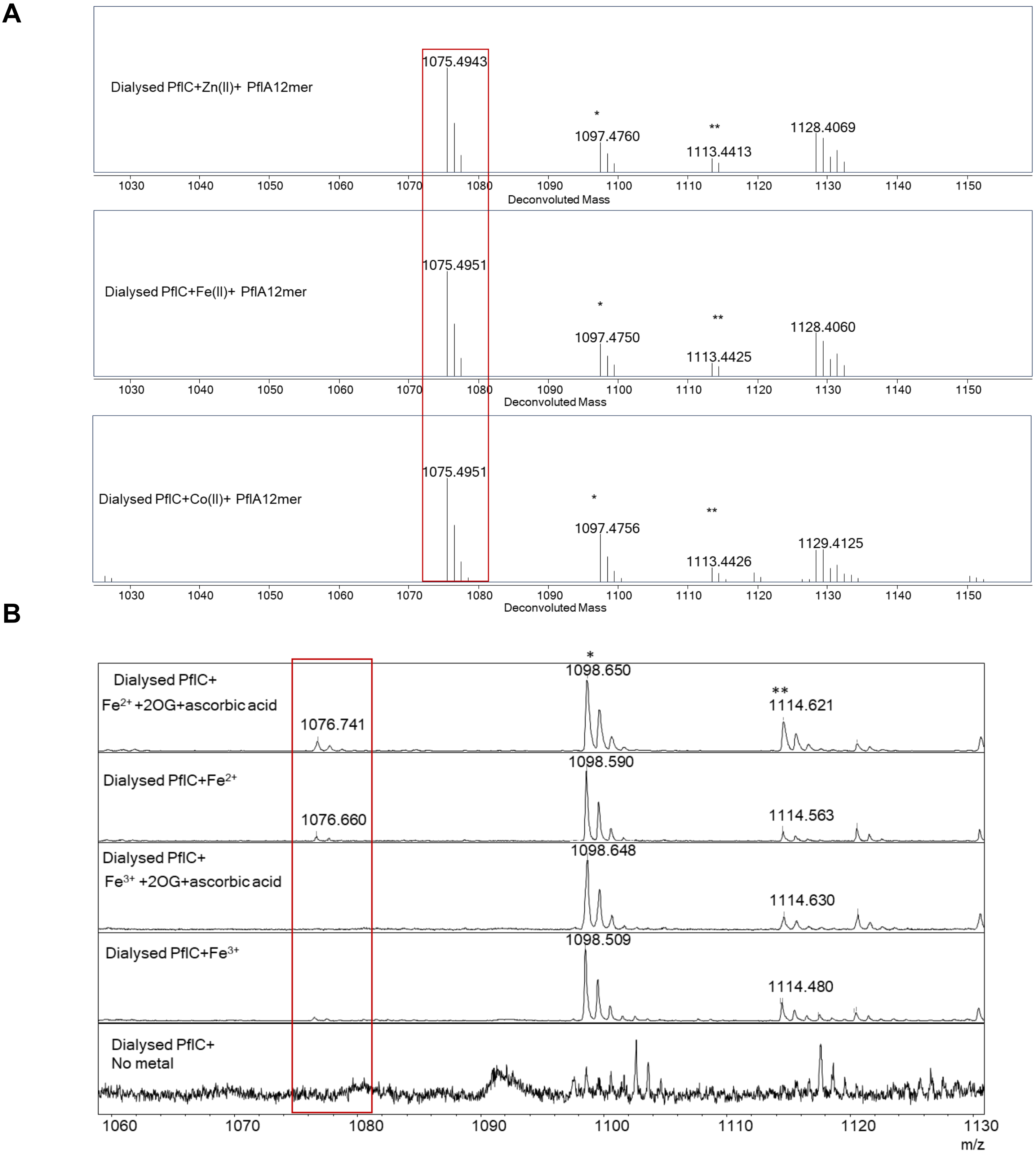


**Figure S29**. Observed hydrolytic activity with different reconstituted divalent metal ions. **A**) qToF-LC-MS data, shown as deconvoluted mass. Calculated monoisotopic mass 1075.4934. * $Na^+$ form of the ion; ** $K^+$ form of the ion. **B**) MALDI-ToF mass spectra showing $[M+H]^+$, $[M+Na]^+$, and $[M+K]^+$.

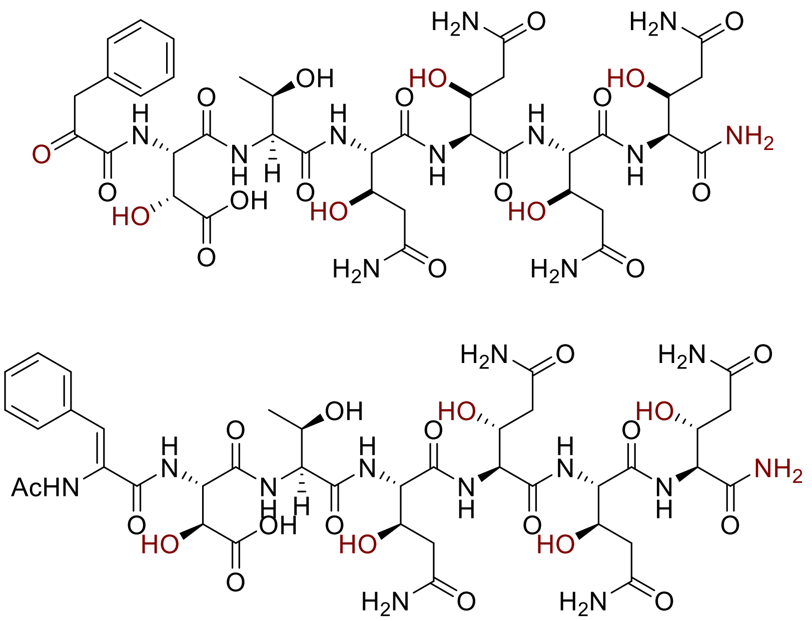

**Figure 30**. Potential final products of the *pos* BGC. If an aminopeptidase or protease were to remove the peptide sequence up to the ΔPhe, previous studies on N-terminal Dha/Dhb peptides would predict rapid hydrolysis of the resulting enamine to the ketone as shown in the top product. However, that would not explain the presence of the acetyltransferases PosB/PflB that are present in most, but not all, homologous BGCs (Figure S1). One possibility is that the ΔPhe after proteolysis is more stable than Dha/Dhb as a result of the additional resonance stabilization afforded by conjugation of the phenyl group, and that acetylation/acylation of the N-terminal amine is catalyzed by PosB/PflB (shown for acetylation in the bottom drawing). Another possibility is that PosB/PflB act after proteolysis, explaining their inactivity in co-expression experiments, and acylate/acetylate one of the hydroxyl groups similar to the recently reported acylation of the hydroxyl group of a lasso peptide.[6]

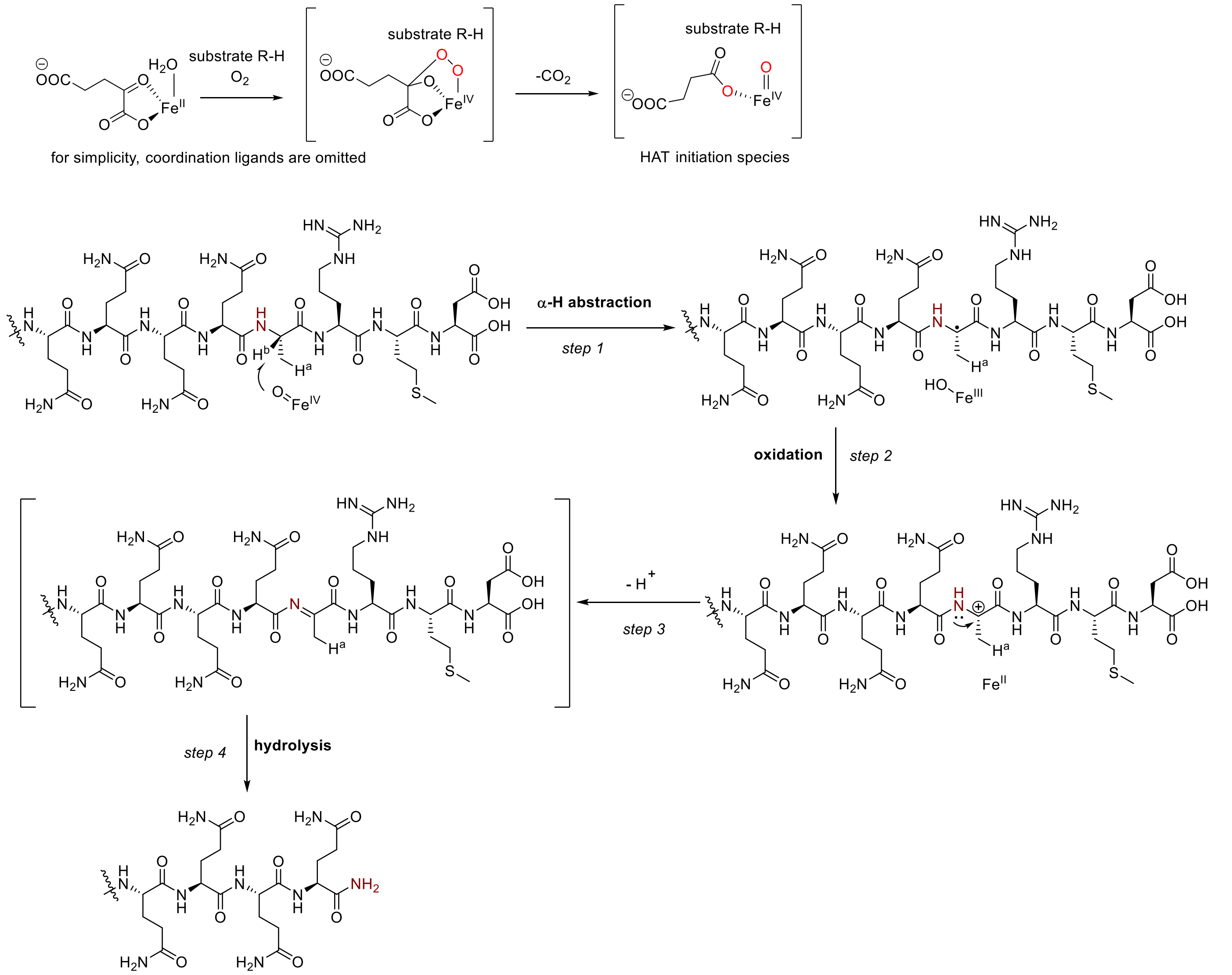


### Alternative step 2:

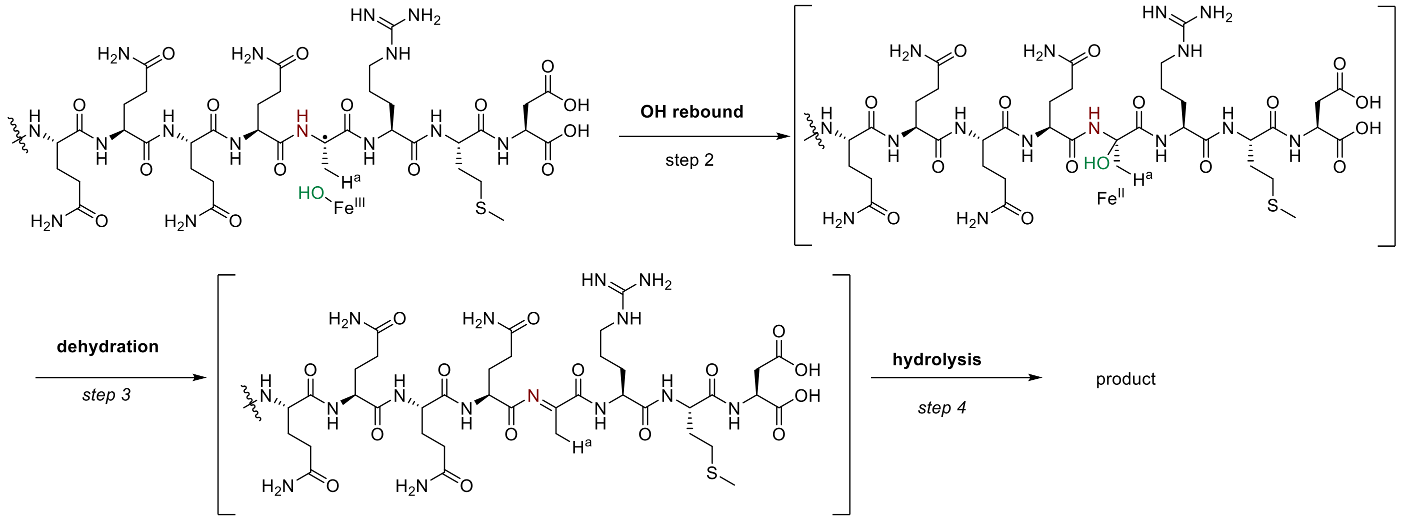


**Figure 31**. Mechanistic proposals for PflC-catalyzed hydroxylation and amide formation via α-hydrogen atom transfer.

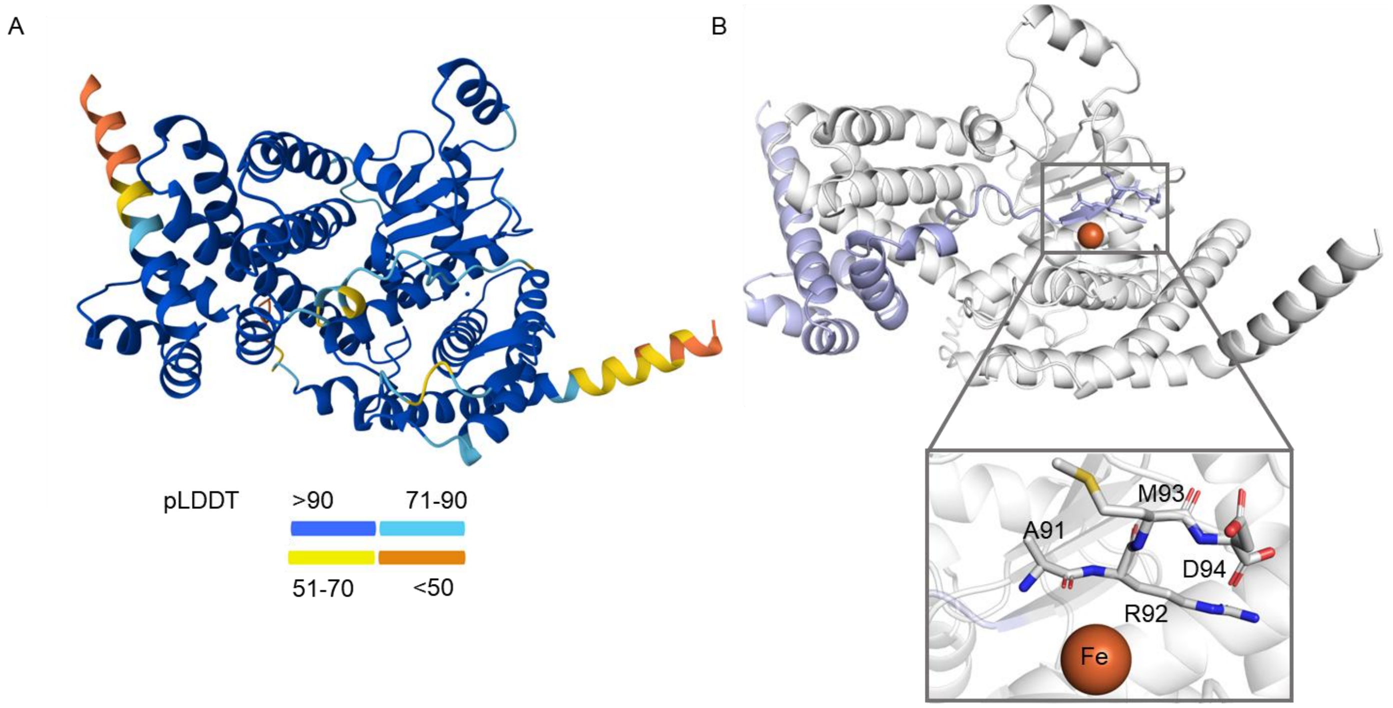


**Figure S32**. AlphaFold3[7] predicted model of PosC in complex with PosA. (A) pLDDT score-colored output. (B) Zoom-in image of the catalytic iron center and the C-terminal ARMD motif (shown in sticks, colored by elements) shows that the iron is right next to the cleavage site. Purple: PosA. Grey: PosC.

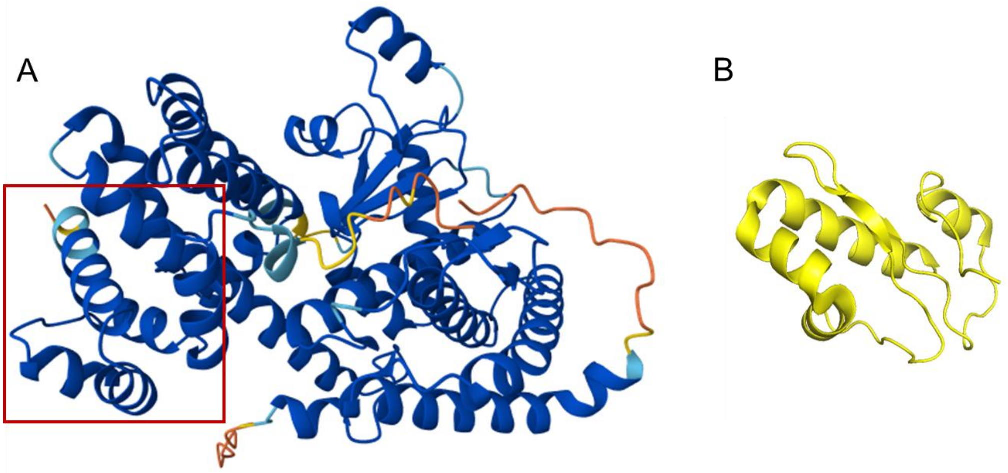


**Figure S33**. (A) AlphaFold3[7] model colored by pLDDT values for the complex of PflA with PflC. The red box shows the well-folded leader peptide region. DALI search confirmed that this N-terminal region is structurally like the NHLP solved by NMR. (B) NMR determined structure of an NHLP. PDB: 8TB1.[8]
